# Neurovascular Coupling in the Basolateral Amygdala Modulates Negative Emotions

**DOI:** 10.1101/2023.02.13.527949

**Authors:** Jiayu Ruan, Xinghua Quan, Huiqi Xie, Yiyi Zhang, Dongdong Zhang, Wentao Wang, Bingrui Zhao, Danping Lu, Yuxiao Niu, Mengyuan Diao, Wei Hu, Jie-Min Jia

## Abstract

Emotion induces changes in regional cerebral blood flow (CBF), a manifestation of neurovascular coupling (NVC), yet whether NVC might feed back to modulate emotion actively remains unexplored. Here, we demonstrate that NVC actively and bidirectionally modulates stress-induced negative emotions. We established bidirectional manipulations of NVC in freely moving mice by employing integrated pharmacological, genetic, and arteriolar optogenetic approaches. Our results showed that both systemic and basolateral amygdala (BLA) region-specific NVC deficiencies heightened emotional responses when mice transitioned from a safe, familiar environment to anxiogenic environments, and local restoration of NVC in the BLA normalized these responses. Mechanistically, NVC dysfunction impairs the capacity of BLA neuronal scaling during state transitions, manifesting as a characteristic biphasic pattern of c-Fos topology. Namely, the NVC-deficient animal aberrantly adopts high-stress configurations under mild stress but regresses to low-stress templates during high-demand survival threats, thereby compromising defensive sustainability. Notably, the genetic NVC-enhancement model counteracts NVC impairments caused by chronic stress, thereby alleviating stress-driven emotional distress. These findings established NVC in the BLA as an allostatic program that fine-tunes neural circuit activity for emotional responses, with implications for understanding and treating emotional disorders.

## INTRODUCTION

Emotion is a complex, integrative response to changes in internal and external environments.^1^ Emotional flexibility—the capacity to dynamically regulate and adapt these responses—is fundamental for mental health and survival.^2^ Conversely, deficits in this adaptive capacity result in emotional rigidity, a hallmark of dysregulation strongly associated with psychiatric disorders such as depression and post-traumatic stress disorder (PTSD).^3, 4^ Therefore, delineating the molecular and cellular mechanisms that govern the balance between emotional flexibility and rigidity is critical for elucidating the biological basis of resilience.^5, 6^

Neurovascular coupling (NVC), the core physiological foundation of blood oxygen level-dependent (BOLD) functional MRI (fMRI) signals, has long been categorized as a passive neural proxy.^7–10^ Clinical observations in the basolateral amygdala (BLA)—a pivotal hub for processing negative affect—reveal pervasive neurovascular dysregulation across diverse psychiatric landscapes.^11^ Specifically, exaggerated BOLD responses occur in post-traumatic stress disorder (PTSD), bipolar disorder, high body mass index (BMI), and early-life stress-exposed cohorts;^8, 12–15^ In contrast, autosomal recessive Urbach-Wiethe disease, stemming from *ECM1* (coding extracellular matrix protein 1) gene mutations, is marked by a lack of hemodynamic responsiveness: Such patients exhibit BLA vascular perturbations leading to localized calcification, alongside a selective deficit in fear perception when confronted with threat.^16, 17^

Consistent with this clinical phenotype, NVC impairments are recapitulated in animal models, where acute, chronic, and neuroendocrine stressors significantly impair NVC fidelity within stress-sensitive circuits.^18–20^ Yet, a fundamental question of causality remains: do these deficits merely reflect secondary consequences of neuronal dysfunction, or do they actively drive emotional pathology?

Emerging evidence indicates that vascular-derived cues actively modulate diverse neural processes. For instance, nitric oxide generated during NVC promotes adult hippocampal neurogenesis,^21^ while vascular-derived nitric oxide depolarizes the membrane potential of the optic nerve.^22^ In the hypothalamus, salt-loading-induced vasoconstriction during NVC exerts positive feedback to boost local parvalbumin interneuron activity.^23^ Additionally, whole-cell patch recordings in brain slices demonstrate that altered parenchymal arteriolar tone modulates nearby neuronal firing–decreasing pyramidal neuron activity and increasing interneuron activity, with effects depending on the mechanosensitive cation channel.^24^ These observations imply that NVC possesses an intrinsic capacity, likely via chemical and/or mechanical cues, to actively regulate neural circuits, extending beyond its traditional role in metabolic support.^25^

However, its potential as a real-time gatekeeper of emotional circuitry remains unexplored. To address this gap, we investigated NVC within the BLA as a model system. We hypothesize that BLA NVC functions as a critical regulatory checkpoint that calibrates the intensity of negative emotions, thereby maintaining emotional flexibility.^26^

In this study, employing a multidimensional emotion assessment framework combined with cell-type-specific genetic manipulations and arteriolar optogenetics, we demonstrate that NVC in the BLA actively and bidirectionally modulates emotional reactivity. Unlike its non-contribution in the somatosensory barrel cortex (S1BF), we reveal that neurovascular decoupling in the BLA precipitates emotional rigidity—a maladaptive state defined by a collapse in the dynamic range of emotional responses—manifesting as a distinctive biphasic dysregulation where physiological and behavioral outputs are pathologically exaggerated under mild stress, yet paradoxically blunted under intense stress. Mechanistically, we demonstrate that this behavioral rigidity is underpinned by corresponding biphasic shifts in BLA neural activity across graded stressors. Notably, the enhanced NVC model confers emotional resilience against chronic stress by normalizing BLA neural circuit activity. These findings establish NVC as an intrinsic regulatory gain controller in emotional processing and suggest that targeting circuit-specific neurovascular communication offers a novel therapeutic strategy for psychiatric dysregulation.

## RESULTS

### Validation of a quantitative behavioral framework linking graded BLA activation to escalating spatial stress levels

To dissect the neurovascular mechanisms actively modulating emotion, we first established a robust, quantifiable behavioral paradigm that scales linearly with stress intensity. We exposed male age-matched wild-type (*WT*) mice to contexts for 15 min with graded anxiogenic potential: the home cage (HC; minimal stress), the 40 cm, 35 cm, and 10 cm open field tests (OFTs; mild-to-moderate stress), and acute restraint stress (ARS; intense stress) (Fig. 1a).^27–31^ We verified BLA neuronal activation 90 min post-assay by assessing c-Fos expression (Fig. 1b). Immunofluorescence mapping revealed that c-Fos intensity increased in a stress-dependent manner (Fig. 1c). Crucially, this graded BLA activation was mirrored by a stepwise increase in defecation bouts (Fig. 1d), showing a robust linear correlation with BLA c-Fos intensity (*R* = 0.8160, *P* < 0.0001; Fig. 1e). Furthermore, we quantified serum corticosterone levels, the primary stress-responsive glucocorticoid in rodents, in a parallel cohort of mice at 5 min post-assay (Fig.1a). Serum corticosterone concentrations exhibited a stress-dependent trend mirroring that of BLA neural c-Fos (Fig. 1f). Likewise, defecation bouts also showed a strong positive correlation with serum corticosterone concentrations (*R* = 0.7765, *P* = 0.0004; Fig. 1g). Collectively, these results establish defecation as a reliable, robust, and non-invasive marker that accurately tracks systemic and spatially graded stress levels in free-moving live mice.^31–33^

**Fig. 1.**
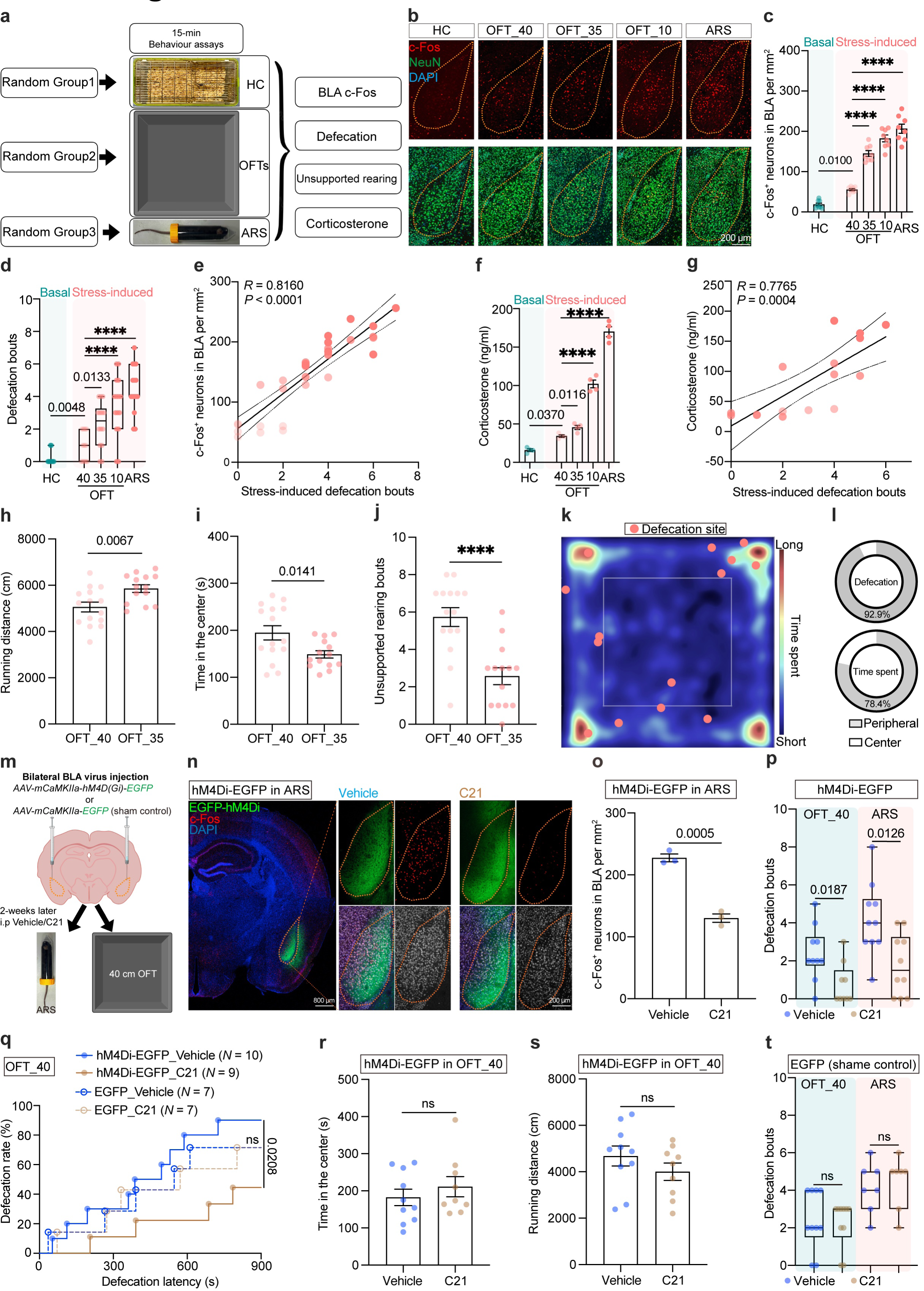
Validation of defecation bouts as a robust quantitative index of negative emotional states. **a** Experimental design for home cage (HC), open field tests (OFTs), and acute restraint stress (ARS) assays. **b, c** Representative images (**b**) and quantification (**c**) of BLA neuronal activation across different stress paradigms shown in (**a**). One-way ANOVA followed by Tukey’s multiple comparisons test (*P* < 0.0001). **d** Quantification of defecation bouts under the paradigms shown in (**a**). Kruskal-Wallis test followed by Dunn’s multiple comparisons test (*P* < 0.0001). **e** Linear correlation between BLA c-Fos-positive neuron counts and defecation bouts. Data were pooled from (**c**) and (**d**). Dotted lines are SEM. **f, g** Serum corticosterone concentrations (**f**) under the paradigms shown in (**a**) and their linear correlation with defecation bouts (**g**). Data were pooled in (**g**) from (**d**) and (**f**). One-way ANOVA followed by Tukey’s multiple comparisons test (*P* < 0.0001). Dotted lines are SEM in (**g**). **h–j** Total running distance (**h**), time spent in the center zone (**i**), and unsupported rearing bouts (**j**) in 40 cm and 35 cm OFT. Student’s *t*-test (**h**, **i**) and Mann-Whitney test (**j**). **k** Representative heatmap of zone preference for a single mouse and the spatial distribution of defecation locations from 15 mice in the 40 cm OFT. Each dot indicates one defecation location. **l** Statistical quantification of the spatial distribution data in (**k**). **m** Experimental schematic of chemogenetic inhibition of BLA neurons using *AAV2/5-hSyn-hM4Di-EGFP*. **n, o** Representative images (**n**) and quantification (**o**) of BLA neuronal activation under chemogenetic inhibition in ARS. Student’s *t*-test. **p** Quantification of defecation bouts in 40 cm OFT and ARS following BLA chemogenetic inhibition. Mann-Whitney test. **q** Defecation rate and defecation latency in the 40 cm OFT. Sample sizes (*N*) are indicated in the panels. Gehan-Breslow-Wilcoxon test. **r, s** Time spent in the center zone (**r**) and total running distance (**s**) following chemogenetic inhibition in the 40 cm OFT. Student’s *t*-test. **t** Defecation bouts in the 40 cm OFT and ARS following BLA injection of the control virus (EGFP). Mann-Whitney test. Except (**k**), each dot represents one mouse. **** indicates *P* < 0.0001. Data are presented as mean ± SEM.

Reducing the arena size from 40 cm to 35 cm prominently increased defecation bouts (Fig. 1d). Although the shift in linear dimensions from 40 cm to 35 cm appears modest, this modification represents a ∼33% reduction in total cubic volume, significantly intensifying the degree of spatial confinement.^34^ This compressed environment provoked a more potent stress state than the 40 cm OFT, characterized by a multi-dimensional escalation in BLA c-Fos density (Fig. 1c), serum corticosterone levels (Fig. 1f), locomotor activity (Fig. 1h), coupled with reductions in classic readouts of exploration, specifically center time (Fig. 1i) and unsupported rearing (Fig. 1j). Spatial mapping further revealed that more than 90% of defecation behavior in the 40 cm OFT occurred in the peripheral zone, consistent with thigmotaxis behavior (Fig. 1k, l).^27, 35^ Together, converging behavioral, neural, and hormonal readouts confirm that the 35 cm OFT elicits a stronger stress response than the standard 40 cm OFT.^36^

To determine whether BLA neuronal activity is necessary for this response, we chemogenetically silenced BLA excitatory neurons using the inhibitory DREADD hM4Di (Fig. 1m).^37^ In mice expressing hM4Di in BLA excitatory neurons treated with the agonist C21 effectively blunted stress-induced BLA c-Fos expression in ARS (Fig. 1n, o). Intriguingly, this also suppressed defecation bouts in both the 40 cm OFT and ARS contexts compared to vehicle-treated control mice (Fig. 1p). Furthermore, BLA silencing significantly reduced the defecation rate (the incidence of mice that defecated) and prolonged the defecation latency (the time elapsed before the production of the first fecal pellet) (Fig. 1q) in the 40 cm OFT. Interestingly, neither center preference nor locomotor activity was altered in the 40 cm OFT following chemogenetic inhibition of the BLA (Fig. 1r-s). Control mice expressing EGFP alone (sham group) exhibited no alterations in defecation metrics (Fig. 1q, t).

Collectively, these data establish a validated, BLA-dependent quantitative framework—the “negative emotion indices”—encompassing BLA neural activation, multi-dimensional defecation metrics (bouts, rate, latency), unsupported rearing, and serum corticosterone concentration to dynamically assess emotional states.

### A genetic model of cerebral SMC-specific GluN2D deficiency exhibits BLA neurovascular decoupling

Equipped with this multi-dimensional assessment framework, we next sought to identify the specific molecular substrates within the BLA neurovascular unit (NVU) that govern these emotional responses. We previously identified that the core N-methyl-D-aspartate (NMDA) receptor subunit GluN1 (encoded by *Grin1*) in smooth muscle cells (SMCs) is essential for NVC at the neuro-smooth muscle junction (NsMJ; Fig. 2a),^38^ but the other regulatory NMDA subunits in SMCs remained to be characterized. To address this, we screened mRNA expression of all NMDA receptor subunits in cultured SMCs sorted from postnatal day 0 (P0) brain parenchymal tissue and identified *Grin2d* (encoding GluN2D) as the sole regulatory subunit detectably expressed (Fig. 2b). High-resolution RNAscope further confirmed that *Grin2d* colocalized specifically with SMC markers in cerebral arterioles with high colocalization coefficients (visualized via *SMA-creER:Ai14* reporter mice; Fig. 2c). To rigorously validate the specificity of our RNAscope probes, we leveraged the well-documented developmental divergence of NMDA subunits in the cerebellum as an internal biological control. We exploited the distinct trajectories where *Grin1* expression increases from P0 to P60, while *Grin2d* expression follows a reciprocal decline.^39, 40^ Our probes accurately recapitulated these age-dependent patterns, thereby confirming their high target specificity (Supplementary information, Fig. S1a, b). This biological validation was further extended to the protein level; our anti-GluN2D antibody successfully distinguished the differential GluN2D protein abundance between P0 and P60 stages in cerebellum tissue, mirroring the transcriptomic data (Supplementary information, Fig. S1c).

**Fig. 2.**
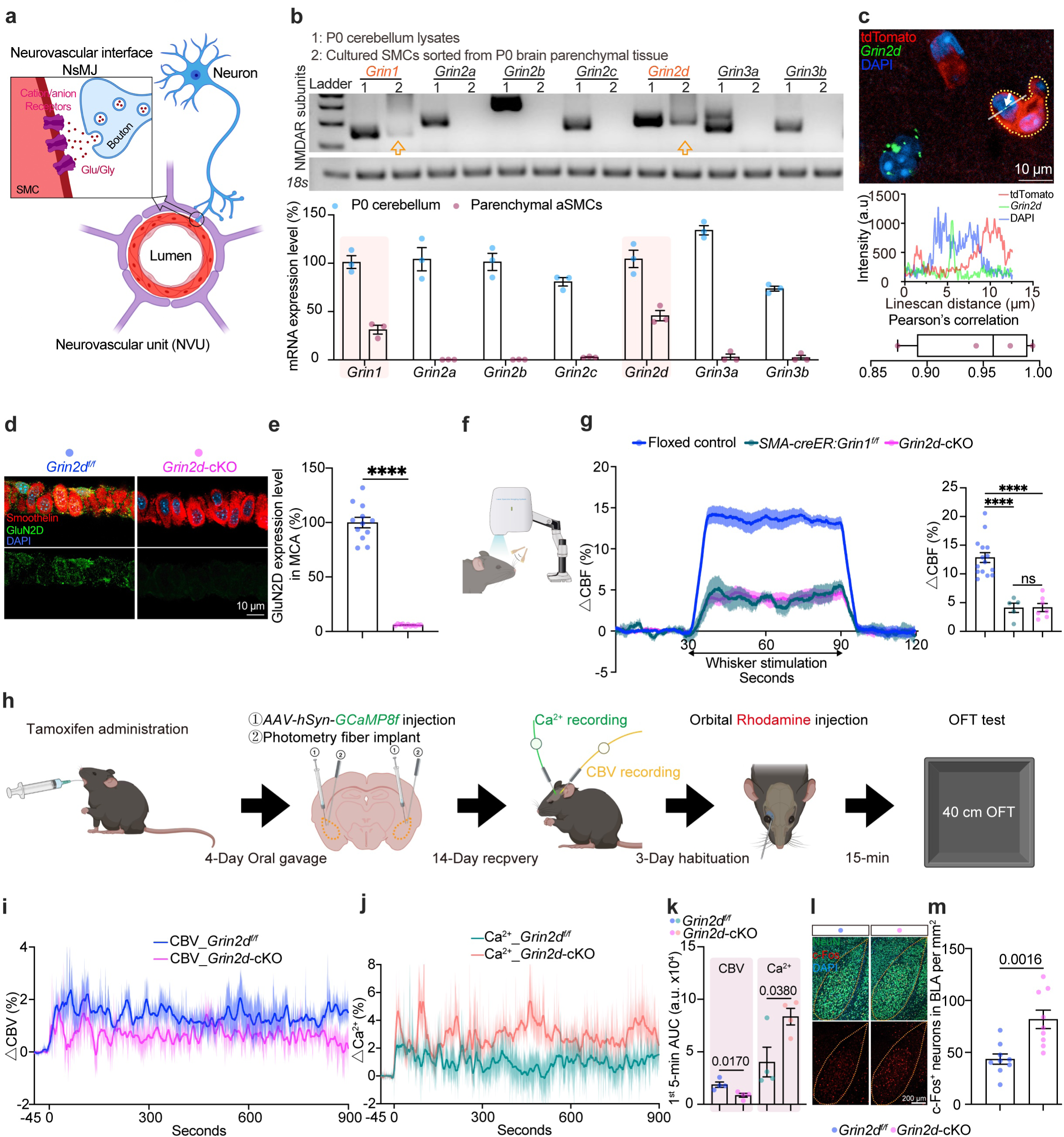
GluN2D in arteriolar SMCs is essential for NVC in S1BF and BLA. **a** Schematic representation of the neuro-smooth muscle junction (NsMJ) within the neurovascular unit (NVU). **b** Electrophoretogram (top) and quantification (bottom) of mRNA levels for NMDA receptor subunits via RT-PCR in cultured SMCs sorted from P0 brain parenchymal tissue. P0 cerebellum lysates used as benchmark. Arrows indicate bands’ locations. **c** Validation of *Grin2d* expression via RNAscope. Top: representative image from the cortex of brain slice (yellow dotted line indicates the arteriolar SMC boundary). Middle: representative fluorescence intensity linescan of the region indicated by white arrows. Bottom: Pearson’s correlation coefficient analysis. Each dot represents one linescan region from one mouse. **d, e** Representative images (**d**) and quantification (**e**) of GluN2D immunofluorescence (IF) staining in the middle cerebral artery (MCA). Each dot represents one analysis region of MCA from a single mouse. Student’s *t*-test. **f-g** Experimental schematic (**f**) and whisker-evoked functional hyperemia in the S1BF (**g**). Panel (**g**) left: time-series traces of functional hyperemia; shaded areas represent SEM across mice. Panel (**g**) right: quantification of the functional hyperemia plateau phase. One-way ANOVA followed by Tukey’s multiple comparisons test (*P* < 0.0001). Floxed control mice group including comparable 8 *Grin2d^f/f^* mice and 7 *Grin1^f/f^* mice. **h** Schematic for concurrent recording of CBV (Through Rhodamine) and neuronal Ca^2+^ (Through *AAV2/9-hSyn-GCaMP8f*) signals in the BLA. **i, j** Time-series recordings of BLA CBV (**i**) and Ca^2+^ signals (**j**) during the 15-min (900 s) 40 cm OFT session. Shaded areas represent SEM across mice. **k** Area under curve (AUC) quantification of the first 5 minutes of CBV and Ca^2+^ responses from data in (**i**) and (**j**). Student’s *t*-test. **l, m** Representative images (**l**) and quantification (**m**) of BLA neuronal activation following the 40 cm OFT. Student’s *t*-test. Except (**c** and **e**), each dot represents one mouse. **** indicates *P* < 0.0001. Data are presented as mean ± SEM. See also Supplementary information, Figs. S1–S8.

Using this validated antibody, immunofluorescence analysis of isolated middle cerebral arteries (MCA) (Fig. 2d, e) and cultured SMCs sorted from P0 brain parenchymal tissue (Supplementary information, Fig. S1d) confirmed the specific ablation of GluN2D in the cerebral SMCs in *SMA-creER:Grin2d^f/f^* (*Grin2d*-cKO) mice. Furthermore, Western blot analysis of whole-cortical lysates revealed that *Grin2d*-cKO did not affect GluN2D expression in other non-SMC populations (Supplementary information, Fig. S1e), confirming the cell-type specificity of our genetic model. To assess the global impact of this genetic depletion, we examined GluN2D expression in the SMCs of peripheral organs, including the heart, liver, spleen, lung, kidney, intestine, and bladder (Supplementary information, Fig. S2a). Notably, GluN2D expression seemed restricted to cerebral SMCs and absent in these peripheral tissues’ SMCs, suggesting that our *Grin2d*-cKO model exerts a brain-restricted impact (Supplementary information, Fig. S2b, c). Furthermore, we systematically verified the physiological status of *Grin2d*-cKO mice; no differences were observed in systemic metabolic homeostasis (Supplementary information, Fig. S3a-g), blood pressure (Supplementary information, Fig. S3h), gastrointestinal function (Supplementary information, Fig. S3i-n), and blood-brain barrier (BBB) integrity^41, 42^ (Supplementary information, Fig. S3o-p) compared to littermate controls. Finally, we assessed key contractile proteins, finding that α-SMA and smoothelin expression remained unaltered (Supplementary information, Fig. S4). These data demonstrate that the basic metabolic functions, gastrointestinal system, and basic contractile structural integrity of SMCs are preserved in *Grin2d*-cKO mice.

To assess the functional impact of SMC-specific *Grin2d* deletion and facilitate a direct, one-to-one comparison with our previously reported *Grin1*-cKO model, we employed the same classical paradigm: measuring whisker stimulation-evoked hemodynamic responses in the somatosensory barrel cortex (S1BF) (Fig. 2f).^38^ Consistently, *Grin2d*-cKO mice exhibited a ∼66% attenuation in functional hyperemia upon whisker stimulation (Fig. 2g), confirming that GluN2D is functionally comparable to GluN1 in regulating cerebral NVC. Control experiments confirmed that tamoxifen administration alone in *Grin2d^f/f^* mice did not alter NVC responses (Supplementary information, Fig. S5).

Having established the NVC deficit in the cortex, we sought to determine whether this NVC dysfunction extends to the BLA. To this end, we developed a *de novo* system based on previous strategies^43, 44^ to simultaneously record neuronal activity and blood supply within the BLA of freely moving mice, thereby enabling direct assessment of NVC. First, we mapped the BLA arteriolar topology by superimposing 3D-reconstructed arterioles in the tissue-cleared brain on a schematic cerebral vasculature. This analysis identified that the corticoamygdaloid artery (coamg) penetrates the BLA, making it suitable for photometric monitoring (Supplementary Information, Fig. S6, Video S1). Second, we validated the fidelity of our cerebral blood volume (CBV) recording method: fiber photometry-based blood plasma fluorescence intensity recording locally in the BLA showed a robust linear correlation with orbital Rhodamine injection concentration (Supplementary information, Fig. S7a, b; *R* = 0.8946, *P* < 0.0001). Signal stability was achieved 15 min post-injection (Supplementary information, Fig. S7c), and this time point was designated as the initiation time for the behavior assay.

Using this optimized paradigm, after tamoxifen administration, we injected *AAV2/9-hSyn-GCaMP8f* into the BLA and implanted optic fibers at the injection site two weeks before the recordings. After three days of habituation (one trial per day for a continuous 3-day period), we monitored NVC during the 40 cm OFT (Fig. 2h-k). Consistent with our earlier c-Fos data (Fig. 1c), the 40 cm OFT elicited robust BLA Ca^2+^ transients that were temporally matched by CBV elevations, indicating tight local NVC in littermate control mice (Fig. 2i-k). In contrast, *Grin2d-cKO* mice exhibited a dissociated neurovascular phenotype, where augmented BLA neuronal Ca^2+^ signals were paradoxically paired with a blunted hemodynamic response (Fig. 2i-k). This inverse pattern—hyperactive neurons coupled with hypoperfused blood supply—was further corroborated by increased c-Fos expression in the BLA 90 min post-assay (Fig. 2l, m). Furthermore, we confirmed that the observed attenuation in functional hyperemia was not attributable to differences in basal perfusion, locomotor activity, or spatial preference within the 40 cm OFT (Supplementary information, Fig. S8)

Collectively, these findings demonstrate robust NVC within the BLA during 40 cm OFT exposure and validate that SMC-specific *Grin2d*-cKO mice exhibit an NVC deficit in this region.

### BLA-target pharmacological restoration of NVC reversed emotional hyper-reactivity in *Grin2d-cKO* mice under mild stress

Building on the BLA’s established role as one of the central hubs for stress responses, we next utilized our negative emotion indices (Fig. 1) to evaluate whether the BLA neurovascular decoupling observed in *Grin2d*-cKO mice (Fig. 2) results in behavioral and physiological impairments. Notably, we observed that *Grin2d*-cKO mice exhibited a ∼12-fold increase in defecation bouts during the first 5 min of the 40 cm OFT, a trend sustained throughout the session (Fig. 3a). This phenomenon was not observed in the home cage context (Supplementary information, Fig. S3c-e, n). Littermate control mice (*Grin2d^f/f^*) displayed a graded increase in defecation bouts in response to escalating spatial stress, consistent with the pattern observed in *WT* mice (Figs. 1d and 3b). In contrast, *Grin2d-*cKO mice failed to show this graded response; instead, they exhibited an abrupt rise to a maximal plateau level of defecation bouts, a response typically elicited by intense ARS stress (Fig. 3b). This hyper-reactivity was further reflected in an increased defecation rate, reduced latency (Fig. 3c), a more than 3-fold reduction in unsupported rearing, and a ∼1.3-fold increase in serum corticosterone levels in the 40 cm OFT (Fig. 3d). Importantly, basal stress levels in the home cage remained indistinguishable between genotypes (Supplementary information, Fig. S9a). Altogether, *Grin2d*-cKO mice displayed a unified shift across multiple-dimensional stress metrics, indicating a robust, exaggerated negative emotional response to mild stressors, such as those encountered in OFTs.

**Fig. 3.**
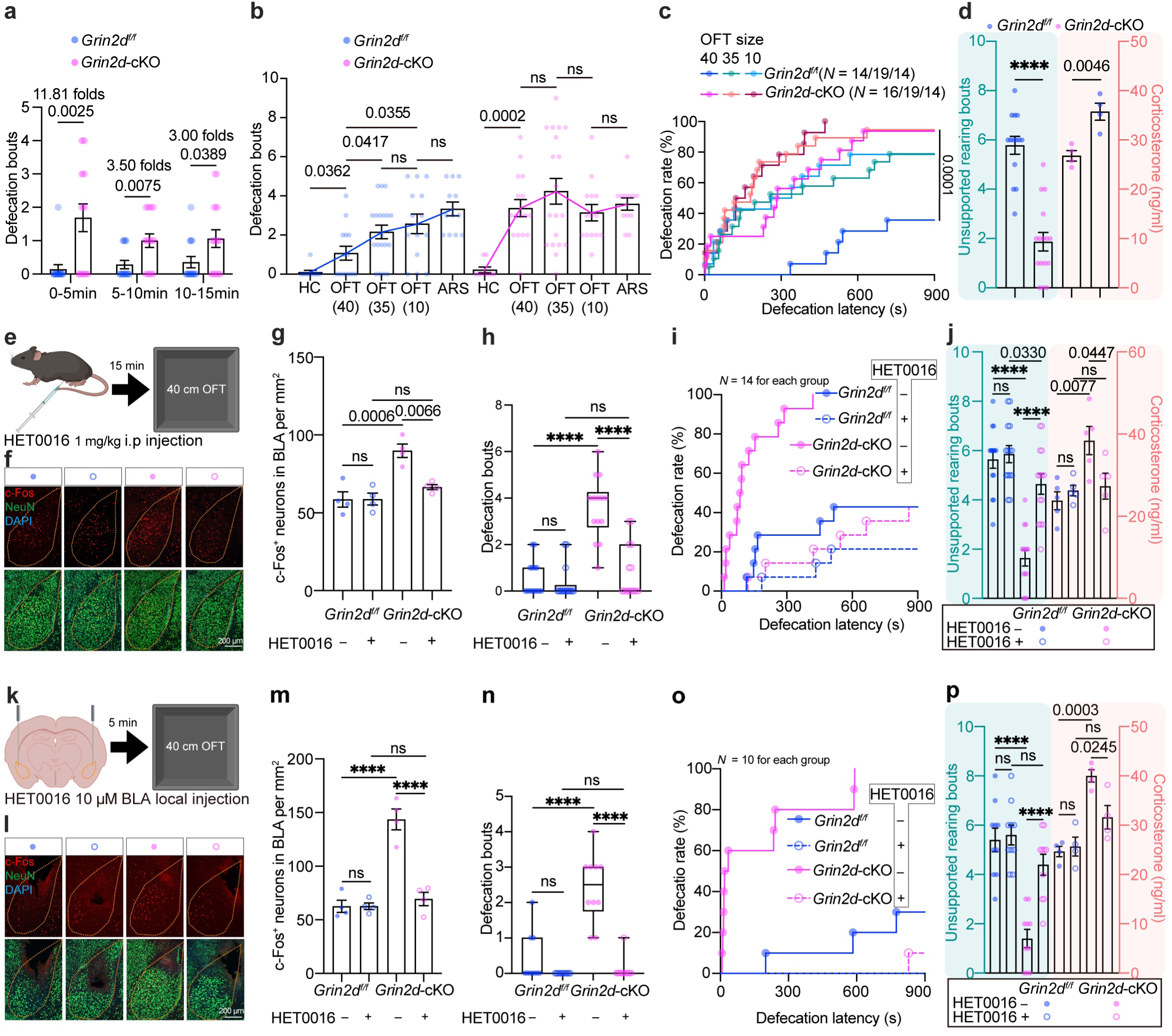
Systemic SMC-specific GluN2D deficiency-driven negative emotionality is pharmacologically rescued via BLA local intervention. **a** Defecation bout in the 40 cm OFT shown in 5-minute intervals. Mann-Whitney test. **b** Comparison of defecation bouts across HC (Home cage), OFTs, and ARS paradigms. Kruskal-Wallis test followed by Dunn’s multiple comparisons test (*P* = 0.0365). **c** Defecation rate and defecation latency in the OFTs. Sample sizes (*N*) are indicated in the panels. Gehan-Breslow-Wilcoxon test. **d** Unsupported rearing bouts (left) and serum corticosterone concentrations (right) in the 40 cm OFT. Mann-Whitney test (left) and Student’s *t*-test (right). **e** Experimental schematic of systemic HET0016 administration. **f, g** Representative images (**f**) and quantification (**g**) of BLA neuronal activation following systemic HET0016 treatment in the 40 cm OFT. Two-way ANOVA followed by Tukey’s multiple comparisons test (*P* = 0.0104). **h–j** Quantification of defecation bouts (**h**), defecation rate and latency (**i**), and unsupported rearing bouts and serum corticosterone concentrations (**j**) in the 40 cm OFT following systemic HET0016 administration. Two-way ANOVA on rank-transformed data followed by Sidak’s multiple comparisons test (**h**, *P* < 0.0001; **j left**, *P* = 0.00003), Gehan-Breslow-Wilcoxon test (**i**), and Two-way ANOVA followed by Tukey’s multiple comparisons test (**j right**, *P* = 0.0212). Sample sizes (*N*) are indicated in the panel (**i**). **k** Experimental schematic of local BLA microinjection of HET0016. **l, m** Representative images (**l**) and quantification (**m**) of BLA neuronal activation following local BLA HET0016 treatment in the 40 cm OFT. Two-way ANOVA followed by Tukey’s multiple comparisons test (*P* < 0.0001). **n–p** Quantification of defecation bouts (**n**), defecation rate and latency (**o**), and unsupported rearing bouts and serum corticosterone concentrations (**p**) in the 40 cm OFT following local BLA HET0016 administration. Two-way ANOVA on rank-transformed data followed by Sidak’s multiple comparisons test (**n**, *P* < 0.0001, **p left**, *P* = 0.0020), Gehan-Breslow-Wilcoxon test (**o**), and Two-way ANOVA followed by Tukey’s multiple comparisons test (**p right,** *P* = 0.0190). Sample sizes (*N*) are indicated in the panel (**o**). Each dot represents one mouse. **** indicates *P* < 0.0001. Data are presented as mean ± SEM. Detailed statistical data for panel (**c**, **i**, and **o**) are shown in the supplementary table S1, sub-table 1-3. See also Supplementary information, Figs. S9–S12.

Given that *Grin2d*-cKO mice showed NVC dysfunction beyond the BLA, to determine the contribution of BLA local NVC in hyper-reactivity, we employed administration of HET0016, a selective inhibitor of CYP450ω-hydroxylases that suppresses the synthesis of 20-HETE (a potent vasoconstrictor), thereby facilitating vasodilation to compensate for the NVC deficit.^45, 46^

To strictly validate the pharmacological efficacy of HET0016 before its site-specific delivery to the BLA, we first replicated the systemic administration protocol described in our previous study.^38^ Systemic HET0016 administration dose-dependently reduced ARS-induced defecation bouts (Supplementary Information, Fig. S10a). This behavioral rescue provided a clear rationale for selecting 1 mg/kg as the optimal dose for subsequent investigations. Under this condition, we confirmed that 1 mg/kg HET0016 produced a ∼3-fold increase in whisker-evoked functional hyperemia amplitude in the S1BF compared with saline controls (Supplementary Information, Fig. S10b, c)

Notably, HET0016 effectively rescued the hyper-reactive phenotype in *Grin2d*-cKO mice, both in systemic (Fig. 3e-j) and local (Fig. 3k-p) administrations in the 40 cm OFT. Both administration routes effectively normalized BLA neural activation and multi-dimensional defecation metrics—including bouts, rate, and latency—to littermate control levels (Fig. 3f-i and l-o). Importantly, this vascular intervention also significantly normalized unsupported rearing bouts and reduced serum corticosterone concentrations (Fig. 3j and p) in the 40 cm OFT. In contrast, in the home cage, neither systemic nor BLA local HET0016 impacted serum corticosterone levels (Supplementary information, Fig. S9b, c).

Collectively, local pharmacological compensation in the BLA sufficiently normalized the hyper-reactivity features in the *Grin2d*-cKO genetic model, despite NVC deficits in other brain regions, thereby establishing that intact NVC in the BLA is essential for maintaining appropriate emotional responses to mild stress.

We observed consistent behavioral phenotypes in the 40 cm OFT across various mechanistically distinct models of NVC dysfunction, each targeting different nodes of the NVU (Supplementary information, Fig. S11, S12). Specifically, either perturbing upstream neural signaling (*Emx1-cre:Ptgs2^f/f^*) (Supplementary information Fig. S11a-h, and S12o-p)^47–49^ or disrupting the neurovascular interface–achieved via pan-mural cell-specific *Grin2d*-cKO (*Pdgfrβ-creER:Grin2d^f/f^*) or SMC-specific *Grin1-Grin2d* double-cKO (*SMA-creER:Grin1^f/f^:Grin2d^f/f^*) (Supplementary information Fig. S11i-r and S12o-p)–consistently recapitulated the NVC deficits and behavioral phenotypes observed in the primary *Grin2d*-cKO mice in the 40 cm OFT (Fig. 2, 3).

To ultimately disturb contractile protein machinery within SMC, we targeted the *Acta2* gene (encoding α-SMA).^50, 51^ This model (*Pdgfrβ-creER:Acta2^f/f^* mice, shorten as *Acta2*-cKO) indistinguishably mirrored the hyper-reactive phenotypes of the signaling-deficient models in the 40 cm OFT (Supplementary information, Fig. S11s-v, x-aii, and S12o, p). Surprisingly, a high-fat diet (HFD)—known to diminish NVC in both animals and humans via multiple factors^14, 52^—produced an identical phenotype of *Grin2d*-cKO mice in the 40 cm OFT (Supplementary information, Fig. S12a-d, g-h, j-n, o, and p). These findings indicate that this emotional hyper-reactivity was not restricted to the single *Grin2d*-cKO model but represented a universal consequence of neurovascular dysfunction within the 40 cm OFT, reinforcing the universality of NVC-driven emotional dysregulation.

Although local HET0016 delivery to the BLA successfully restored emotional metrics in other models (Fig. 3k-p, Supplementary information, Fig. S12i-n) in the 40 cm OFT, it failed to produce similar effects in *Acta2*-cKO mice (Supplementary information, Fig. S11w-aii) in the same context. This indicates that once the downstream contractile apparatus is disrupted, upstream regulators are unable to exert their regulatory functions. Finally, we confirmed that these convergent effects did not involve BBB leakage (Supplementary information, Fig. S3o, p; S12e, f).^53^ Together, these findings establish that NVC integrity per se is indispensable for emotional homeostasis.

### Caldesmon acts as the downstream effector of GluN2D in NVC thereby modulating emotional reactivity

Given that direct disruption of the vascular contractile apparatus (*Acta2*-cKO) phenocopies the emotional hyper-reactivity of *Grin2d*-cKO mice (Fig. 3; Supplementary information, Fig. S11; Fig. S12), we hypothesized that GluN2D couples to the SMC cytoskeletal machinery. To this end, we performed co-immunoprecipitation (Co-IP) followed by mass spectrometry to map the GluN2D interactome in vascular SMCs. This unbiased screen identified caldesmon (encoded by *Cald1*), an actin-binding protein critical for regulating actomyosin contractility, as a top candidate enriched in *Vascular Smooth Muscle Contraction* pathway (Fig. 4a).^54, 55^ This physical association between GluN2D and caldesmon was further confirmed by Western blotting of the immunoprecipitated complex (Fig. 4b).

**Fig. 4.**
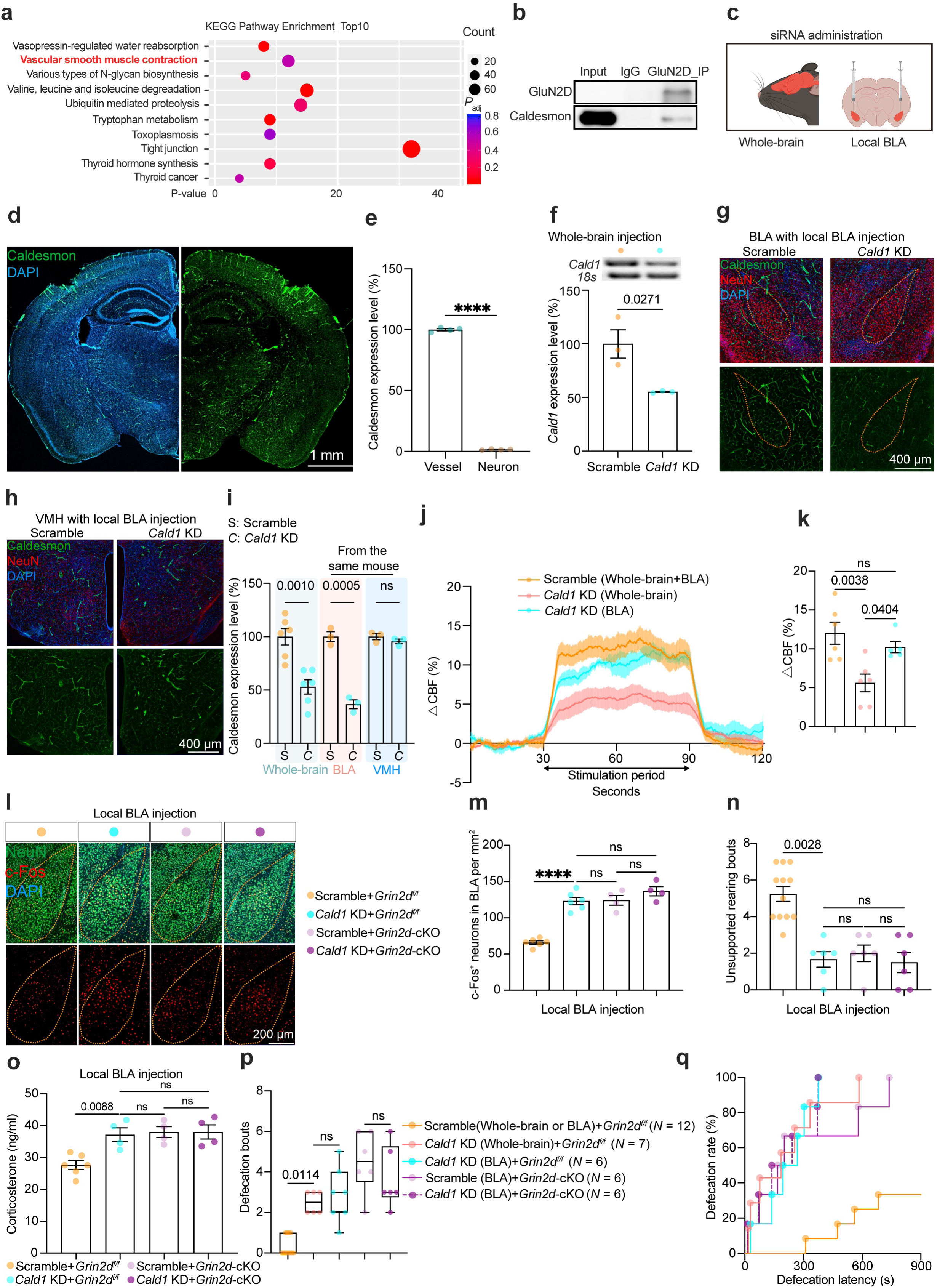
Caldesmon interacts with GluN2D in the SMCs to regulate neurovascular coupling and further modulate emotional states. **a** KEGG pathway analysis of GluN2D-interacting proteins identified via proteomic analysis from cultured SMCs sorted from P0 brain parenchymal tissue. **b** Immunoprecipitation validation of the interaction between GluN2D and caldesmon from cultured SMCs sorted from P0 brain parenchymal tissue. **c** Experimental schematic for whole-brain (left) and local BLA (right) *Cald1* knockdown (KD) using siRNA. **d, e** Representative images (**d**) and quantification (**e**) of caldesmon immunofluorescence (IF) staining. Student’s *t*-test. **f** Electrophoretogram (top) and quantification (bottom) of *Cald1* mRNA expression in the cortex lysates following whole-brain *Cald1* siRNA injection. Student’s *t*-test. **g, h** Representative immunofluorescence (IF) images of caldesmon in the BLA (**g**) and ventromedial hypothalamus (VMH, **h**) of mice received local BLA *Cald1* siRNA injection. **i** Quantification of caldesmon expression levels from the experiments shown in (**g, h)**. Student’s *t*-test. **j, k** Functional hyperemia in the S1BF during whisker stimulation (**j**) and quantification of the plateau phase (**k**). Shaded areas represent SEM across mice. One-way ANOVA followed by Tukey’s multiple comparisons test (*P* = 0.0041). Floxed control mice group including comparable 3 mice with cisterna magna scramble siRNA injection and 3 mice with local BLA scramble siRNA injection. **l, m** Representative images (**l**) and quantification (**m**) of BLA neuronal activation in the 40 cm OFT. One-way ANOVA followed by Tukey’s multiple comparisons test (*P* < 0.0001). **n** Unsupported rearing bouts in the 40 cm OFT. Kruskal-Wallis test followed by Dunn’s multiple comparisons test (*P* = 0.0001). **o** Serum corticosterone concentrations in the 40 cm OFT. One-way ANOVA followed by Tukey’s multiple comparisons test (*P* = 0.0013). **p, q** Defecation bouts (**p**), defecation rate, and latency (**q**) in the 40 cm OFT. Kruskal-Wallis test followed by Dunn’s multiple comparisons test (**p**, *P* < 0.0001) and Gehan-Breslow-Wilcoxon test (**q**). Sample sizes (*N*) are indicated between panels (**p** and **q**). Each dot represents one mouse. **** indicates *P* < 0.0001. Data are presented as mean ± SEM. Detailed statistical data for panel (**q**) are shown in the supplementary table S1, sub-table 4. See also Supplementary information, Figs. S13–S14.

Caldesmon serves as a “blocker” on smooth muscle cells’ contraction; its disruption is predicted to impair cross-bridge cycling and thereby compromise NVC. To test this, we employed siRNA-mediated knockdown (KD). Unlike GluN2D, caldesmon expression was restricted exclusively to vascular mural cells (Fig. 4c–e),^56^ allowing for vascular-specific manipulation without the risk of direct neuronal off-target effects. We validated the efficacy and spatial specificity of this approach: both whole-brain and local BLA siRNA delivery robustly reduced caldesmon expression in background-matched groups (both established in *Grin2d^f/f^* mice) (Fig. 4f-i). Crucially, gene silencing was spatially restricted to the injection site with anatomical precision: local BLA *Cald1* KD did not impact caldesmon levels in the adjacent Ventromedial Hypothalamus (VMH) (Fig. 4g–i), nor did S1BF-restricted *Cald1* KD alter expression in the BLA (Supplementary information, Fig. S13).

Functionally, local S1BF *Cald1* KD—but not local BLA *Cald1* KD—attenuated whisker-evoked functional hyperemia in the barrel cortex (Fig. 4j, k), confirming *Cald1* KD as an effective tool for disrupting NVC with regional precision. We then focused on the emotional circuit. Behaviorally, both whole-brain and local BLA *Cald1* KD recapitulated the hyper-reactive phenotype of *Grin2d*-cKO mice in the 40 cm OFT: *Cald1* KD mice exhibited exaggerated negative emotional responses, characterized by BLA neuronal hyperactivation (c-Fos), reduced unsupported rearing, elevated serum corticosterone concentrations, and intensified defecation dynamics (increased bouts/rate, decreased latency) (Fig. 4l–q). In contrast, local S1BF *Cald1* KD did not alter the above parameters, confirming regional specificity (Supplementary information, Fig. S14).

To elucidate the functional relationship between GluN2D and caldesmon, we applied the local BLA *Cald1* KD approach in *Grin2d*-cKO mice. This epistasis analysis showed no additive phenotype (Fig. 4l-q), suggesting that caldesmon likely functions downstream of GluN2D within a hierarchical vascular signaling cascade. Together, these results identify caldesmon as a critical effector linking GluN2D to SMC contractility and demonstrate that localized NVC deficits in the BLA–despite intact NVC in other brain regions–are sufficient to impair emotional responses to mild stress.

### BLA arteriolar SMCs’ optogenetic suppression of NVC precipitates negative emotional states under conditions of mild and negligible stress

To achieve acute suppression of NVC within a millisecond timescale–unattainable through genetic or pharmacological manipulations–we employed an SMC-optogenetic approach to reversibly “clamp” vascular diameter.^57^

We generated SMC-specific ChR2-expressing mice (*SMA-creER:Ai32;* Fig. 5a) and optimized a fiber-optic setup to locally activate BLA SMCs while simultaneously monitoring regional CBV (Fig. 5b). In the home cage under minimal external stress, we optimized a protocol that reliably induced a consistent ∼3% CBV decrease (Fig. 5c red line). This hemodynamic manipulation was rapid and reversible (Fig. 5d).

**Fig. 5.**
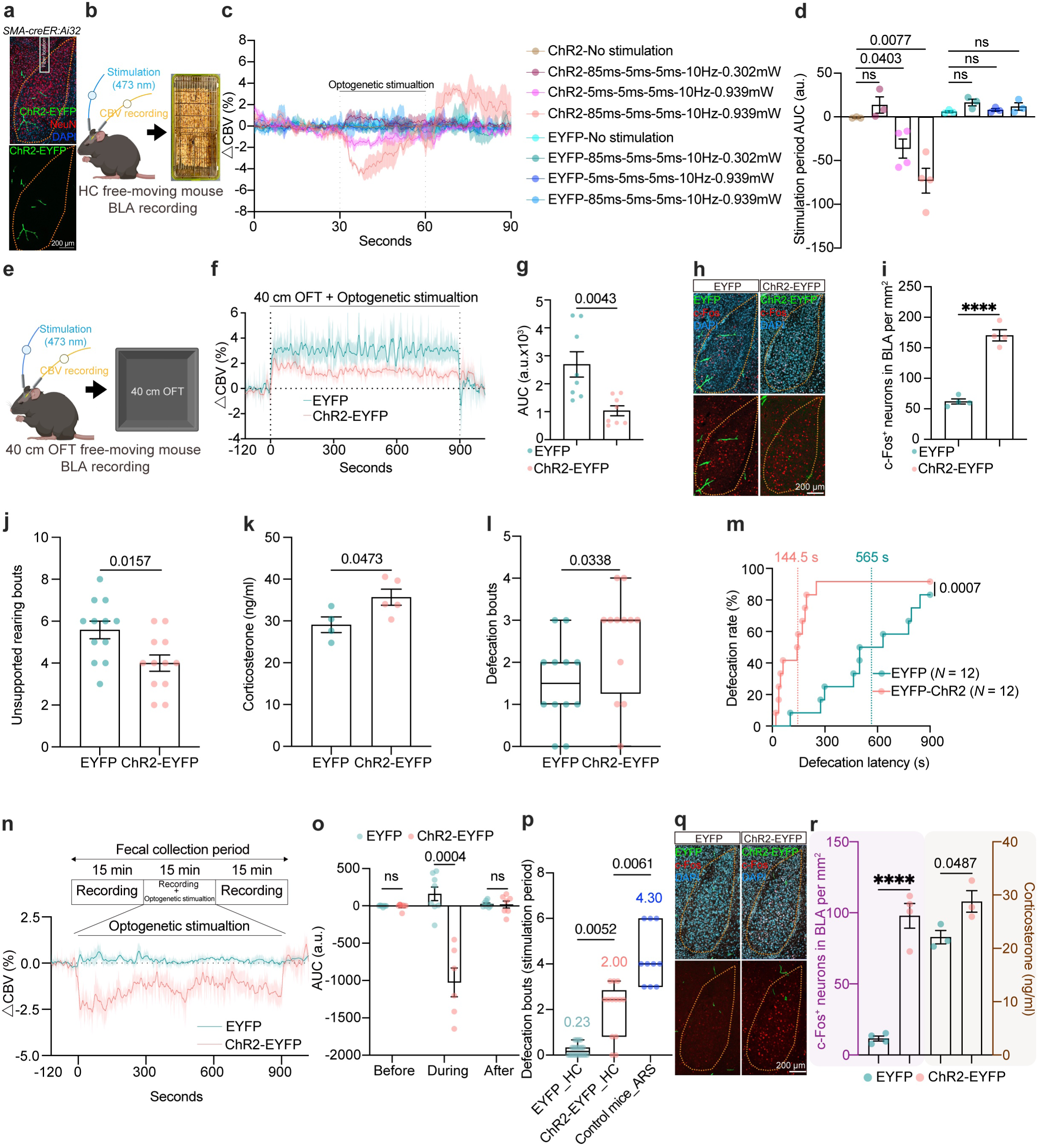
Optogenetic vascular clamping in the BLA is sufficient to drive negative emotionality. a, **b** Experimental schematic showing the fiber optic implantation site (**a**) and the setup for optogenetic stimulation in a freely moving mouse within the home cage (**b**). **c, d** Representative CBV traces showing optogenetically induced CBV changes under various stimulation protocols (**c**) and the corresponding quantification (**d**). Each dot represents the area under curve (AUC) change from one mouse during the stimulation period. Shaded areas represent SEM across mice. One-way ANOVA followed by Tukey’s multiple comparisons test (*P* < 0.0001). **e–g** Experimental schematic (**e**), representative CBV traces (**f**), and quantification (**g**) in the 40 cm OFT under optogenetic vascular clamping. Each dot represents the AUC change during the optogenetic stimulation from one mouse in (**g**). Shaded areas represent SEM across mice in (**f**). Dotted lines in (**f**) indicate *X* = 0 and *Y* = 0. Student’s *t*-test. **h, i** Representative images (**h**) and quantification (**i**) of BLA neuronal activation following optogenetic stimulation in the 40 cm OFT. Student’s *t*-test. **j–m,** Quantification of unsupported rearing bouts (**j**), serum corticosterone concentrations (**k**), defecation bouts (**l**), and defecation rate and latency (**m**) in the 40 cm OFT during optogenetic stimulation. Coloured dotted lines in (**m**) indicate the median defecation latency for each group. Mann-Whitney test (**j**, **l**), Student’s t-test (**k**), and Gehan-Breslow-Wilcoxon test (**m**). Sample sizes (*N*) are indicated in the panel (**m**). **n, o** Experimental schematic of 15-min home cage optogenetic stimulation (**n**, **top**), corresponding BLA CBV response (**n**, **bottom**), and quantification (**o**). Each dot represents the area under curve (AUC) change during the optogenetic stimulation from one mouse in the home cage. Shaded areas represent SEM across mice. Student’s *t*-test. **p** Quantification of defecation bouts during home cage optogenetic stimulation. Colored numbers indicate the mean defecation bouts for each group. Kruskal-Wallis test followed by Dunn’s multiple comparisons test (*P* < 0.0001). **q, r** Representative images (**q**) and quantification (**r**) of BLA neuronal activation (**r left**) and serum corticosterone concentrations (**r right**) in the home cage optogenetic stimulation. Student’s *t*-test. Each dot represents one mouse. **** indicates *P* < 0.0001. Data are presented as mean ± SEM. See also Supplementary information, Figs. S15–S16.

Consistent with earlier data obtained from *Grin2d^f/f^* mice (blue line in Fig. 2i, blue line), the stress during the 40 cm OFT elicited functional hyperemia in an independent cohort of *SMA-creER:Ai47* (EYFP) mice (Fig. 5e-g, green line). Optogenetic clamping of BLA arterioles partially blocked this hemodynamic response (Fig. 5e–g, red line). Furthermore, power spectral density analysis of the hemodynamic traces revealed a significant reduction in blood flow power across low-frequency bands—typically associated with vascular vasomotion—during optogenetic stimulation (Supplementary information, Fig. S15). This suppression of spectral energy confirms that optogenetic clamping effectively stabilized vascular diameter.

This acute NVC suppression precipitated an emotional shift: ChR2-EYFP mice exhibited an exacerbation of negative emotional responses, characterized by markedly increased BLA neural activation (Fig. 5h, i), reduced unsupported rearing (Fig. 5j), and elevated serum corticosterone concentrations (Fig. 5k), and shifted defecation metrics (Fig. 5l, m) in the 40 cm OFT. These alterations were consistent with the *Grin2d*-cKO phenotype (Fig. 3a-d) in the 40 cm OFT. Locomotor activity and center preference remained unaffected, ruling out motor confounds (Supplementary information, Fig. S16a, b) in the 40 cm OFT. This optogenetically driven, temporally controlled suppression of NVC reveals that preventing the stress-induced vascular surge removes a critical physiological component, thereby amplifying negative emotion.

Next, we investigated whether BLA-targeted vascular clamp, in the absence of actual external stress, is sufficient to drive stress-like behaviors. We applied the same optogenetic stimulation protocol (∼3% CBV decrease) to mice in the safe home cage environment for 15 min (Fig. 5n, o). Strikingly, this hemodynamic mismatch alone was sufficient to induce a stress-like state: While basal defecation was negligible in the home cage (0.23 pellets/15 min), optogenetic stimulation induced a ∼6.4-fold cumulative CBV change (Fig. 5o) coupled with a ∼10-fold surge in defecation bouts (Fig. 5p). These changes were associated with increased BLA neural activation and elevated serum corticosterone levels (Fig. 5q, r).

Crucially, we verified that this optogenetic manipulation did not induce hypoxic injury. Unlike the Stroke Induced by Magnetic ParticLEs (SIMPLE) stroke model^58–60^ which triggers robust HIF-1α upregulation, our optogenetic protocol elicited no obvious HIF-1α response after optogenetic stimulation in the home cage (Supplementary information, Fig. S16c, d). This confirms that the observed behavioral shifts stem from functional neurovascular modulation, not ischemic stroke or tissue hypoxia.

### NVC dysfunction conversely exhibited hyporeactivity under high-intensity stress

Our findings from the standard 40 cm OFT established that NVC dysfunction precipitates a hyper-reactive, anxiety-like state under mild stress (Fig. 3). We next characterized how the NVC-deficient BLA responds to imminent, life-threatening high-intensity stress.^61^ To this end, we employed two paradigms: a looming assay^62^ and a virtual reality (VR_eagle_) simulation of a diving eagle^63^, both of which mimic an approaching aerial predator.

Prior to looming stimulation, basal locomotor activity was comparable between groups (Fig. 6a, blue region; Supplementary information, Fig. S17a). Upon stimulus, both groups exhibited immediate and robust freezing (Fig. 6a, yellow region), confirming that NVC dysfunction did not impair sensory detection, threat evaluation, or the initiation of defensive strategies.^64^

**Fig. 6.**
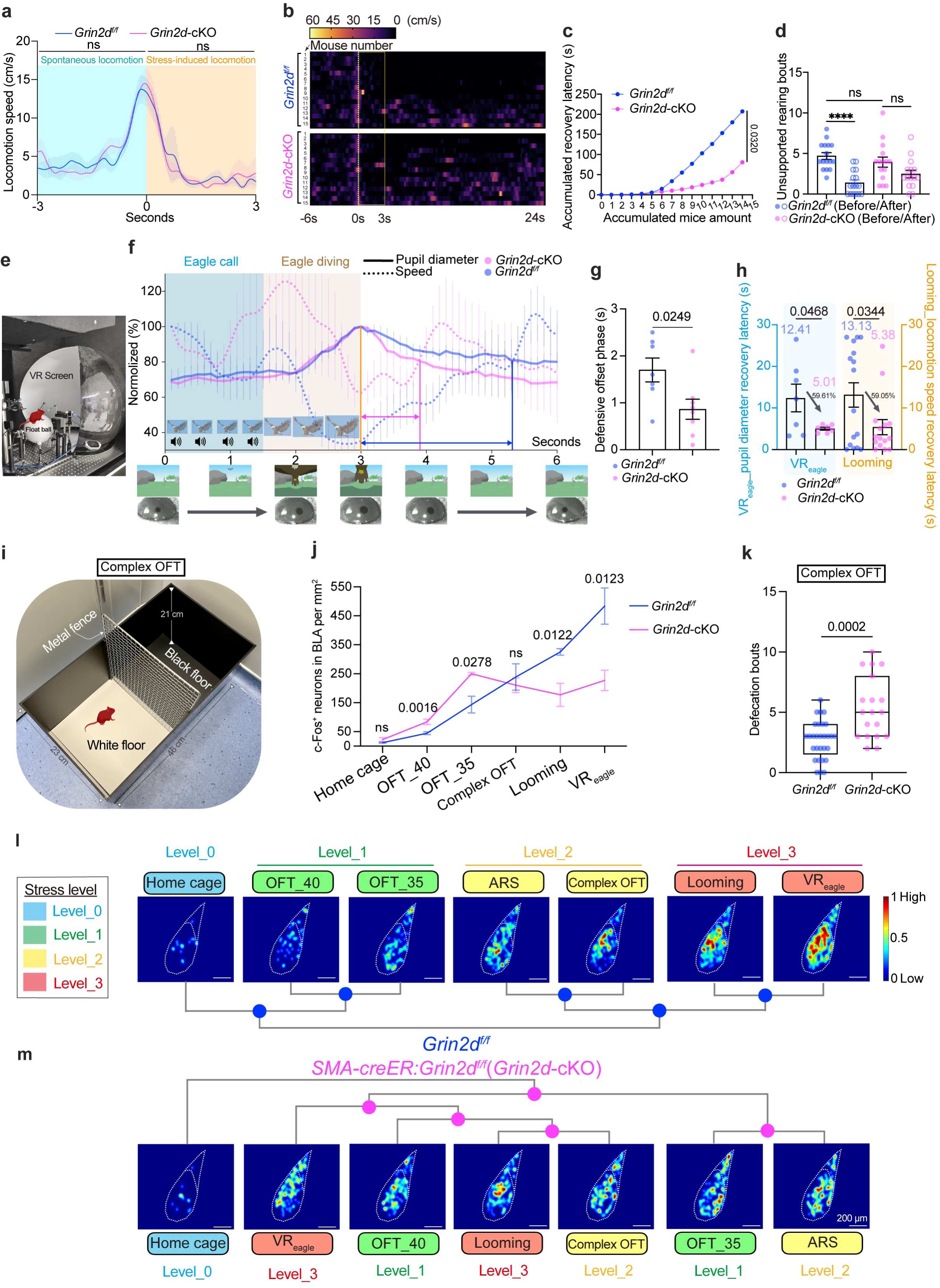
Compromised NVC impairs defensive sustainability and alters BLA neural topological scaling. a, **b** Time-series plots (**a**) and heatmaps (**b**) profiling locomotion speed dynamics before (**a, blue region**) and during (**a, yellow region**) looming stimulation. shaded areas represent SEM across mice in (**a**). The yellow dotted box in (**b**) indicates the 3-second looming duration. **c, d** Quantification of the accumulated recovery latency from freezing (**c**) and unsupported rearing bouts 5 min before and after looming stimulation (**d**). *N* = 15 mice per group. Kolmogorov-smirnov test (**c**) and Two-way ANOVA on rank-transformed data followed by Sidak’s multiple comparisons test (**d**, *P* < 0.0001). **e** Experimental schematic of the virtual reality facility for visual threat stimulation. **f** Time-series traces showing normalized pupil diameter and locomotion speed during (0–3 s) and after (3–6 s) VR_eagle_ stimulation. The blue region (first 1.5 s) indicates the eagle call, and the yellow region (1.5–3.0 s) indicates the eagle diving phase. Representative images of the VR_eagle_ stimulus and the mouse eye with a clearly visualized pupil are shown on the bottom. Shaded areas represent SEM across mice. **g, h** Quantification of defensive offset phase (**g**) dynamics and recovery latencies for pupil diameter and locomotion (**h**) from the data in (**b** and **f**). *N* = 7 mice per group in VR_eagle_ assay, and *N* = 15 mice per group in looming assay. Student’s *t*-test. **i** Experimental schematic of the complex open field test (c-OFT) arena. **j** Quantification of BLA neuronal activation in contexts of home cage, 40 cm OFT, 35 cm OFT, c-OFT, Looming, and VR_eagle_ assays. Data of 40 cm OFT groups are shared with Fig. 2m. Student’s *t*-test. **k** Defecation bouts of mice in the c-OFT. Mann-Whitney test. **l, m** Structural Similarity Index (SSIM)-based unbiased classification of BLA c-Fos topological patterns across seven standard stress contexts in littermate control mice (**l**) and *Grin2d*-cKO mice (**m**). Heatmaps indicate the modified c-Fos intensity in the BLA (processing details can be seen in the supplementary information, **Figure S18a**). **** indicates *P* < 0.0001. Data are presented as mean ± SEM. See also Supplementary information, Figs. S17–S18.

However, a striking divergence emerged during the maintenance phase: while control mice sustained a prolonged freezing state, *Grin2d*-cKO mice exhibited a “premature exit” phenotype (Fig. 6b). Velocity heatmaps confirmed that *Grin2d*-cKO mice resumed locomotor activity significantly earlier than controls (Fig. 6b, c). This behavioral collapse was mirrored by an inappropriate increase in unsupported rearing bouts—a high-risk behavior exposing the animal to recurring predation—immediately following stimulus termination (Fig. 6d). This dissociation (intact initiation but shortened maintenance) in free-moving mice underscores the critical role of NVC in sustaining defensive states.

The second life-threatening paradigm, virtual reality (VR_eagle_) eagle, allowed for head-fixed monitoring of pupil diameter in awake mice—a robust index of autonomic arousal^65^—alongside locomotor activity on a float ball (Fig. 6e). Baseline metrics (pupil diameter and locomotor activity) were comparable between genotypes (Supplementary information, Fig. S17b, c). Upon exposure to the eagle call (0–1.5 s, Figure 6f blue region) and visual dive (1.5–3 s, Figure 6f yellow region), both groups exhibited robust defensive engagement, characterized by immediate freezing and sharp pupil dilation sustained throughout the stimulus (0–3 s, Fig. 6f; Supplementary information, Fig. S17d and Video S2). This confirms that sensory processing and fear initiation remain preserved in *Grin2d*-cKO mice, consistent with observations made in the looming assay.

However, a striking divergence emerged during the “defensive offset phase”, defined as the interval from stimulus termination to the intersection of pupil diameter and locomotion speed in tracing plots for the recovery period (Fig. 6f, double-arrows lines). While littermate control mice maintained high arousal, *Grin2d*-cKO mice exhibited a rapid collapse of the defensive state (Fig. 6g). Moreover, the recovery time for pupil diameter back to baseline was reduced by ∼59% in *Grin2d*-cKO mice, consistent with the accelerated recovery time of locomotion speed observed in the looming assay (Fig. 6h). Collectively, these two independent paradigms demonstrate that NVC-deficient mice cannot physiologically sustain high-load fear states.

To establish an intermediate stress condition between the OFTs and a life-threatening predator scenario (looming and VR_eagle_), we modified the standard OFT to create a complex OFT (c-OFT). This modification involved the addition of a metal fence barrier that separated an aversive start zone (white floor) from a safe zone (black floor) (Fig. 6i).^29^ c-Fos mapping of littermate controls revealed that c-OFT paradigm indeed represented an intermediate level of BLA activation, which fell between the mild stress elicited by the OFTs and the intense stress induced by predator simulation (Fig. 6j, blue line). In contrast, c-Fos mapping in *Grin2d*-cKO mice demonstrated exaggerated neural activation under mild stress yet blunted activation under intense stress, matching their corresponding behavioral phenotypes (Fig. 6j, magenta line).

Notably, under the intermediate c-OFT condition, one-dimensional analysis of c-Fos density failed to correlate with the observed heighted defecation bouts in *Grin2d*-cKO mice (Fig. 6k). This discrepancy prompted us to extend beyond simple intensity quantification and conduct further analyses of the topological distribution patterns of c-Fos positive neurons within the BLA (Fig. 6l-m; supplementary information, S18).

We therefore decoded c-Fos topology using the structural similarity index measure (SSIM) and unbiased clustering to dissect BLA ensemble configurations (Fig. 6l, m). In littermate controls, BLA topology exhibited dynamic “stress scaling”: activation patterns evolved distinctly from minimal (Level 0) to high-threat states (Level 3), forming a precise structural hierarchy (Fig. 6l). In contrast, *Grin2d*-cKO ensembles displayed topological disorganization and a lack of hierarchy (Fig. 6m).

Mapping *Grin2d*-cKO neural SSIM signatures against the structural hierarchy of littermate control mice revealed a striking inversion of emotional scaling (Supplementary information, Fig. S18b-g): Under mild stress (OFTs), *Grin2d*-cKO patterns aberrantly clustered with high-threat control templates (Levels 2 and 3), providing a topological explanation for their hyper-reactivity (Supplementary information, Fig. S18d, e). Conversely, under high load (looming, VR_eagle_), these patterns failed to scale appropriately and instead regressed to low-stress templates (Level 1), underpinning their functional collapse (Supplementary information, Fig. S18f, g). Specifically, hierarchical clustering revealed that the *Grin2d*-cKO topological signature under c-OFT aberrantly converged with that of control mice under ARS, rather than their stage-matched controls. This pathological shift was absent in the home cage, where topologies of both genotypes remained tightly clustered (Supplementary information, Fig. S18b, c). These findings reveal that BLA NVC deficiency decouples behavior from c-Fos intensity by inducing a nuanced reconfiguration of BLA neural topology.

Collectively, these data reveal a stress load-dependent biphasic pattern (Supplementary information, Fig. S18h): NVC dysfunction precipitates topological hyper-reactivity under low stress load (OFTs) but leads to hypo-sustainability under high stress load (looming and VR_eagle_), demonstrating that functional NVC is indispensable for the topological scaling of neural networks during state transitions.

### Genetic NVC-enhancement model of *Glrb-*cKO mice

Our preceding findings established that the cation-permeable NMDA receptor (GluN2D) acts as the essential “accelerator” for NVC. Its deficiency disrupts NVC, locking the brain into a state of behavioral rigidity—manifested as hyper-reactivity under mild stress yet functional collapse under high load contexts. Guided by the fundamental principle of electrochemical balance, we hypothesized that if cation influx drives NVC, a converse mechanism mediated by anion channels might functionally restrict hemodynamic gain. Consequently, inhibiting such anionic “brakes” could theoretically potentiate NVC and confer emotional resilience.

To identify these regulators, we screened transcriptomic databases for anion channels enriched in mural cells.^56^ This unbiased screen identified *Glrb* (encoding the chloride-permeable glycine receptor subunit beta, GlyRβ) as the top candidate (Supplementary information, Fig. S19)^56^. RNAscope and immunofluorescence mapping revealed a distinct spatial segregation: *Glrb* (GlyRβ) was predominantly expressed in venous SMCs (vSMCs) (Fig. 7a–d). We validated this expression pattern using *Pdgfrβ-CreER:Glrb^f/f^* (*Glrb*-cKO) mice, which exhibited a significant reduction in mural *Glrb* (GlyRβ) levels (Fig. 7c, e, f). To determine whether these venous receptors are functionally innervated, we examined their molecular and ultrastructural organization: Molecularly, GlyRβ puncta closely colocalized with gephyrin (Fig. 7g), a scaffolding protein essential for clustering inhibitory receptors,^66^ suggesting the presence of postsynaptic-like machinery; Structurally, confocal imaging and serial electron microscopy (EM) revealed frequent direct physical contacts between neuronal axons and venular SMCs (Fig. 7h, i). Collectively, these molecular and ultrastructural features provide the morphological substrate for potential neuro-venular communication sites. Unlike the NVC deficit caused by *Grin2d* ablation, *Glrb* deletion resulted in a “super-coupled” state: *Glrb*-cKO mice exhibited significantly enhanced functional hyperemia upon whisker stimulation (Fig. 7j-m). To pinpoint the cellular origin of this enhancement, we performed a genetic dissection. First, deletion restricted to arteriolar SMCs (driven by *SMA-creER*) failed to recapitulate the phenotype (Supplementary information, Fig. S20), ruling out an arteriolar origin. Second, given that GlyRβ is selectively enriched in venular SMCs—and considering the established view that capillaries nearly lack the contractile machinery for active dilation^50^—the enhanced NVC in *Pdgfrβ* mutants is primarily attributable to venular disinhibition. Consequently, the enhanced NVC in *Glrb* mutants suggests that venular GlyRβ may physiologically constrain the magnitude of hemodynamic responses.

**Fig. 7.**
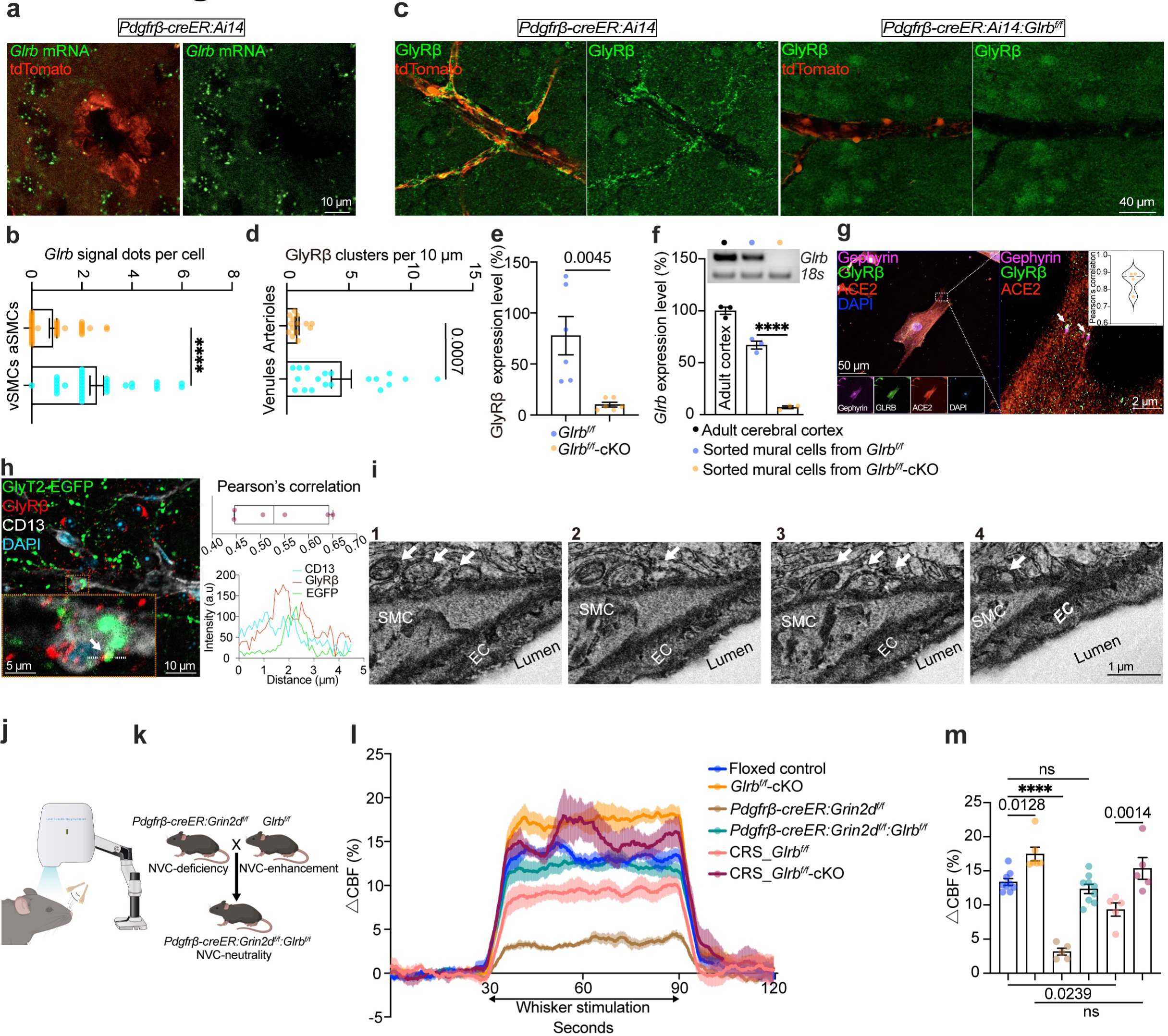
Mural cell-specific GlyRβ deficiency establishes an enhancement model for NVC. a, **b** Representative RNAscope images (**a**) and quantification (**b**) of *Glrb* mRNA expression via RNAscope in arteriolar SMCs (aSMCs) and venous SMCs (vSMCs) from brain slices. Each dot represents one dot per mural cell; *N* = 4 mice per group. Mann-Whitney test. **c–e** Representative images of GlyRβ immunofluorescence from brain slices (**c**) and quantification comparing arterioles vs. venules in littermate control mice (**d**) and venules between littermate control and *Glrb*-cKO mice (**e**). In panel (**d**), each dot represents the number of clusters per 10 μm vessel; *N* = 4 mice per group. In panel (**e**), each dot represents one analysis region of vascular mural cells from a single mouse. Mann-Whitney test (**d**) and Student’s *t*-test (**e**). **f** Electrophoretogram (top) and quantification (bottom) of *Glrb* mRNA expression in acute sorted Pdgfrβ-positive parenchymal mural cells compared with P60 medullary reticular formation lysates (as positive control). Each dot represents one mouse. One-way ANOVA followed by Tukey’s multiple comparisons test (*P* < 0.0001). **g** Co-localization of GlyRβ with gephyrin in cultured human umbilical venous SMCs (HU-vSMCs). The panel on the right shows the Pearson’s correlation coefficient analysis. Each dot represents the mean correlation coefficient of ten cells from an independent biological replicate. **h** Co-localization of GlyRβ with glycinergic axon terminals in the cortex of brain slice. The orange dotted box indicates the magnified area. Top right: Pearson’s correlation coefficient analysis. Bottom right: Fluorescence intensity linescan across the region indicated by the white arrow. Each dot represents one linescan region from one mouse. **i** Sequential electron microscopy (EM) sections of an ascending venule in the mouse cerebral cortex. White arrows indicate axonal terminals structural contacts with venous SMCs. EC: endothelial cell; SMC: smooth muscle cell. **j, k** Experimental schematics of the whisker stimulation paradigm (**j**) and the transgenic mouse mating strategy (**k**). **l, m** Functional hyperemia in the S1BF during whisker stimulation (**l**) and quantification of the corresponding plateau phase (**m**). Shaded areas represent SEM across mice in (**l**). Each dot represents one mouse. One-way ANOVA followed by Tukey’s multiple comparisons test (*P* < 0.0001). Control data (*Grin2d^f/f^* and *Pdgfrβ-creER:Grin2d^f/f^*) are shared with **Fig. S11j**. Floxed control mice group including comparable 3 *Grin2d^f/f^* mice, 3 *Glrb^f/f^* mice, and 3 *Grin2d^f/f^*:*Glrb^f/f^* mice. **** indicates *P* < 0.0001. Data are presented as mean ± SEM. See also Supplementary information, Figs. S19–S20.

Given that pharmacological vasodilation (HET0016) successfully rescued the *Grin2d*-cKO phenotype, we hypothesized that genetic removal of *Glrb* could serve as a genetic rescue strategy to counterbalance the *Grin2d* deficit. To test this genetic disinhibition hypothesis, we generated double-knockout mice (*Pdgfrβ-CreER:Grin2d^f/f^:Glrb^f/f^*; Fig. 7k). Remarkably, the concomitant deletion of *Glrb* effectively reversed the NVC deficiency triggered by *Grin2d* loss. Whisker stimulation in *Pdgfrβ-CreER:Grin2d^f/f^:Glrb^f/f^* mice elicited functional hyperemia responses comparable to littermate controls (Fig. 7l, m). This result implied that *Glrb*-cKO mimics the effect of HET0016. This finding establishes a genetic paradigm of hemodynamic rescue, demonstrating that venular disinhibition can effectively counterbalance the loss of the arteriolar accelerator. We next asked the fundamental translational question: Does this enhancement of neurovascular coupling confer behavioral resilience against stress?

### NVC enhancement confers resilience against both acute and chronic stress

Given that *Glrb* deletion unlocks a “super-coupled” hemodynamic state (Fig. 7), we first asked whether this vascular enhancement translates into immediate behavioral resilience against acute stress. We subjected mice to ARS. While littermate controls exhibited characteristic stress-induced physiological and behavioral shifts, *Glrb*-cKO mice appeared remarkably stress-buffered. They displayed significantly attenuated BLA neuronal activation (Fig. 8a, b) and blunted serum corticosterone concentrations (Fig. 8c) in ARS. Behaviorally, this physiological resilience was mirrored by stabilized defecation dynamics, with *Glrb*-cKO mice exhibiting significantly fewer defecation bouts in ARS (Fig. 8d).

**Fig. 8.**
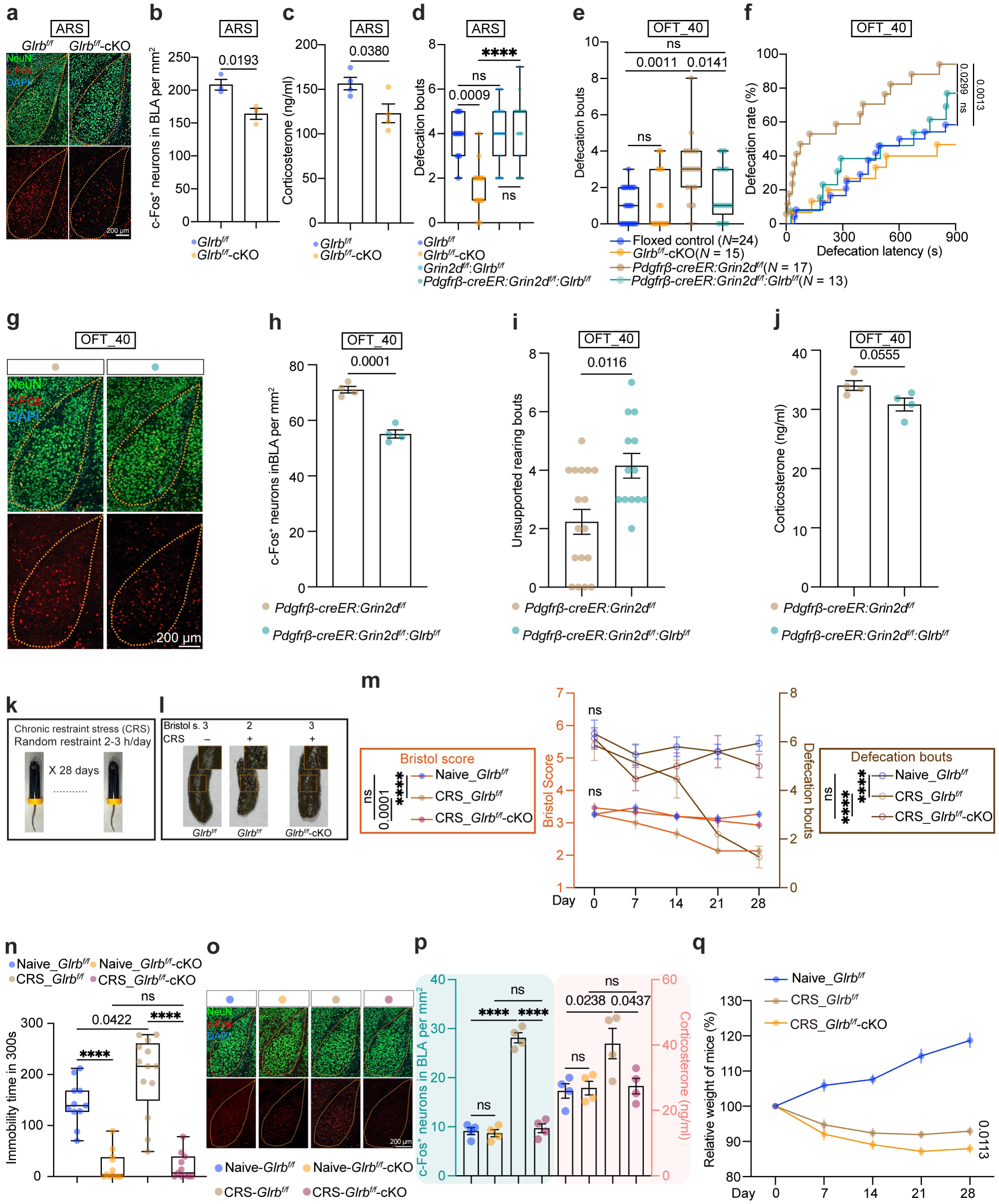
An enhancement model of NVC promotes resilience against acute and chronic emotional distress. a, **b** Representative images (**a**) and quantification (**b**) of BLA neuronal activation in ARS. Student’s *t*-test. **c** Serum corticosterone concentrations in ARS. Student’s *t*-test. **d-f** Defecation bouts in ARS (**d**) and 40 cm OFT (**e**), and defecation rate and latency in the 40 cm OFT (**f**). Kruskal-Wallis test followed by Dunn’s multiple comparisons test (*P* < 0.0001 for **d** and *P* = 0.0014 for **e**). Sample sizes (*N*) are indicated in the panel (**f**). Gehan-Breslow-Wilcoxon test for (**f**). Floxed control data in (**e** and **f**) (14 *Grin2d^f/f^* mice and 17 *Pdgfrβ-creER:Grin2d^f/f^* mice) are shared with Fig. 3b, c and **S12o**, **p**. Floxed control group in (**e**, **f**) contains comparable 14 *Grin2d^f/f^* mice, 5 *Glrb^f/f^* mice, and 5 *Grin2d^f/f^:Glrb^f/f^* mice. **g, h** Representative images (**g**) and quantification (**h**) of BLA neuronal activation in the 40 cm OFT. Student’s *t*-test. **i, j** Unsupported rearing bouts (**i**) and serum corticosterone concentrations (**j**) in the 40 cm OFT. Mann-Whitney test (**i**) and Student’s *t*-test (**j**). **k** Experimental schematic of the chronic restraint stress (CRS) paradigm. **l, m** Representative fecal appearance (**l**) and statistical analysis of Bristol scores and defecation bouts during the CRS session (**m**). All fecal scoring was performed by an independent observer blinded to the experimental groups. *N* = 15 mice per group. Two-way ANOVA on rank-transformed data followed by Sidak’s multiple comparisons test (*P* < 0.0001). Bristol s.: Bristol score. **n** Immobility time in the forced swim test (FST) after CRS modelling. One-way ANOVA followed by Tukey’s multiple comparisons test (*P* < 0.0001). **o, p** Representative images (**o**) and quantification (**p**) of BLA neuronal activation (**p left**) and serum corticosterone concentrations (**p right**) after CRS. Quantification was performed by an independent observer blinded to the experimental groups. One-way ANOVA followed by Tukey’s multiple comparisons test (*P* < 0.0001 for **p left**; *P* = 0.0400 for **p right**). **q** Body weight changes during the CRS paradigm. *N* = 15 mice per group. Student’s *t*-test. Each dot represents one mouse. **** indicates *P* < 0.0001. Data are presented as mean ± SEM. Detailed statistical data for panel (**f**) are shown in the supplementary table S1, sub-table 5. Naïve indicates mice without CRS modelling as controls. See also Supplementary information, Figs. S21–S23.

The observation that *Glrb* removal in mural cells confers stress resilience prompted us to investigate whether this NVC-enhancement model could counteract the deficits caused by *Grin2d*-cKO. We found that *Pdgfrβ-CreER:Grin2d^f/f^:Glrb^f/f^* mice effectively neutralized the core hyper-anxious phenotype observed in *Grin2d-cKO* mutants (Fig. 8d–j). Specifically, *Pdgfrβ-CreER:Grin2d^f/f^:Glrb^f/f^* mice exhibited defecation metrics comparable to NVC-intact littermate controls in ARS and 40 cm OFT (Fig. 8d-f), accompanied by attenuated BLA neural activation (Fig. 8g, h) and a significant increase in unsupported rearing bouts (Fig. 8i) in the 40 cm OFT. Collectively, the observation that *Pdgfrβ-CreER:Grin2d^f/f^:Glrb^f/f^* mice closely resemble NVC-intact mice indicates that emotional reactivity is dynamically calibrated by the precise homeostatic balance between opposing vasoconstrictive and vasodilatory signaling pathways within the BLA.

To determine whether the stress resilience observed in systemic mural cell-specific *Glrb*-cKO mice is driven by localized NVC within the BLA, we targeted the disruption of BLA NVC via local delivery of *Cald1* siRNA. In intense ARS, local BLA NVC disruption abolished the resilience-conferring effects typically seen in the *Glrb*-cKO mice, effectively reverting neural, behavioral, and hormonal stress responses to littermate control levels (Supplementary information, Fig. S21a-d). Furthermore, laser speckle contrast imaging (LSCI) revealed that whole-brain knockdown of *Cald1* via cisterna magna delivery did not alter basal CBF at the cortical surface (Supplementary information, Fig. S21e). This indicates that *Cald1* deficiency did not compromise basal vascular integrity, but specifically impairs the vascular response to neuronal activation.

While *Glrb* deletion effectively buffers the immediate impact of acute stress, a more rigorous test of neurovascular integrity is whether this protection can sustain itself against chronic adversity. We thus escalated our paradigm to chronic restraint stress (CRS, 3 hours/day for 28 days) (Fig. 8k).^19, 67^ We first validated the efficacy of the CRS model in littermate controls, which exhibited a comprehensive state of emotional and physiological exhaustion. Specifically, CRS-treated littermate control mice exhibited significantly attenuated functional hyperemia in S1BF during whisker stimulation (Fig. 7l, m).^19^ Concurrently, these mice developed a distinct constipation-like phenotype, characterized by progressive reductions in both defecation frequency and Bristol scores over the 28-day stress period (Fig. 8l, m). This was accompanied by classical anxiety-and depression-like behaviors, including significantly reduced time in the open arms of the elevated plus maze (EPM; Supplementary information, Fig. S22a), heightened neural activation in the lateral habenula (LHb; Supplementary information, Fig. S22b, c)—a critical node in depression-related circuitry^68^—and prolonged immobility in the forced swim test after the CRS modelling (FST; Fig. 8n). Furthermore, we verified that CRS exposure did not compromise BBB integrity (Supplementary information, Fig. S23). This confirms that the observed impairments in NVC and subsequent emotional and behavioral dysregulation after CRS were not confounded by non-specific vascular leakage or barrier disruption.^42, 69^

Under this sustained adversity, *Glrb*-cKO mice displayed profound resilience. Specifically, they preserved intact whisker-evoked functional hyperemia and remained resistant to the stress-induced constipation-like phenotype, maintaining normal defecation dynamics throughout the 28-day CRS modelling (Fig. 7l, m and Fig. 8l, m). Furthermore, *Glrb*-cKO mice displayed high mobility in the FST, independent of CRS exposure (Fig. 8n). At the cellular and endocrine levels, *Glrb*-cKO mice were protected from the maladaptive sensitization of stress circuits, showing no significant increase in baseline BLA neural activity and serum corticosterone concentrations compared with CRS-treated littermate controls at the end of CRS (Fig. 8o, p).

Importantly, *Glrb*-cKO mice exhibited normal locomotion and spatial preference in the 40 cm OFT (Supplementary information, Fig. S22d, e), ruling out hyperactivity or altered exploratory drive as confounding factors for the observed behavioral improvements. Finally, it is important to note that *Glrb*-cKO mediated protection may brain-region specific. While emotional and neurovascular homeostasis was preserved, *Glrb*-cKO mice still experienced stress-induced weight loss comparable to CRS-treated littermate controls (Fig. 8q). This suggested that while enhanced NVC effectively shields the brain’s emotional circuits, it did not negate the systemic physiological costs of chronic stress.

## DISCUSSION

In this study, we demonstrate that bidirectional control of NVC enables bidirectional modulations of the negative emotion states. Far from being a passive metabolic read-out of neuronal activity, NVC within the BLA–but not in the primary somatosensory cortex–operates as an indispensable active regulator of the affective negative emotion network. By combining genetic, pharmacological, and optogenetic approaches, our findings reveal that the vascular interface acts as a gatekeeper, tuning stress signals to either be amplified into anxiety-like under mild stress or dampened into resilience to strong stress. These findings redefine the BLA NVC as a proactive component of affective circuitry.

More than a century after the first measurements of regional CBF laid the foundation for modern NVC research, the field has come to a new door–what next?^70^ We knock on this door by beginning with addressing the role of BLA NVC in emotion modulation, offering a conceptual framework for exploring the regulatory capacity of NVC across functionally distinct brain regions and their respective outputs.

During optogenetic manipulations, the ∼3% hemodynamic changes we detected (Fig. 5n) is highly consistent with recent work by Hauglund et al.^57^ showing that analogous ∼3% CBV changes sufficiently drive cerebrospinal fluid flow. This concordance across two separate research groups verifies that subtle, persistent shifts in basal blood volume can be sufficient to elicit notable functional effects, for instance, elevated neural activity in the BLA (Fig. 5h, i, q, r left).

Our arteriolar optogenetic data provide genuine *in vivo* validation that the neurovascular unit (NVU) acts as a critical regulator of proximal neural activity (Fig. 5e-m). Even under the minimal-stress home cage context with negligible external stress, arteriole contraction potently activated neighboring neurons (Fig.5q, r left), defining an *in vivo* form of vasculo-neuronal coupling (VNC) that mirrors earlier *ex vivo* observations.^24^ These findings imply an implication that NVC and VNC operate as two arms of a fast, closed loop, enabling real-time, bidirectional crosstalk between vascular and neuronal compartments.

NVC is a widely recognized, evolutionarily conserved biological process.^71^ The “premature exit” observed in NVC-deficient mice under predator-like stress (Fig. 6) suggests that NVC is innately encoded to optimize the kinetics underlying the predatory fear response cycle. As a survival-critical program, this regulatory property may explain the evolutionary conservation of NVC across species.

The specific mechanism by which *Glrb* deficiency enhances NVC warrants further investigation. In mural cells, the intracellular chloride concentration (Cl-*_i_*) is typically higher than the extracellular concentration (Cl^-^ ), creating an electrochemical gradient where the opening of chloride-permeable glycine receptors leads to Cl^-^ outflow—a sharp contrast to the Cl^-^ influx observed in mature neurons.^72–75^ This efflux is predicted to depolarize mural cells and promote contraction; thus, the ablation of *Glrb* likely diminishes the baseline contractile ability of mural cells. Given that *Glrb* (GlyRβ) is selectively enriched in venular SMCs (vSMCs) (Fig. 7a–d) and that its deletion in arteriolar SMCs (aSMCs) fails to recapitulate the “super-coupled” phenotype (Supplementary information, Fig. S20a, b), we propose that *Glrb* (GlyRβ) functions as a physiological constraint on venular conductance. Upon its removal, neural inputs—such as those during whisker stimulation—elicit a significantly amplified NVC response (Fig. 7l, m).

While *Grin2d* and *Glrb* appear to exert their primary effects on distinct vascular segments—the arteriolar and venular compartments, respectively—their mutual counteraction suggests a complex hemodynamic synergy (Fig. 7l, m). One plausible interpretation is that vSMCs also possess a regulatory weight within the arteriole-capillary-venule network that is functionally comparable to that of aSMCs. These findings suggest that NVC operates as an integrated network property in the vasculature system, where enhanced venular hemodynamics functionally offsets arteriolar deficits to maintain an apparent partial equilibrium (Figures 7 and 8).

At the molecular level, we propose that GluN2D activates a signaling cascade in arteriolar SMCs—specifically, a GluN2D–Ca^2+^-bound calmodulin–caldesmon–α-SMA axis. Caldesmon typically binds to α-SMA filaments and sterically blocks their interaction with myosin, thereby suppressing cross-bridge formation.^55^ Ca^2+^-bound calmodulin, generated following Ca^2+^ entry through GluN2D-containing NMDARs, is known to relieve this inhibition.^55^ Consequently, GluN2D activation is predicted to release the caldesmon “brake”, modulating α-SMA–myosin cross-bridge cycling to increase CBF. Although this pathway warrants further direct validation, our prediction is supported by genetic epistasis analysis: *Cald1* knockdown effectively phenocopies the *Grin2d*-cKO phenotypes (Fig. 4l-q), and the lack of an additive effect when combining these two perturbations (*Grin2d*-cKO+local BLA *Cald1* KD) suggests that they potentially operate within the same axis (Fig. 4l-q).

This study has two primary limitations. First, the current lack of a venular SMCs-specific *CreER* line necessitated the use of the pan-mural *Pdgfrb-CreER* driver to perturb *Glrb* in arteriolar SMCs, pericytes, and venular SMCs. Second, we utilized male cohorts to minimize hormonal variability; future investigations involving female mice are warranted to establish the sex-dependent generalizability of these neurovascular mechanisms.

In conclusion, we identified that NVC functions as an active physiological gatekeeper rather than a passive support system. This study establishes NVC as a key target for basic research into neural circuits and mood disorders.

## MATERIALS AND METHODS

### Animal care and subject selection

All animal procedures complied with the Institutional Animal Care and Use Committee (IACUC) guidelines of the School of Life Sciences, Westlake University (Protocol 24013JJM). Mice were housed in the Westlake University Laboratory Animal Resource Centre under strictly controlled conditions: a 12-h light/dark cycle at 25 °C, with maximum six mice per individually ventilated cage, and *ad libitum* access to standard chow and water. To exclude potential confounding effects from the estrous cycle on emotional behavior, we exclusively used age-matched male C57BL/6J mice (60–80 days old at the onset of experiments).^30^

### Rigor and bias control for behavior-related assay

We implemented rigorous protocols to minimize behavioral variability arising from habituation or handling. To ensure stable baseline conditions, animals were first acclimated to the facility for at least two weeks and then habituated to the testing room environment for at least 90 min prior to each specific assay. All handling and behavioral assays were performed by a single trained experimenter who was strictly blinded to genotype and group assignments; blinding was maintained by a separate researcher who managed genotyping results. Furthermore, testing was confined to a fixed daily time window (13:00–19:00). To prevent exposure to uncontrolled olfactory or auditory cues, testing areas were strictly controlled for light, noise, and odor: apparatuses were thoroughly cleaned with 70% ethanol and ventilated between trials, and animals were transported in standardized containers. Finally, animals were randomly distributed across experimental groups from different cages to control for potential cage effects.^30, 76, 77^

### Home cage non-invasive physiological monitoring

To assess physiological parameters with minimal disturbance, mice were monitored in their home cages. Blood pressure was measured using a non-invasive tail-cuff system (Kent/Biopa), and the occupation of fat was measured using EchoMRI^TM^-100H (EchoMRI) during the fixed experimental time window (13:00–19:00); crucially, mice were habituated to the restrainer for three consecutive days prior to recording to minimize stress artifacts.

For baseline metabolic and excretory profiling, mice were housed in groups of four. Over three consecutive 3-day cycles, animals were provided with standardized quantities of food (60 g), water (340 g), and bedding (100 g). At the end of each cycle, body weight, ingestive behaviors (food and water intake), fecal output, and urination volume (derived from bedding weight gain) were recorded. For home-cage baseline assessments where individual excreta could not be definitively assigned in group-housed conditions, data were averaged per mouse within each cage. To ensure statistical rigor and avoid pseudoreplication, the cage mean was utilized as the independent experimental unit (*N* = 4 cages). To facilitate cross-group comparisons, weight data were normalized to the littermate control baseline. Specifically, the value of each sample was divided by the mean value of the control group (defined as 100%). To record defecation bouts, mice were individually housed in new home cages for 3 days of habituation. Existing fecal pellets were cleared 15 min prior to the assay, and the number of defecation bouts was recorded during the subsequent 15-min interval. Following monitoring, mice were randomly assigned to terminal procedures, including gastrointestinal inspection, retro-orbital blood collection for ELISA, blood-brain barrier (BBB) integrity assessment, and brain extraction for c-Fos immunofluorescence mapping. All assessors remained blinded to group identity throughout data collection and analysis.

### Behavioral assays

We performed all behavioral assays using a rigorous double-blind protocol. One trained experimenter conducted the assays, while a second, independent experimenter performed the video scoring and data analysis. To eliminate olfactory cues and ensure consistency, we cleaned all apparatus with 70% ethanol and fully aerated them between subjects.

### Open field tests (OFTs)

To assess emotional responses under graded stress intensities, the OFT was conducted in square arenas of three specific dimensions: standard (40 × 40 cm), intermediate (35 × 35 cm), and constricted (10 × 10 cm). Mice were placed in the center of the arena, and behavior was recorded for 15 min using an overhead camera. Unsupported rearing—defined as standing on hindlegs without contacting the arena walls—was quantified during the first 5 min. Defecation metrics (bouts, rate, and latency), running distance, and area preference were measured across the entire 15-min session. For zonal analysis, the arena floor was digitally divided into 25 equal squares (5 × 5 grid); the center zone was defined as the inner 9 squares, excluding the 16 peripheral squares adjacent to the walls.

After testing, mice were transferred to clean, individual cages. Retro-orbital blood samples for ELISA analysis were collected 5 min post-assay. Mice designated for immunofluorescence were perfused 90 min after the assay to capture peak c-Fos expression. For linear correlation analyses, behavioral and physiological data were paired from the same individual subjects.

To adhere to the 3Rs principles (Replacement, Reduction, and Refinement) and minimize animal usage, 40 cm OFT data (including whisker stimulation data) for *SMA-creER:Grin1^f/f^* mice were obtained through the retrospective re-analysis of unpublished datasets independently generated in our laboratory. This approach enhances the reproducibility of our findings, as these data were collected by a different experimenter at a distinct time point using identical experimental parameters. Due to the nature of the original experimental recordings, which were restricted to behavioral videos and whisker-evoked hemodynamic measurements, c-Fos and serum corticosterone data were not available for the *SMA-creER:Grin1^f/f^* line.

For fiber photometry and optogenetic-based OFT, only 40 cm OFT used. Tamoxifen induction 21 days prior to formal testing. For real-time NVC monitoring, *AAV2/9-hSyn-GCaMP8f* was stereotaxically injected into the BLA (AP: −1.6 mm; ML: ±3.45 mm; DV: −4.00 mm), and optic fibers were implanted immediately above the injection site (DV: −3.95 mm). Following a two-week recovery and three days of fiber-connection habituation, mice received a retro-orbital injection of Rhodamine B (Sigma, R9379; 100 μl, 5 mg/ml for formal assay) 15 min prior to the 40 cm OFT. The experimental timeline comprised three sequential recording phases: pre-test (home cage), test (40 cm OFT), and post-test (home cage).

To rigorously isolate genuine functional signals from hemodynamic and motion-related artifacts, we employed a multi-wavelength fiber photometry system (Thinkertech) for all fiber photometry assays referred published methods.^78, 79^ Fluorescence signals were acquired via 473 nm excitation for GCaMP8f (Ca^2+^-dependent) and 580 nm for Rhodamine (blood volume). A 405 nm channel was utilized as an isosbestic reference; at this wavelength, GCaMP8f fluorescence remains Ca^2+^-insensitive, serving as a precise proxy for non-specific absorption and motion artifacts that affect both the green and red emission spectra. To decontaminate the functional data, the 405 nm reference trace was independently fitted to the 473 nm and 580 nm raw signals using least-squares linear regression. These fitted components were then subtracted from their respective raw traces. This dual-correction strategy effectively neutralized common-mode noise and ensured that the reported △F/F for both GCaMP8f and Rhodamine reflects true physiological dynamics—neuronal activity and functional blood volume changes, respectively—undistorted by physical movement or non-functional optical fluctuations. These data processing and normalization protocols were applied consistently to optogenetic experiments to ensure cross-experimental comparability (details and formula shown below).

For optogenetic manipulation of vascular contractility,^43, 44, 57^ optic fibers were implanted in *SMA-creER:Ai32* (ChR2-EYFP) and control *SMA-creER:Ai47* (EYFP) mice using the coordinates described above (without *AAV2/9-hSyn-GCaMP8f* injection). Mice received tamoxifen induction 21 days prior to testing. Following habituation and Rhodamine injection, blue light pulses (473 nm) were delivered to induce optogenetic stimulation while simultaneously monitoring cerebral blood volume (CBV) via Rhodamine fluorescence (580 nm). To capture the full hemodynamic response profile, recordings were segmented into three phases: pre-stimulation baseline, optical stimulation, and post-stimulation recovery. Optogenetic stimulation cycle consisted of an 85 ms stimulation pulse followed by a 5 ms interval and a 5 ms recording window (10 Hz, 0.939 mW laser power).

As for isosbestic correction for hemodynamic artifacts, similar to BLA photometry recording, signal fluctuations at the 405 nm isosbestic point are calcium-independent and primarily reflect changes in light scattering and absorption by dynamic hemoglobin concentration changes (functional hyperemia). We applied a linear regression correction algorithm. As mentioned above, the 405 nm signal was fitted to the 473 nm signal, and the predicted hemodynamic/motion component was subtracted. This correction confirms that the CBV recording during the optogenetic stimulation represents a genuine hemodynamics.

Two distinct normalization strategies were applied to quantify CBV dynamics:

1. Static Baseline Comparison (Cross-group): To allow for cross-group comparisons of basal vascular hemodynamics, data were normalized to the mean raw value of the control group. Specifically, the raw value of each individual sample was divided by the mean raw value of the littermate control group (defined as 100%). This approach standardizes the readout, ensuring that reported values reflect specific genotypic differences in perfusion capacity relative to a physiological baseline.
2. Dynamic Response Quantification (Time-series): For time-series recordings of CBV (as well as Ca^2+^ signal recording and Laser Speckle Contrast Imaging recording), a temporal normalization strategy was applied. The baseline signal (F_baseline_) was defined as the mean value recorded during the defined pre-stimulation period (e.g., the 30-second baseline window). The relative change at each time point (F_t_) was calculated as a percentage of its own baseline using the formula:

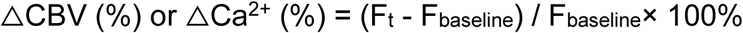

This self-normalization approach isolates the evoked hemodynamic response from background variations. This peak-aligned averaging strategy ensures that the quantification of the metabolic plateau is temporally synchronized to the individual hemodynamic kinetics of each animal.

Fourier transform plots were generated in MATLAB, with frequency and power spectral density represented on the *X* and *Y* axes, respectively. The analysis was conducted using the acquisition parameters specified in the 40 cm OFT optogenetic stimulation section. To resolve the spectral characteristics of the BLA hemodynamic response, Fast Fourier Transform (FFT)^80–82^ was applied to the CBV time series. For visualization, the low-frequency range (0–1 Hz) is presented in the representative spectral panel. For statistical quantification, the Area Under the Curve (AUC) was calculated specifically for the ultra-low frequency band (0–0.3 Hz) to represent oscillatory power across experimental groups.^83, 84^ The relevant codes can be found at the GitHub website: https://github.com/JialabEleven/Functional_hyperemia_regulates_emotion-master/tree/master/Part7_Frequency_analysis.

### Home cage optogenetic stimulation and recording

To determine if BLA vasoconstriction is sufficient to induce negative emotion in a non-aversive environment, simultaneous optogenetics and photometry were performed in the home cage. *SMA-creER:Ai32* (ChR2-EYFP) and control *SMA-creER:Ai47* (EYFP) mice received tamoxifen induction 21 days prior to testing. Optic fibers were implanted above the BLA (AP: −1.6 mm; ML: ±3.45 mm; DV: −3.95 mm) two weeks before the assay, and mice were habituated to the recording tether for three days. The experimental procedures for optogenetic stimulation in the home cage were identical to those conducted in the 40 cm OFT, with the exception of the testing environment.

On the test day, Rhodamine B (100 μL, 5 mg/mL) was administered via retro-orbital injection 15 min prior to recording. The session comprised three 15-min epochs: baseline, stimulation, and recovery. To ensure signal fidelity and prevent spectral crosstalk, an optimized interleaved illumination sequence was employed: (1) 405 nm for isosbestic control; (2) 580 nm for continuous CBV recording; and (3) 473 nm for optogenetic stimulation (delivered exclusively during the second epoch). The duty cycle consisted of an 85 ms stimulation pulse followed by a 5 ms interval and a 5 ms recording window (10 Hz, 0.939 mW laser power). To standardize baseline conditions, all existing fecal pellets were removed immediately before recording, and fresh fecal output was quantified at the end of the session. Data analysis and processing for optogenetics in the home cage is similar to that in the 40 cm OFT.

### Tube stress assay

Restraint stress was induced using modified 50 mL conical tubes (Corning, Cat# 430290).^85, 86^ To prevent hypoxia and hyperthermia, tubes were equipped with multiple ventilation apertures drilled into the cap and body, ensuring adequate airflow while strictly restricting physical movement.

For the acute restraint stress (ARS) paradigm, mice were restrained for a single 15-min session, and fecal pellet output was quantified immediately upon release. To interrogate the causal role of BLA activity in stress responses, we utilized a chemogenetic BLA neural silencing strategy. *AAV2/5-mCaMKIIa-hM4D(Gi)-EGFP-ER2-WPRE-pA* (inhibitory DREADD) or the control virus *AAV2/5-mCaMKIIa-EGFP-ER2-WPRE-pA* was bilaterally injected into the BLA (AP: −1.6 mm; ML: ±3.45 mm; DV: −4.0 mm). Following a two-week recovery period, mice were subjected to behavioral testing (including the 40 cm OFT described above). To ensure peak DREADD activation during the stressor, the agonist Compound 21 (C21; MCE, Cat# HY-100234; 0.1 mg/kg, i.p.) was administered 45 min prior to initiating restraint. Following ARS, mice were individually housed in clean cages. Retro-orbital blood samples for ELISA analysis were collected 5 min post-assay. To capture stress-induced neuronal activation, mice were perfused 90 min post-assay for c-Fos immunofluorescence mapping.

For the chronic restraint stress (CRS) paradigm, mice were subjected to daily restraint sessions for 28 consecutive days. To prevent habituation to the stressor, both the duration (randomized between 120 and 180 min) and the daily onset time (within the light phase) were varied. The physiological impact of chronic stress was assessed weekly (at the end of each 7-day block) by recording fecal output and stool consistency (Bristol Score)^87^ during a 60-min monitoring period in a clean home cage. Bristol scores analysis was conducted by an independent researcher blinded to the mouse genotypes and treatment groups. Mice designated for Elevated Plus-Maze (EPM) Test, Forced Swim Test (FST), Blood-brain-barrier (BBB) integrity assessment, ELISA inspection, and immunofluorescence analysis were sacrificed 24 hours after the final CRS session.

### Complex-open field test

To simulate an environment of high visual contrast and frustrated escape, the complex OFT (c-OFT) was conducted in a specialized chamber divided into two compartments: a high-illumination white zone (aversive) and a low-illumination black zone.^29, 88^ Crucially, these zones were separated by a barred metal partition that permitted visual contact with the safe environment but precluded physical entry. Mice were placed in the center of the white compartment at trial onset, and behavioral responses to this context of unattainable safety were continuously recorded for 20 min. Defecation bouts inspection after the assay. To capture stress-induced neuronal activation, mice were perfused 90 min post-assay for c-Fos immunofluorescence mapping.

### Looming assay

Innate defensive behaviors against aerial predators were assessed by adapting established protocols,^62^ with specific optimizations implemented to standardize threat perception. Mice were first acclimated to the arena for 10 min to stabilize baseline anxiety levels. To ensure that defensive responses (freezing or flight) were distinct from baseline immobility or thigmotaxis, a state-dependent triggering protocol was employed. Unlike continuous stimulation paradigms, the looming stimulus—a rapidly expanding overhead black disk—was manually triggered only when the mouse met specific behavioral criteria: spontaneous locomotion (speed 13–15 cm/s) and positioning within the arena center (away from the walls). Behavioral responses were recorded using a high-resolution overhead camera. Each mouse underwent a single stimulation trial to prevent habituation. Immediately following the assay, mice were housed individually in clean cages for 90 min prior to perfusion for c-Fos analysis.

### Virtual reality eagle (VR_eagle_) assay

To simulate an aerial predator attack within a highly controlled, multimodal environment, we employed a custom-built virtual reality (VR) flight simulation system. Assay design modified from previous work.^62, 63^ The apparatus consisted of an air-supported spherical treadmill (float ball) allowing unrestricted locomotion, surrounded by a panoramic display system to create an immersive visual field. Two weeks prior to testing, mice were implanted with titanium headplates. To strictly decouple restraint stress from predator-induced responses, animals underwent a rigorous daily habituation protocol (one week) to the head-fixation apparatus and the floating ball mechanism until they demonstrated stable voluntary locomotion. On the testing day, mice were head-fixed and allowed a 5-min adaptation period within a neutral visual landscape. To ensure the capture of active defensive responses, a state-dependent triggering protocol was enforced: the predator stimulus was triggered only after the mouse initiated spontaneous locomotion following the adaptation phase. The stimulus consisted of a high-fidelity eagle avatar diving rapidly from the dorsal to the proximal visual field (3 s duration), temporally synchronized with an eagle call (audiovisual integration, ∼75 dB at first 1.5 s). Pupil diameter was tracked via an infrared camera, and locomotion speed was recorded via rotary encoders sampling the spherical treadmill rotation. Data acquisition continued for 2 min post-stimulation to assess recovery kinetics. Each mouse underwent a single stimulation trial to prevent habituation. Immediately following the assay, mice were housed individually in clean cages for 90 min prior to perfusion for c-Fos analysis.

### Elevated Plus-Maze (EPM) Test

To assess anxiety-like behavior, the EPM was conducted using an apparatus elevated 50 cm above the floor, consisting of two open arms (65 × 5 cm) and two closed arms (65 × 5 cm) intersecting at a central platform.^89, 90^ At the trial onset, mice were placed in the center zone facing an open arm and allowed to explore freely for 5 min. The duration of time spent in the open arms was recorded.

### Forced Swim Test (FST)

To evaluate behavioral despair, the FST was performed in a transparent glass cylinder filled with water (25 °C) to a depth sufficient to preclude caudal support.^91^ Behavior was recorded for a 5-min session. The duration of immobility—defined as floating with the absence of active struggling, exhibiting only minimal movements necessary to maintain the head above water—was quantified. Immediately post-assay, mice were dried and returned to warmed home cages.

### Enzyme-linked immunosorbent assay (ELISA)

To assess physiological stress levels, blood samples were allowed to clot at controlled room temperature (22–25 °C) for 2 hours. Serum was isolated by centrifugation at 3,000 rpm for 20 min at 4 °C, followed by a secondary clarification spin at 8,000 rpm for 3 min to remove residual debris. Corticosterone concentrations were quantified using a commercial ELISA kit (Absin, Cat# abs554164-96T) strictly following the manufacturer’s protocol. Optical density (OD) was measured using a Cytation 5 microplate reader (Agilent, Model MS-SP104).

### Stereotaxic viral transduction

Mice were anesthetized with isoflurane and secured in a stereotaxic frame. To minimize bias, surgical procedures were performed by investigators who were blinded to the group allocations. For manipulations targeting the BLA, viral vectors were microinjected at the following coordinates relative to Bregma: AP: −1.6 mm; ML: ±3.45 mm; DV: −4.0 mm. For chemogenetic inhibition, mice received bilateral injections (150 nL per side) of the neural inhibitory DREADD virus *AAV2/5-mCaMKIIa-hM4D(Gi)-EGFP-ER2-WPRE-pA* (2.3 × 10¹² VG/mL) or the control virus *AAV2/5-mCaMKIIa-EGFP-ER2-WPRE-pA* (2.3 × 10^12^ VG/mL).^37^ For neuronal calcium recording: The calcium indicator *AAV2/9-hSyn-jGCaMP8f-WPRE-pA* (2.0 × 10^12^ VG/mL) was injected bilaterally (150 nL per side) using the same coordinates. To label glycinergic neurons in the medullary reticular formation, *rAAV2/4-Glyt2-EGFP-WPRE-hGHpolyA* (5 × 10^12^ VG/mL) was injected bilaterally (500 nL per side) at coordinates: AP: −3.5 mm; ML: ±0.84 mm; DV: −2.8 mm.^92^

Following surgery, mice were allowed to recover for at least two weeks to ensure optimal viral expression and physiological recovery prior to behavioral testing or brain collection.

### Immunofluorescence staining and microscopy

To preserve tissue integrity and antigenicity, mice were deeply anesthetized with 1% pentobarbital sodium (80 mg/kg, i.p.) and transcardially perfused with ice-cold PBS (5 min) followed by 4% paraformaldehyde (PFA) in PBS (5 min). Brains and peripheral organs were harvested, post-fixed in 4% PFA for 6 hours at 4 °C, and sectioned at 50 μm using a vibratome (Leica Histo-VT1200S). Cultured cells were washed with PBS and subsequently fixed in 4% PFA for 10 min at room temperature.

For immunolabeling, free-floating sections were permeabilized and blocked in PBS containing 1% Triton X-100 and 0.25% normal donkey serum (NDS) for 60 min at room temperature. Following three washes in PBS, sections were incubated overnight at 4 °C with primary antibodies in a buffer containing 0.1% Triton X-100. After extensive washing, sections were incubated with species-specific fluorescence-conjugated secondary antibodies (in 0.1% Triton X-100) for 2 hours at room temperature. Nuclei were counterstained with DAPI (1:1000; Beyotime, Cat# C1002). Sections were mounted using anti-fade medium (SouthernBiotech, Cat# 0100-01) and imaged using a Zeiss LSM800 confocal microscope.

To explicitly quantify the spatial distribution of proteins relative to the vasculature (e.g., GlyRβ distribution), orthogonal linescan profiling was performed. Regions of interests (ROIs) were selected where vessels appeared in longitudinal cross-section (brain slice) or appropriate places in the boundary of cultured Human Umbilical Vein Smooth Muscle Cells. Line profiles (width: 5 pixels) were drawn to capture the fluorescence intensity gradient. Besides, to validate the robustness of these spatial associations and control for selection bias, Pearson’s Correlation Coefficient (PCC) was calculated across the entire ROI using the ImageJ Coloc 2 plugin.^93^ This global metric confirmed the degree of pixel-by-pixel co-localization between the target protein and the vascular marker, independent of specific linescan trajectories.

As for cellular quantification and normalization, distinct quantification and normalization strategies were applied based on the specific biological metrics:

1. Cellular Intensity (e.g., c-Fos): The total number of NeuN-positive nuclei was identified within the defined ROI. Preliminary analysis confirmed that the density of NeuN-positive neurons was consistent across groups, validating the use of area-based normalization. Activation intensity was calculated by dividing the positive neural count by the total area of the ROI, expressed as c-Fos positive neurons per mm^2^. Additionally, to facilitate comparison with alternative quantification methodologies, the proportion of c-Fos-positive neurons relative to the total neuronal population within the BLA and LHb was also calculated. These supplementary data are provided in the supplementary information (Supplementary information, Fig. S24).
2. Relative Protein Expression (e.g., GluN2D, HIF-1α, and IgG): Mean Fluorescence Intensity (MFI) was measured specifically within the somatic ROIs of NeuN-positive neurons. To facilitate cross-group comparisons, data were normalized to the littermate control baseline. Specifically, the MFI of each sample was divided by the mean MFI of the littermate control group (defined as 100%).
3. Signal colocalization (e.g., COX-2): Background fluorescence in the COX-2 channel (546 nm) was first subtracted across the field of view. The colocalization between the COX-2 signal and the NeuN signal (488 nm) within NeuN-positive regions was then quantified using the Pearson correlation coefficient (PCC, Coloc 2 plugin in ImageJ software).

### Pharmacological interventions

To induce Cre-mediated recombination, Tamoxifen (MCE, Cat# HY-13757A) was dissolved in corn oil (Aladdin, Cat# C116023) at a concentration of 10 mg/mL. The solution was administered via oral gavage (200 μL daily). Dosing schedules were tailored to specific genotypes: *Acta2*-cKO mice were treated from postnatal day 21 (P21) to P24, whereas all other transgenic lines received treatment from P60 to P63, unless otherwise specified.

For acute pharmacological manipulations, two distinct delivery routes were employed:

1. Systemic Administration: Drugs were delivered via intraperitoneal (i.p.) injection at a standardized volume of 200 μL per mouse. Behavioral assays commenced 15 min post-infusion to target the peak efficacy window.
2. Local BLA injection: To target the BLA specifically, a stainless-steel guide cannula (RWD, Cat# 800-00245-00) was stereotaxically implanted at coordinates AP: −1.6 mm, ML: ±3.45 mm, DV: −4.0 mm. Drugs were microinfused (1 μL total volume) at a controlled rate of 250 nL/min using an automated syringe pump. HET0016 (10 μM) was microinfused locally into the BLA. Behavioral assays commenced 5 min post-infusion to target the peak efficacy window.

### Cell isolation and culture

To characterize SMC-specific molecular mechanisms, primary cultures were established by the SMCs sorted from P0 brain parenchymal tissue of *SMA-creER:Ai47* mice. Cortical parenchymal tissue was dissected following hypothermia-induced anesthesia and digested in 0.25% trypsin (Thermo Scientific, Cat# 25200072) for 15 min at 37 °C. Digestion was terminated with fetal bovine serum (FBS; Gibco, Cat# 10099). The suspension was passed through a 40 μm cell strainer (Falcon, Cat# 352340) and centrifuged at 1,600 × g for 5 min at 4 °C. Pelleted cells were resuspended in Smooth Muscle Cell Medium (SMCM; ScienCell, Cat# 1101) and plated in T25 flasks (Corning, Cat# 430639).

To purify SMC lineages, primary cultures (∼80%) were treated with Tamoxifen (5 μM in DMSO; Solarbio, Cat# D8371) to induce EGFP expression specifically in α-SMA-positive cells. Four days post-induction, EGFP-positive cells were purified using an MA-900 cell sorter (Sony, FCCF-MBR). Sorted cells were maintained in SMCM for downstream assays. The purity of SMCs isolated via this protocol has been previously validated.^94^

To capture the *in vivo* transcriptional profile of mural cells (e.g., validation of *Glrb* expression), freshly isolated cells were used rather than cultured populations. Pdgfrβ-positve cells were sorted directly from dissociated whole-brain single-cell suspensions for P60 mice using an anti-Pdgfrβ antibody and immediately lysed in TRIzol reagent to minimize transcriptional alterations. For the assay in Fig. 7g, Human Umbilical Vein Smooth Muscle Cells (HU-vSMC; ScienCell, Cat# 8020) were cultured under standard conditions recommended by the manufacturer.

### Semi-quantitative reverse transcription PCR (RT-PCR)

Total RNA was extracted from two distinct sources using TRIzol reagent (Sangon Biotec, Cat# B511311-0100): (1) Cultured Cells: Sorted EGFP-positive SMCs harvested at ∼90% confluence; and (2) Brain Tissue: Cortices or cerebella dissected from P60/P0 mice under 1% pentobarbital sodium (80 mg/kg, i.p.)-induced (P60) or hypothermia-induced (P0) anesthesia.

First-strand cDNA was synthesized (50 ng/μL) using the HiScript III 1st Strand cDNA Synthesis Kit (Vazyme, Cat# R312-02). For transcript abundance analysis, semi-quantitative PCR amplification was performed using Green Taq Mix (Vazyme, Cat# P131-AA). The reaction mixture contained 10 μL Taq Mix, 2 μL cDNA, 2 μL primer pairs (10 μM), and 6 μL RNase-free water. PCR amplicons were resolved on 2% agarose gels (BIOWEST, Cat# 111860) via electrophoresis (220 V for 30 min) and imaged using a Tanon 1600 system.

To quantify relative mRNA abundance, the mean grey value of electrophoretogram bands was quantified using ImageJ software: Band intensities were first normalized to the corresponding internal loading control (e.g., *18s*) to correct for variations in lane-to-lane loading. Then, to determine relative expression levels across groups, these normalized values were subsequently divided by the mean value of the littermate control group (defined as 100%).

### RNAscope *in situ* hybridization

To validate cell-type-specific gene expression, 15 μm horizontal brain sections were prepared using a Leica CM1950 cryostat. Genetically modified lines were utilized to map specific targets: *SMA-creER:Ai14* mice for *Grin2d* expression in arteriolar SMCs, and *Pdgfrβ-creER:Ai14* mice for *Glrb* in mural cells; *WT* mice were used for probe specificity validation (*Grin1* and *Grin2d*). Multiplex fluorescent *in situ* hybridization was performed using the RNAscope Multiplex Fluorescent Reagent Kit v2 (ACDbio, Cat# 323100) strictly adhering to the published works and manufacturer’s guidelines.^38, 89^ Specific probes included *Grin2d* (Cat# 425951-C3), *Grin1* (Cat# 431611), and *Glrb* (Cat# 1225261-C1).

Spatial quantification, including orthogonal linescan profiling and co-localization assessment, was performed identically to the immunofluorescence protocol described above. Briefly, regions of interests (ROIs) were selected where vessels appeared in longitudinal cross-section. Line profiles (width: 5 pixels) were drawn to capture the fluorescence intensity gradient. Besides, to validate the robustness of these spatial associations and control for selection bias, Pearson’s Correlation Coefficient (PCC) was calculated across the entire ROI using the ImageJ Coloc 2 plugin.

### Western blot

Protein extraction was performed on two distinct sample types: (1) Brain Tissue: Dissected regions were homogenized in ice-cold RIPA buffer (Thermo Scientific, Cat# 89901) supplemented with a protease inhibitor cocktail (CWBIO, Cat# CW22008); and (2) Cultured Cells: Cells were lysed in the same buffer at ∼90% confluence. Lysates were centrifuged at 12,000 × g for 5 min at 4 °C to remove insoluble debris. Protein concentrations were determined using a Micro BCA Protein Assay Kit (CWBIO, Cat# CW2011S). Samples were denatured by heating at 100 °C for 10 min in SDS-PAGE loading buffer. Proteins were resolved on 8% SDS-polyacrylamide gels and transferred onto 0.45 μm PVDF membranes (Millipore, Cat# IPVH00010). Following blocking with 5% non-fat milk in TBST, membranes were incubated overnight at 4 °C with primary antibodies, washed, and subsequently incubated with HRP-conjugated secondary antibodies for 45 min at room temperature. Immunoreactive bands were visualized using ECL reagents (Abbkine, Cat# BMU102-CN) and captured on a GE AI680RGB imaging system.

To quantify relative protein abundance, the mean grey value of immunoreactive bands was quantified using ImageJ software. A rigorous dual-normalization strategy was applied identically to the RT-PCR protocol: Band intensities were first normalized to the corresponding internal loading control (e.g., GAPDH) to correct for variations in lane-to-lane loading. Then, to determine relative expression levels across groups, these normalized values were subsequently divided by the mean value of the littermate control group (defined as 100%).

### siRNA administration

To achieve broad cerebrovascular knockdown; siRNA was delivered via injection into the cisterna magna following established protocols.^38^ A mixture containing 0.8 nmol siRNA and FITC-dextran (10 mg/mL; Sigma) was prepared to allow for visual tracking of the distribution (20 μL total volume, infused 1 ul/min). 10 min post-injection, successful delivery was rigorously verified by visualizing perivascular FITC fluorescence through the open scalp using a Zeiss Axio Zoom V16 microscope. Crucially, only mice exhibiting confirmed widespread perivascular perfusion were included in downstream analyses. Functional outcomes (Whisker Stimulation and 40 cm OFT) were assessed 48 hours post-injection to align with the window of peak knockdown efficiency.

For local BLA and S1BF knockdown, siRNA solution (1 nM, 150 nL) was microinjected bilaterally directly into the BLA at coordinates: AP: −1.55 mm; ML: ±3.40 mm; DV: −4.0 mm (BLA)/ DV: −0.5 mm (S1BF). The 40 cm OFT and ARS was conducted 48 hours post-injection.

### Whisker stimulation and Laser Speckle Contrast Imaging (LSCI)

To assess functional hyperemia, mice were anaesthetized with 1% sodium pentobarbital (80 mg/kg, i.p.). Assay design referred published paper.^38, 48, 60^ Following anesthesia, the head was secured in a stereotaxic frame, and the scalp was retracted to expose the skull. We stimulated whiskers longitudinally using a motor-driven brush (12 Hz, 1 min) and recorded CBF dynamics in the contralateral barrel cortex using a laser speckle contrast imaging system (RFLSI ZW/RFSLI III, RWD Life Science).^95^ Images were acquired at a resolution of 512 x 512 pixels with an exposure time 5 ms, optimized for 10 frames per second (fps) sampling.

The LSCI system provides data in “Perfusion Units (PU)”, which represent a relative blood flow index derived from the spatial blurring of laser speckle patterns. This index is related to the product of average speed and concentration of moving red blood cells in the tissue sample volume. While these raw PU values were used to compare basal perfusion levels between groups, functional hyperemia data were normalized to correct for inter-individual variations in cranial optical properties. Specifically, the CBF response was calculated as the percentage change relative to baseline:

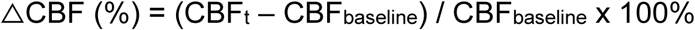

Where CBF_t_ is defined as the perfusion value of each frame; CBF_baseline_ is defined as the mean perfusion value during the 30-second pre-stimulation period.

To characterize the temporal dynamics of functional hyperemia, the peak response time was identified using a custom MATLAB algorithm (Signal Processing Toolbox). Specifically, local maxima in the △CBF time-series were detected using the findpeaks function, with empirical constraints applied to reject signal noise (minimum peak height threshold > 2.2%; minimum peak separation **>** 50 frames). The global maximum detected within the stimulation window was defined as the peak functional hyperemia. To quantify the magnitude of the sustained vascular response, we calculated the mean △CBF over a period from the identified peak latency to 90 seconds (which marks the conclusion of the whisker stimulation).

### Tissue clearing and 3D vascular reconstruction

To visualize the 3D vascular architecture, mouse brains were washed in PBS (3 × 2 hours) on a shaker (40 rpm) at room temperature. Tissue clearing was performed using a commercial kit (Nuohai, NH-CR-210701), adhering strictly to the manufacturer’s protocol. Cleared whole-brain samples were imaged using light-sheet fluorescence microscopy (LSFM; Nuohai, LS18) with a 561 nm excitation laser. For structural analysis, raw datasets were stitched using LS18, software and downsampled (0.25 ×) in ImageJ.

The datasets, comprising 1,500 sagittal slices were registered to the Allen Mouse Brain Common Coordinate Framework (CCFv3) using the BIRDS plugin. Specifically, nine reference slices were selected, and their corresponding atlas indices were manually defined to guide the registration algorithm. The automated registration for the entire dataset was then executed based on these reference points. To ensure spatial accuracy, each slice was monitored during the process, followed by randomized spot-checks post-registration. Any misalignments were resolved through manual adjustment or by re-running the registration pipeline. Based on this refined registration, Imaris Surface modules were generated for specific brain regions (eg., BLA). Vascular structures were segmented based on fluorescence signal intensity, and vascular volume was quantified using Imaris software.

### High-Fat Diet (HFD) challenge

To evaluate NVC integrity under clinically relevant metabolic stress, a diet-induced obesity model was employed. Mice were randomized into HFD (SYSE, Cat# PD6001; 60% kcal from fat) or normal fat diet (NFD) control groups for four weeks. Body weight and food intake were monitored weekly. Physiological impact was assessed by quantifying home-cage fecal output as described above. Following the 4-week modelling, mice were subjected to the 40 cm OFT, and whisker stimulation assays to assess emotional and hemodynamic function, respectively. For ELISA analysis, serum was collected 5 min after the 40 cm OFT. For c-Fos immunofluorescence (IF) staining, brain tissues were harvested 90 min post- 40 cm OFT.

### Immunoprecipitation (IP) assay

To map the GluN2D interactome, cultured SMCs sorted from P0 *SMA-creER:Ai14* mice brain parenchymal tissue (via tdTomato-based sorting) were collected and subsequently expanded in SMCM. At ∼90% confluence, cells were transduced with lentivirus encoding Grin2d-3XFlag (*Lenti-EF1-EGFP-CMV-Grin2d-3XFlag-WPRE*, 5 × 10^12^ VG/mL). One-week post-transduction, infected cells (EGFP-tdTomato double positive) were purified using an MA-900 sorter and expanded in 10 cm dishes. Upon reaching 90% confluence, cells were lysed in RIPA buffer. Lysates were incubated with Anti-GluN2D or IgG control antibodies conjugated to Protein G Magnetic Beads (Epizyme, Cat# YJ002) to pull down the protein complex. Eluates were analyzed via Western blot and mass spectrometry.

### Blood-Brain Barrier (BBB) permeability assay

To evaluate BBB integrity, the Evans Blue dye extravasation assay was performed.^96^ Mice were injected retro-orbitally with 2% Evans Blue (Merck, Cat# E2129; dissolved in saline; 4 μL/g). 45 min post-injection, mice were anesthetized, and whole brains were harvested. Macroscopic dye leakage was first assessed qualitatively using a Zeiss Axio Zoom V16 microscope. For quantitative assessment, brains were weighed and homogenized in 1 mL of 50% trichloroacetic acid (Merck, Cat# 190780). Following centrifugation at 10,000 × g for 20 min at 4 °C, the supernatant was collected, and Evans Blue concentration was quantified by measuring absorbance at 620 nm. To detect microscopic leakage, immunofluorescence staining for Immunoglobulin G (IgG) was performed as described in the Immunofluorescence Staining section. We utilized brain sections from mice subjected to non-perfusion as positive controls—defining their mean fluorescence intensity as the 100% benchmark—to quantify the IgG intensity.

### Colorectal endoscopy

To assess stress-induced gastrointestinal pathology, representative images of the distal colon were captured using the ENDOQ colonoscopy system. To ensure unbiased evaluation, a gastroenterologist blinded to the experimental groups scored mucosal morphology and signs of inflammation based on the endoscopic images.

### Hematoxylin and Eosin (H&E) staining

For histological analysis, mice were anesthetized and perfused as described above. Harvested tissues were post-fixed in 4% PFA for 6 hours at 4 °C. Following standard paraffin embedding, 8 μm sections were cut (Histo. Microtome, Leica FC-LRM), and H&E staining was performed to visualize gross morphological structures.

### Topological characterization of BLA activation patterns

To objectively classify emotional states based on the spatial configuration of neuronal ensembles, a topology-based estimation framework was developed, integrating the Structural Similarity Index (SSIM) with unsupervised hierarchical clustering.

To transition from discrete cellular signals to a continuous representation of regional activation, raw c-Fos immunofluorescence staining images were preprocessed in MATLAB. Images were converted to greyscale, inverted, and normalized to a [0, 1] intensity scale. A multi-scale Gaussian smoothing protocol (σ1 = 6 pixels, σ2 = 12 pixels) was applied to generate a continuous spatial intensity map representing the macro-scale topography of neuronal activation within the BLA. This step effectively minimizes high-frequency noise from individual cell variations while preserving the global spatial structure.

To quantify the topological similarity between experimental conditions, pairwise SSIM values were computed between the activation intensity maps. Unlike traditional metrics that rely on absolute cell counts or pixel-by-pixel subtraction, SSIM assesses the preservation of structural information by integrating comparisons of luminance (*l*), contrast (*c*), and structure (*s*). The similarity index between image *x* and *y* is calculated as:

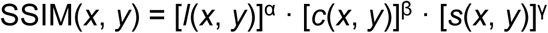

This metric provides a robust measure of spatial fidelity, converting image-based topographical correlations into a numerical similarity matrix.

The resulting SSIM similarity matrix was utilized for cluster analysis in R. We defined the topological distance (*D*) between any two conditions as *D* = 1 - SSIM. Based on this distance matrix, unsupervised hierarchical clustering was performed using Ward’s minimum variance method (method = “ward.D2”). This algorithm minimizes total within-cluster variance to construct a dendrogram, effectively grouping behavioral contexts into distinct clusters based purely on topological proximity. These naturally emerging clusters were then mapped to discrete stress Levels (Levels 0–3), defining the subject’s state based on the intrinsic topology of their neuronal response rather than arbitrary thresholds.

To ensure rigor, a double-blind workflow was implemented: image processing and algorithmic analysis were conducted by two independent analysts, distinct from the personnel performing wet-lab experiments. Statistical analyses were performed using GraphPad Prism (v9.4.1) and custom MATLAB scripts. All source code is available at website in the Github shown below: https://github.com/JialabEleven/Functional_hyperemia_regulates_emotion-master.

### Serial Block-Face Scanning Electron Microscopy (SBF-SEM)

To visualize the ultrastructure of the neurovascular interface, we prepared specimens consistent with our previous protocols.^38^ Briefly, we identified ascending venules using X-ray microscopy (Zeiss Xradia 520 Versa) at 4 × magnification (80 kV). We mounted the specimen on an aluminum pin using conductive epoxy (Chemtronics, CW2400) and trimmed it into a 600 × 600 × 700 μm^3^ block using a glass knife and a diamond trimming knife (DiATOME, Trim 90). Prior to imaging, we sputter-coated the sample with gold–palladium (Leica EM ACE200). We acquired images using a Carl Zeiss Gemini 300 scanning electron microscope equipped with a Gatan 3View ultramicrotome and a BSE detector. Imaging parameters were set to 2 kV acceleration voltage under high vacuum (∼1.8 × 10^-3^ mBar) with focal charge compensation. We performed serial sectioning at a thickness of 50–75 nm with a cutting speed of 0.1 mm/s.

### Stroke modelling

To validate the absence of hypoxic injury in our optogenetic model, we generated a positive control for hypoxia using the Stroke Induced by Magnetic ParticLEs (SIMPLE) method.^58–60^ Briefly, we injected magnetic microparticles into the bloodstream and positioned a magnet over the middle cerebral artery (MCA) to induce targeted occlusion. This permanent ischemia triggers robust upregulation of Hypoxia-inducible factor 1-alpha (HIF-1α) within 24 hours. We utilized brain sections from mice subjected to the SIMPLE stroke model as positive controls—defining their mean fluorescence intensity in the stroke infarct area as the 100% benchmark—to quantify HIF-1α expression.

### Software, code, and data availability

Fluorescence images were processed using Zeiss Zen Lite (v3.4) and ImageJ (v1.53u). For fiber photometry, signal demodulation and linear regression correction were performed using the system’s built-in proprietary software (Thinkertech). Whisker stimulation data were acquired and analyzed using the RWD LSCI software (v5.0). Trajectory tracking for the OFTs, complex OFT, and EPM was performed using the Noldus EthoVision XT system (v18), which was also utilized to generate locomotion heatmaps. For the Looming and VR_eagle_ assays, tracking was conducted using custom scripts (Bayonne). Physiological and Topology Analysis: Custom MATLAB scripts were developed to analyze CBF peak kinetics, map aggregate defecation coordinates, and generate neural activation heatmaps for the topology-based estimation framework. KEGG pathway enrichment analysis and SSIM-based hierarchical clustering were executed using custom R algorithms. Statistical analyses were conducted using GraphPad Prism (v9.4.1) and MATLAB. Figures were assembled using Adobe Illustrator 2024 (v28.3). Outliers were identified and excluded using the Interquartile Range (IQR) method (values falling outside the range of Q1 − 1.5 × IQR to Q3 + 1.5 × IQR).^97^ Data are expressed as mean ± standard error of the mean (SEM). The normality of data distribution was assessed using the Shapiro-Wilk test. Specific statistical tests employed for each experiment are detailed in the corresponding figure legends. Exact *P* values are reported to four decimal places. A *P* value < 0.05 was considered statistically significant. Statistical analyses were performed using GraphPad Prism (v9.4.1). A comprehensive data resource checklist, detailing figure attributions and sample sources, is provided in Supplementary Table S2 for reference. Furthermore, to ensure maximum transparency and statistical robustness, the percentage of c-Fos positive neurons in the BLA across all experimental cohorts is quantified and presented in Supplementary Figure S24. All custom MATLAB and R code generated for the topology-based estimation framework and hemodynamic analysis is available at the GitHub website for free searching and use: https://github.com/JialabEleven/Functional_hyperemia_regulates_emotion-master.

## ACKNOWLEDGMENTS

We sincerely thank Prof. Bo Li for his invaluable advice and for editing the manuscript. We are grateful to Prof. Hailan Hu for providing the *Ai32* mice, and to Prof. Yanmei Tao and Dr. Xu Hu for sharing the *Nestin-Cre* and *Emx1-Cre* lines. We also thank Xiaoxuan Zhang for providing stroke-modeling mice; Xuan Li from Prof. Bo Li’s lab for guidance on fiber photometry in the amygdala; and Wuyang Zhang for assisting with data assessment during the revision. We thank Dr. Zhu Zhu for providing critical discussion during the revision. We thank rotation/visiting students Jiaxing Tang, Zeyu Meng, Jiaqi Tu, and Yiran Ge for double-bind analysis of behavioral data. furthermore, we thank the following staff at Westlake University for their technical support: Yi Zhu for sharing psychological insights that facilitated our data interpretation; Yajie Liu at the Laboratory Animal Resource Center (LARC) for guidance on behavioral assays; Dr. Jiongfang Xie at the Microscopy Core Facility for imaging assistance; Dr. Mingzhu Fan at the Mass Spectrometry & Metabolomics Core Facility (MSMCF) for mass spectrometry data collection; and Yanan Tang at the Flow Cytometry Core Facility (FCCF) for assistance with cell sorting.

This work was supported by Westlake Laboratory of Life Sciences and Biomedicine (Project No. 202309002), the National Natural Science Foundation of China (project No. 32571142 and 32170961), State Key Laboratory of Gene Expression (Grant No. 2025ZY01117), Hangzhou Leading Innovation Team (TD2024001), the Joint Fund of the Zhejiang Provincial Natural Science Foundation of China (LHZQN25H160001), the “Pioneer” and “Leading Goose” R&D Program of Zhejiang (grant 2024SSYS0031), and the partially supported from the Construction Fund of Key Medical Disciplines of Hangzhou (2025HZZD04) from H.W.

## AUTHOR CONTRIBUTIONS

J.-M.J. conceived the project, designed the experiments, and supervised this study. J.Y.R. designed and performed key experiments, obtained and organized the majority of original data, and wrote the manuscript draft. X.H.Q. designed the complex OFT, Bristol score assessments, gut health inspection, and S1BF NVC control assays; performed the cluster analysis of topological SSIM data and conducted *Cald1* cisterna magna injections; participated in the 40 cm OFT, looming and VR_eagle_ assays, BBB integrity inspections, and blood sample collection; and provided critical suggestions during the revision. H.Q.X. spearheaded the 3D reconstruction of BLA arterioles by registering tissue-cleared images to the mouse brain atlas; developed a fiber photometry prototype for BLA CBV recording; participated in the Looming and VR_eagle_ assays and blood pressure measurements; and contributed to the assessment of stress-hormone levels. Y.Y.Z. developed the MATLAB-based computational pipeline for functional hyperemia peak detection, mass spectrometry data preprocessing, BLA neural topological analysis, and Fast Fourier Transform (FFT) processing. D.D.Z. provided the raw dataset of the cleared brain of the *SMA-creER:Ai14* mouse, primer pairs of *Grin1*, raw data of the serial electron microscopy, and *SMA-creER:Grin1^f/f^* 40 cm OFT video raw data. W.T.W. contributed to the writing of MATLAB code with Y.Y.Z.. B.R.Z. participated in the double-blind genotyping management of mice and cell sorting. D.P.L. helped the mouse brain section after 35 cm OFT. Y.X.N illustrated the KEGG maps.

## Supplementary information

### Supplementary figure legends

**Fig. S1.**
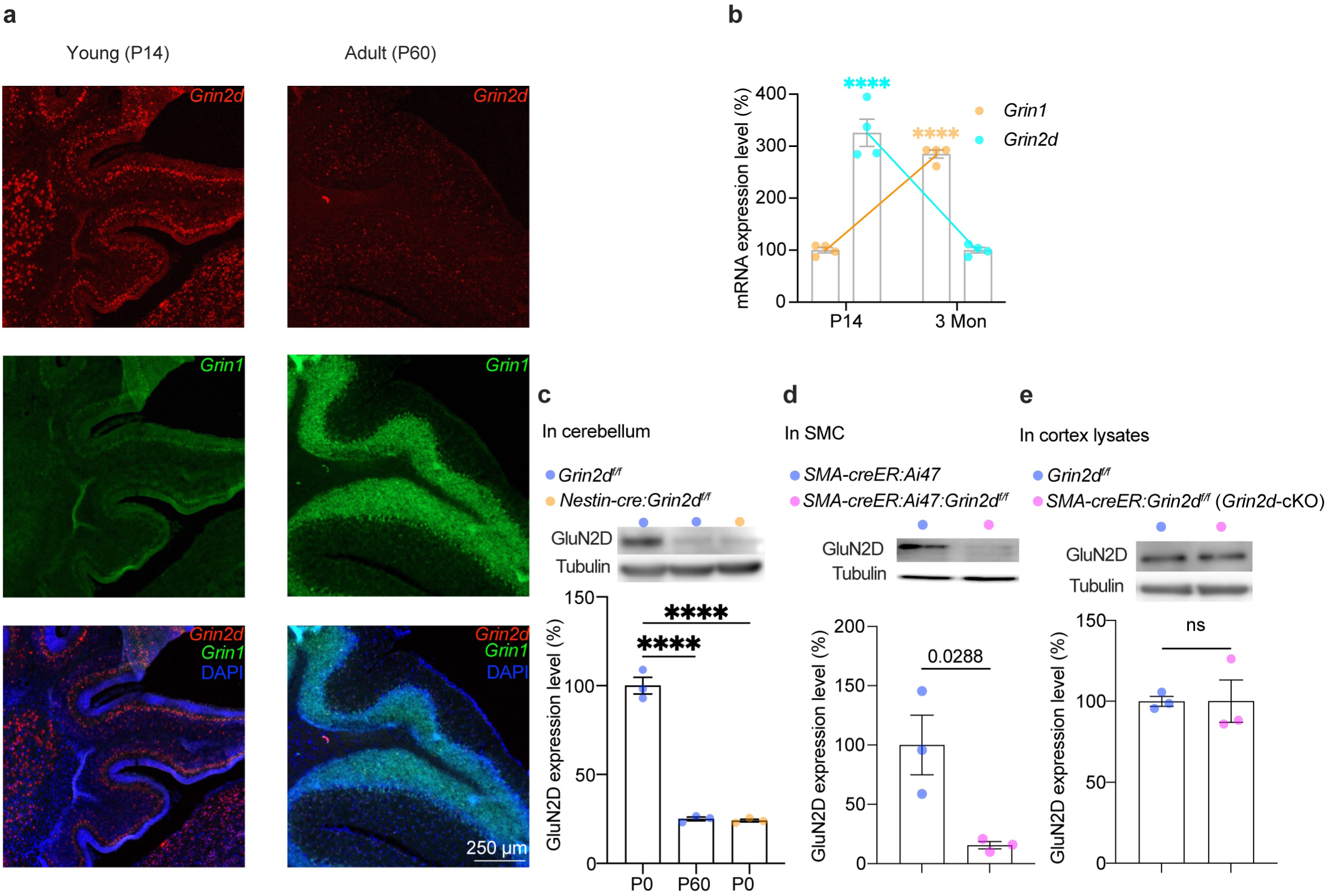
Validation of successful GluN2D conditional Knock-out. a, **b** Representative images (**a**) and quantification (**b**) of *Grin1* and *Grin2d* mRNA expression via RNAscope in the cerebellum. Each dot represents one region of interest (ROI) from a single brain slice per mouse. Student’s *t*-test. **c–e** Representative immunoblot (top) and quantification (bottom) of GluN2D protein levels in P0/P60 cerebellum lysates (**c**), cultured SMCs sorted from P0 parenchymal tissue (**d**), and P60 cortex lysates(**e**). Each dot represents one mouse. One-way ANOVA followed by Tukey’s multiple comparisons test for (**c,** *P* < 0.0001), and Student’s *t*-test for (**d**, **e**). **** indicates *P* < 0.0001. Data are presented as mean ± SEM.

**Fig. S2.**
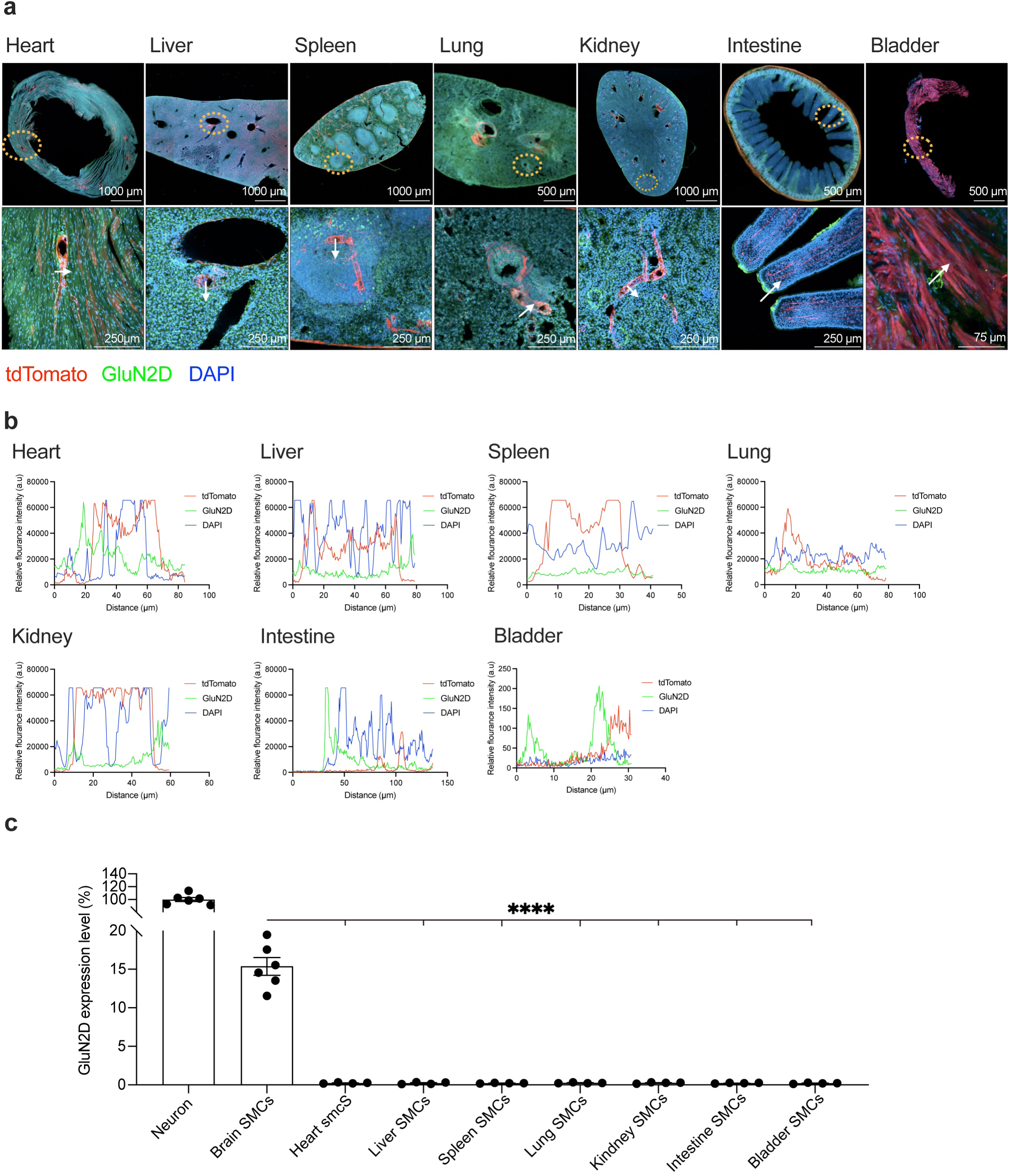
Validation of GluN2D expression in peripheral organs’ SMCs. **a** Representative low-magnification images of tissue sections from the heart, liver, spleen, lung, kidney, small intestine, and bladder of *SMA-creER:Ai14* mice (top). Yellow dotted boxes indicate the magnified areas shown below. White arrows indicate ROIs for fluorescence intensity linescan analysis. **b** Fluorescence intensity distribution (linescan) across the ROIs indicated by white arrows in (**a**). **c** Quantification of GluN2D immunofluorescence (IF) staining intensity of GluN2D in SMCs across tissues. Each dot represents one mouse. One-way ANOVA followed by Tukey’s multiple comparisons test (*P* < 0.0001). **** indicates *P* < 0.0001. Data are presented as mean ± SEM.

**Fig. S3.**
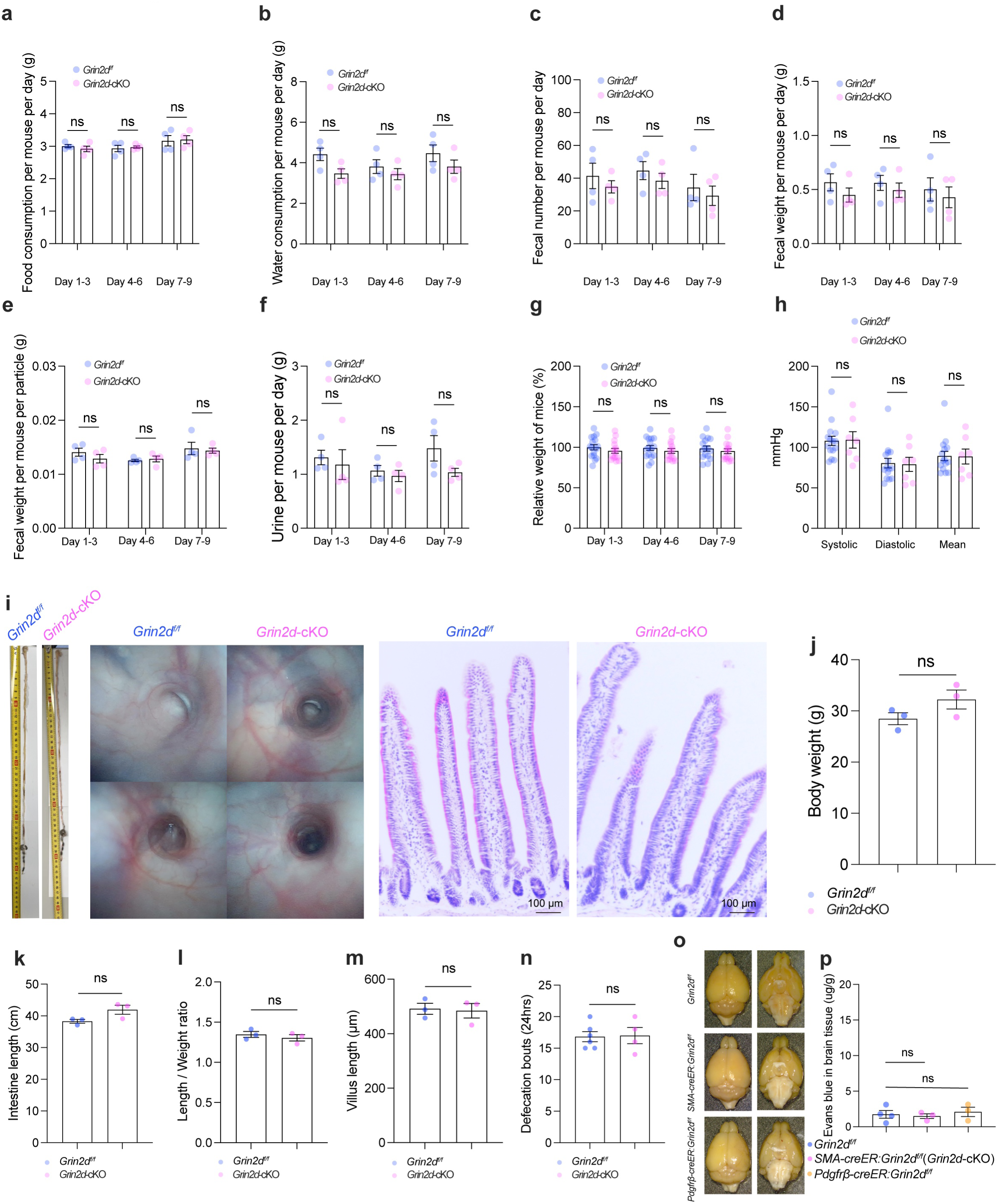
Verification of mouse metabolic status, gut health, and blood-brain barrier integrity. a–h. Quantification of food consumption (**a**), water intake (**b**), number of fecal pellets (**c**), daily fecal weight (**d**), weight per pellet (**e**), urine output (**f**), relative body weight changes (**g**), and blood pressure (**h**) measured in the home cage. For (**a–f**), each dot represents the average value per cage (4 mice per cage). For (**g, h**), each dot represents an individual mouse. Mann-Whitney test (**c**) and Student’s *t*-test (except **c**). **i** Representative images of intestinal morphology (appearance and length) and H&E staining of intestinal sections. **j–n** Quantification of data from (**i**), including mice body weight (**j**), intestinal length (**k**), intestinal length-to-body weight ratio (**l**), villus length (**m**), and total 24-hour defecation count (**n**) for mice in the home cage. Each dot represents one mouse. Student’s *t*-test. **o, p** Representative images of the brain following retro-orbital Evans Blue injection (**o**) and the corresponding quantification (**p**). Each dot represents one mouse. One-way ANOVA followed by Tukey’s multiple comparisons test (*P* = 0.7654). Data are presented as mean ± SEM.

**Fig. S4.**
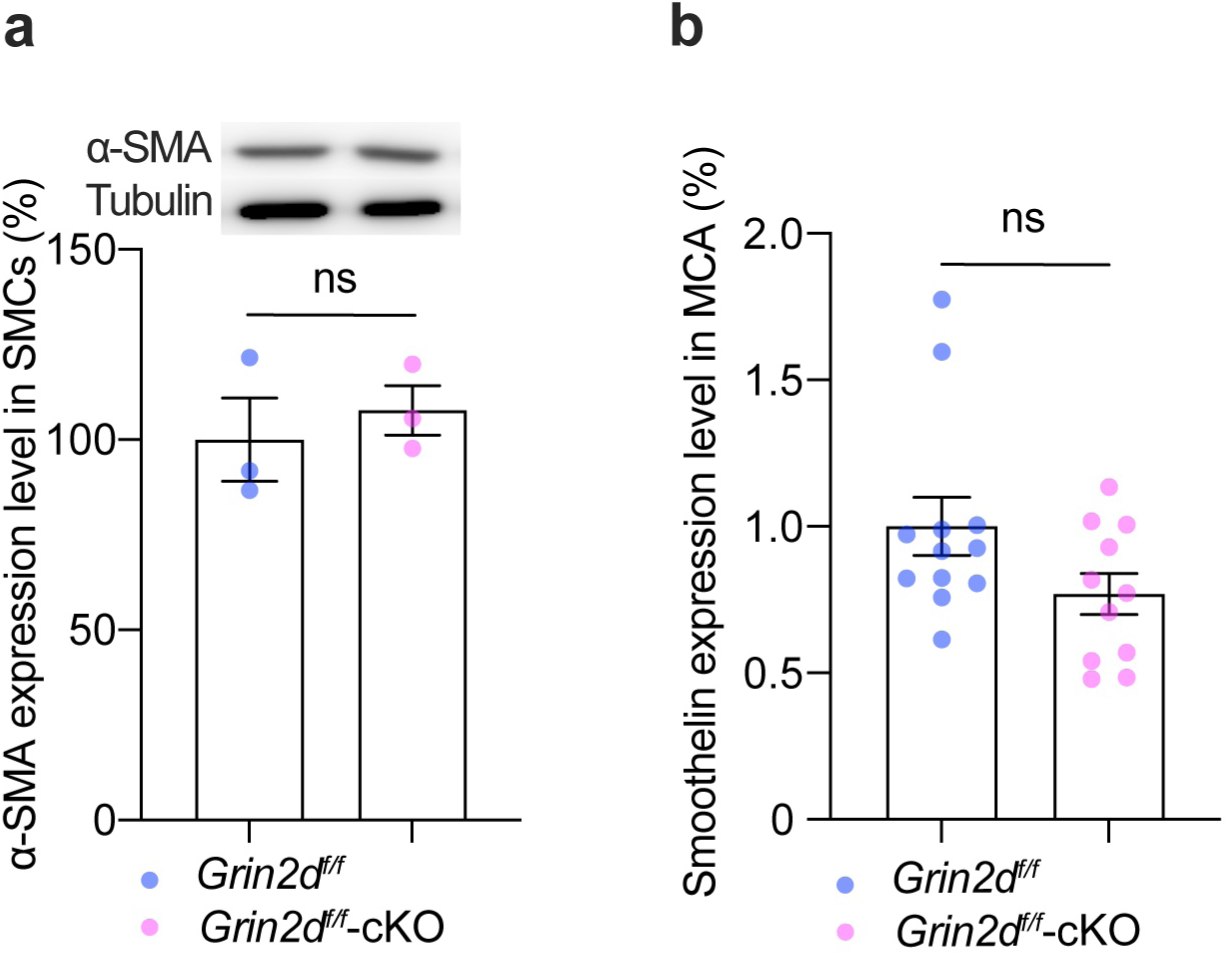
Validation of SMC marker proteins in *Grin2d*-cKO mice. **a** Representative immunoblot (top) and quantification (bottom) of α-SMA expression. Each dot represents one mouse. Student’s *t*-test. **b** Quantification of Smoothelin immunofluorescence (IF) staining in the middle cerebral artery (MCA), corresponding to the representative images in Fig. 2d. Each dot represents one analysis region of MCA from a single mouse. *N* = 4 mice per group. Student’s *t*-test. Data are presented as mean ± SEM.

**Fig. S5.**
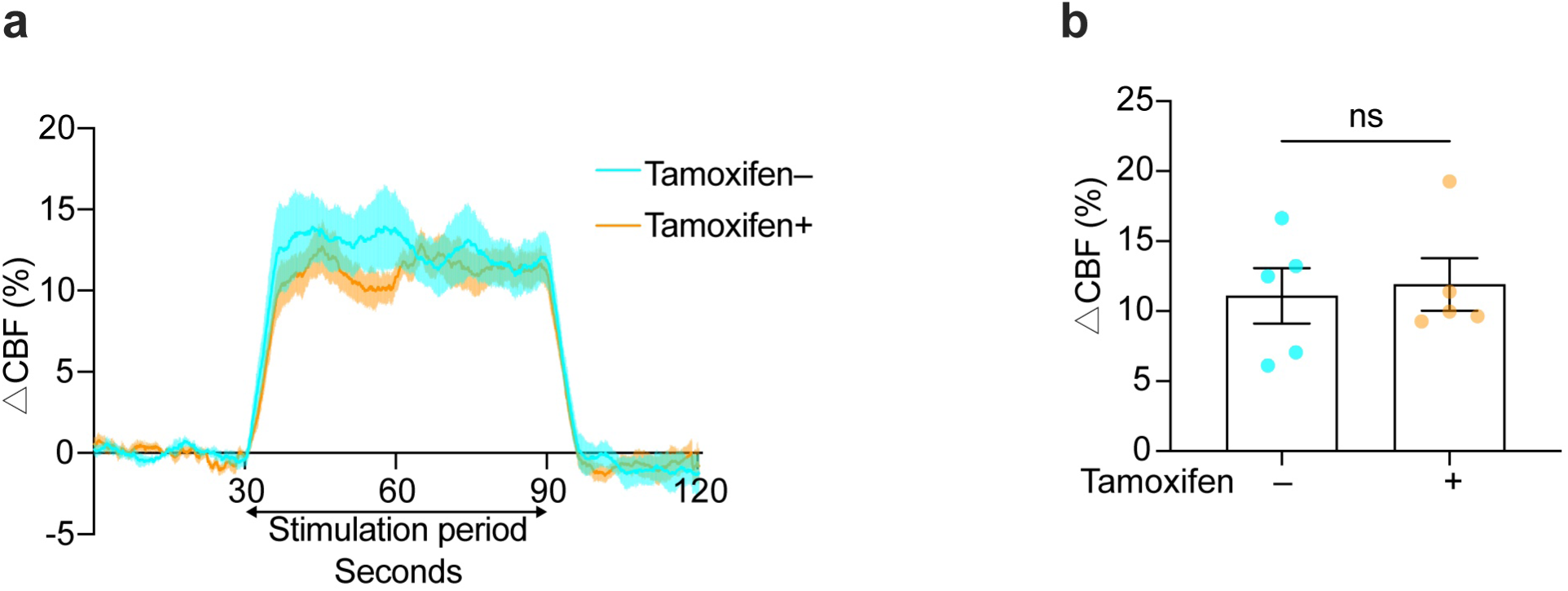
The effect of tamoxifen on NVC. **a** Whisker-evoked functional hyperemia in the S1BF of littermate control mice treated with tamoxifen. Shaded areas represent SEM across mice. **b** Quantification of the functional hyperemia plateau phase from the data in (**a**). Each dot represents one mouse. Student’s *t*-test. Data are presented as mean ± SEM.

**Fig. S6.**
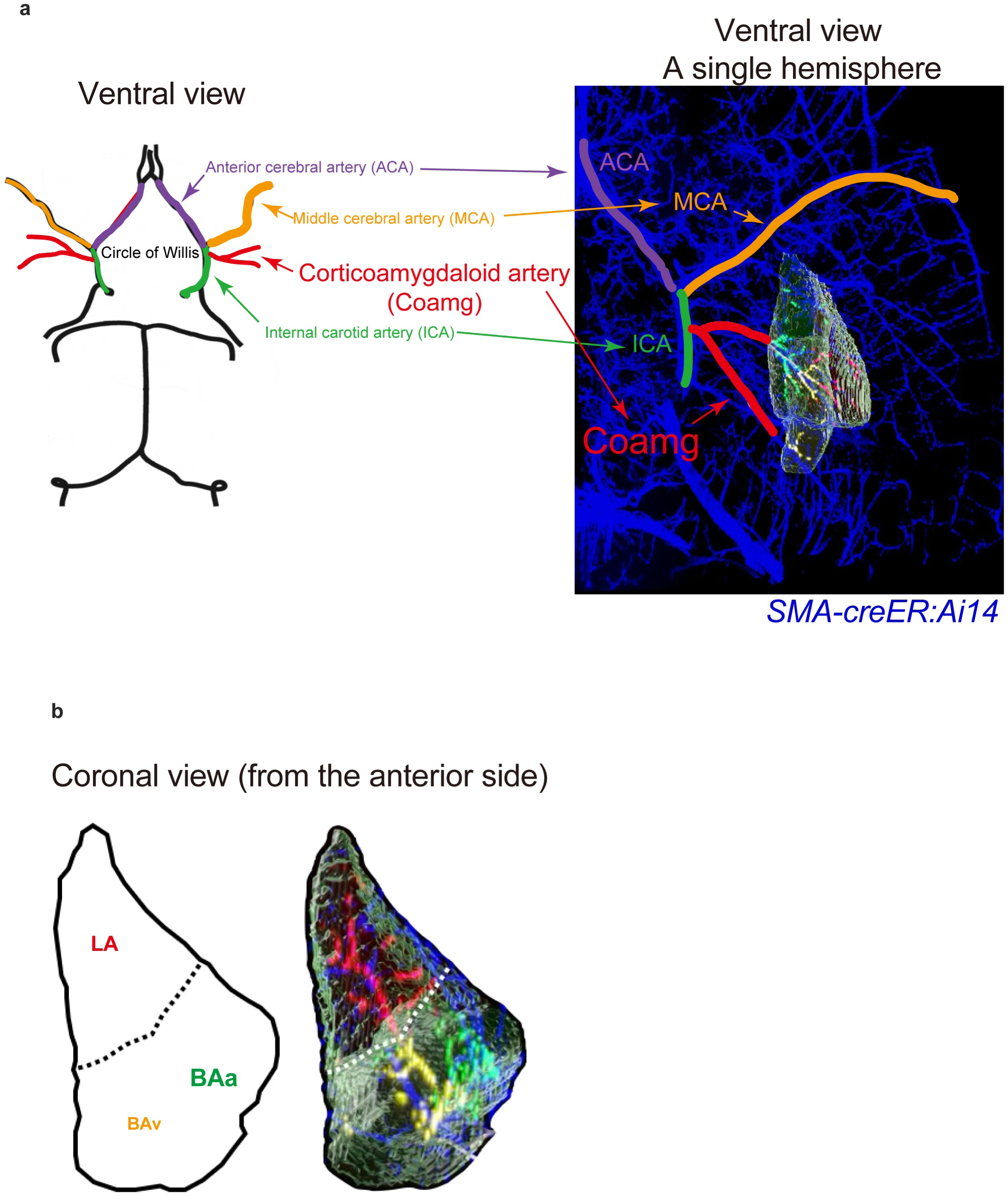
3D-Reconstruction of BLA arteriolar topology through correlative integration of the Allen Brain Atlas with tissue-cleared brains from *SMA-creER:Ai14* mouse. **a** Ventral view of the mouse BLA. Schematic diagram (left) was adapted from *The Mouse Nervous System* (p. 460, Fig. 14.1), and 3D-reconstruciton (right) was generated in Imaris using the Allen Mouse Brain Common Coordinate Framework plugin. BLA is denoted translucent grey. Blue denotes arteriolar SMCs. Colored lines represent distinct arteriolar segments. **b** Coronal view of the mouse BLA along with arterioles inside. Colored lines indicate arterioles in basal amygdala anterior part (BAa), basal amygdala ventral part (BAv), and lateral amygdala (LA).

**Fig. S7.**
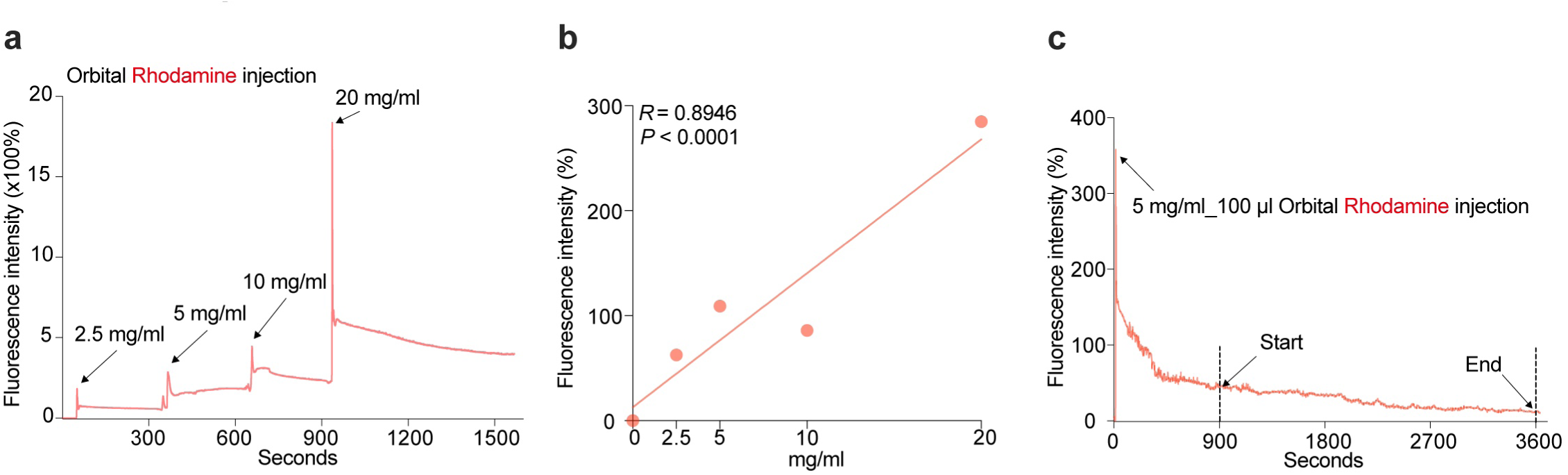
Calibration and validation of plasma Rhodamine fluorescence intensity. **a** Real-time fluorescence intensity measurement following retro-orbital injection of various Rhodamine concentrations. **b** Linear correlation between Rhodamine concentration and fluorescence intensity in (**a**). Each dot represents the average intensity from 120–150 s post-injection for a specific concentration. **c** Real-time fluorescence intensity measurements following retro-orbital injection of 5 mg/mL Rhodamine, showing signal stability from 900–3600 s post-injection.

**Fig. S8.**
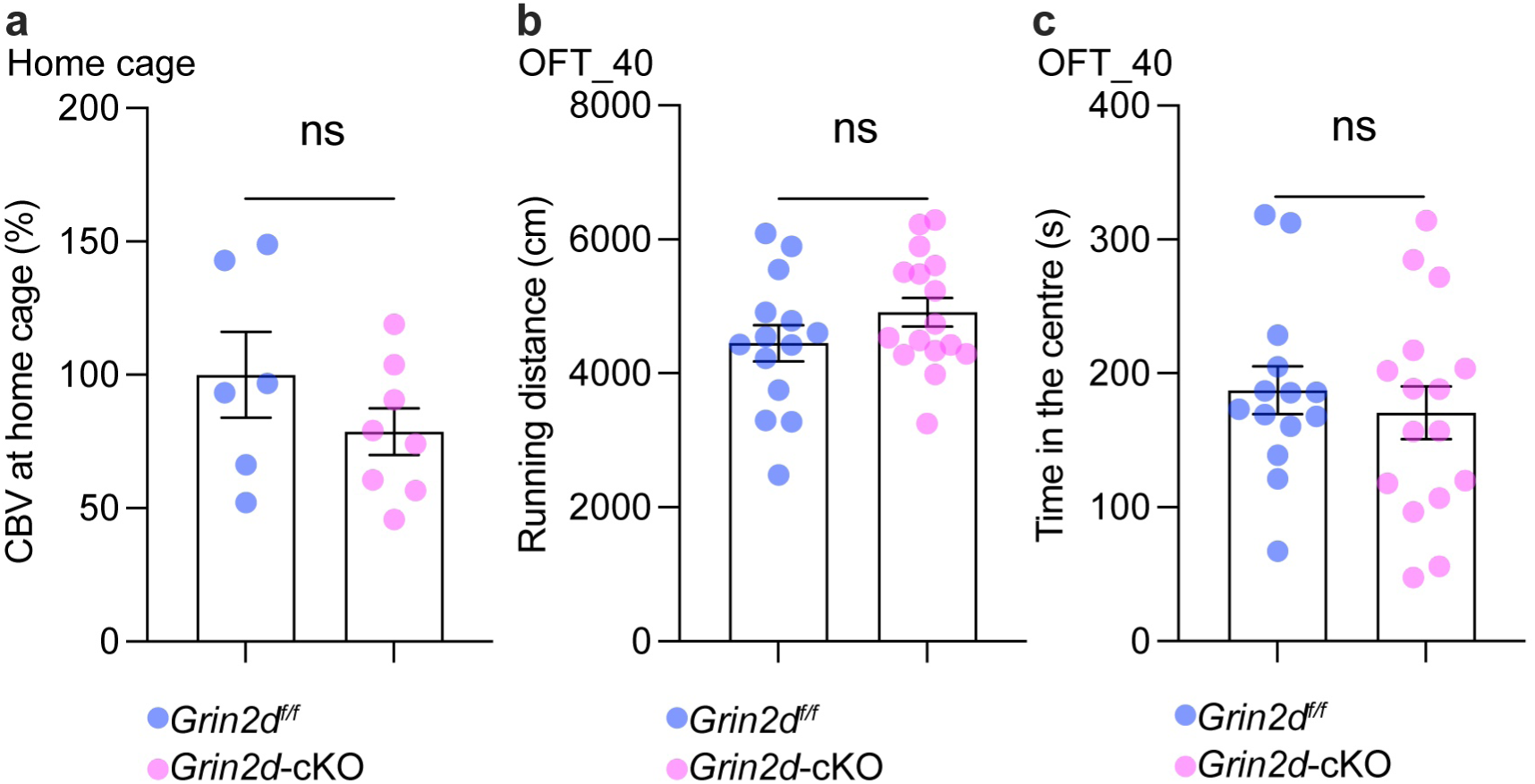
Baseline CBV and OFT performance in *Grin2d*-cKO mice. **a** Baseline CBV in the BLA during the home cage stage. Each dot represents one mouse. Student’s *t*-test. **b, c** Running distance (**b**) and time spent in the center zone (**c**) in the 40 cm OFT. Each dot represents one mouse. Student’s *t*-test. Data are presented as mean ± SEM.

**Fig. S9.**
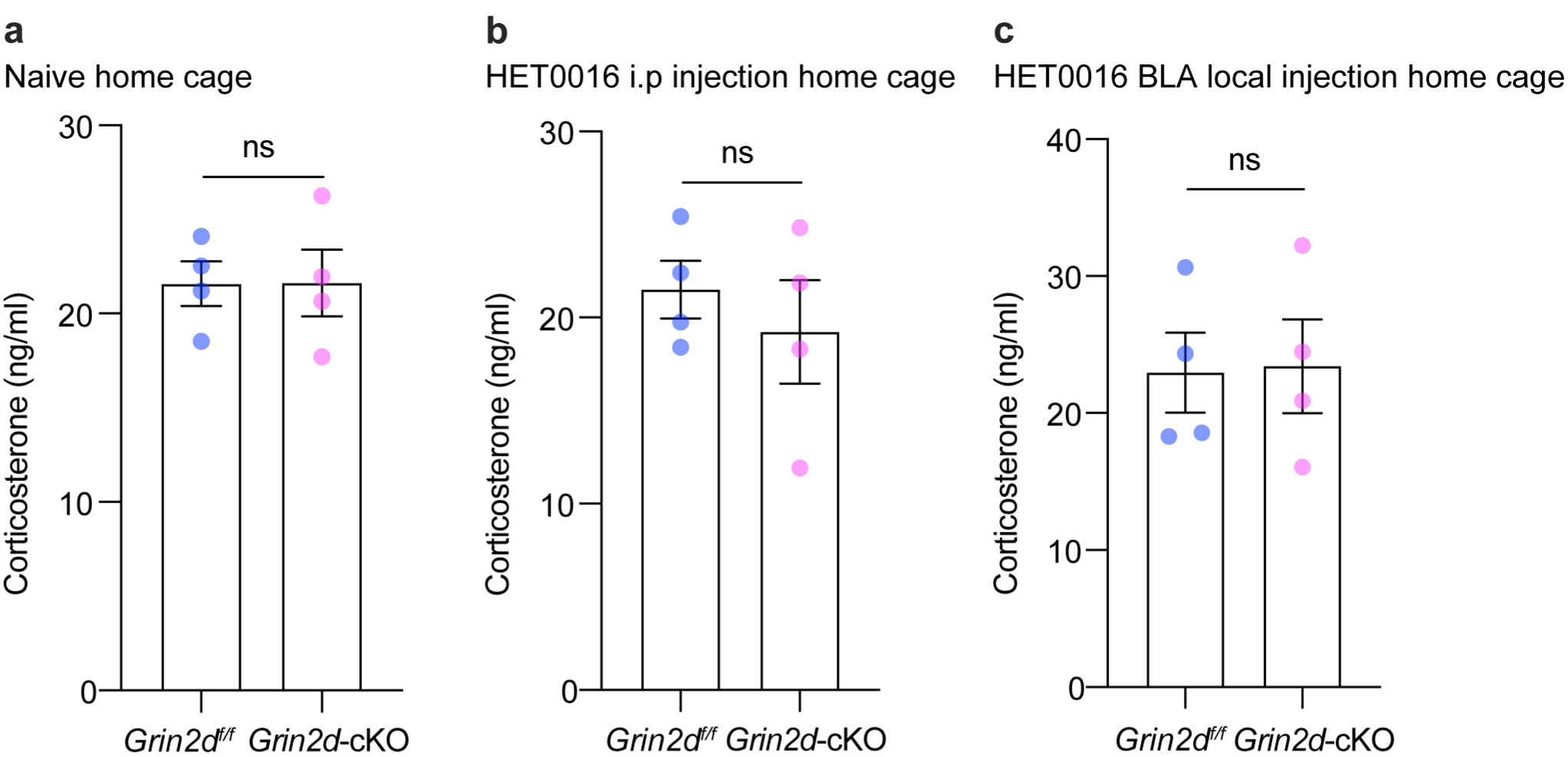
Serum corticosterone concentrations under baseline conditions. a–c. Serum corticosterone concentrations in mice at the home cage condition (**a**), following i.p. injection of HET0016 (**b**), and following local BLA microinjection of HET0016 (**c**). Each dot represents one mouse. Student’s *t*-test. Data are presented as mean ± SEM.

**Fig. S10.**
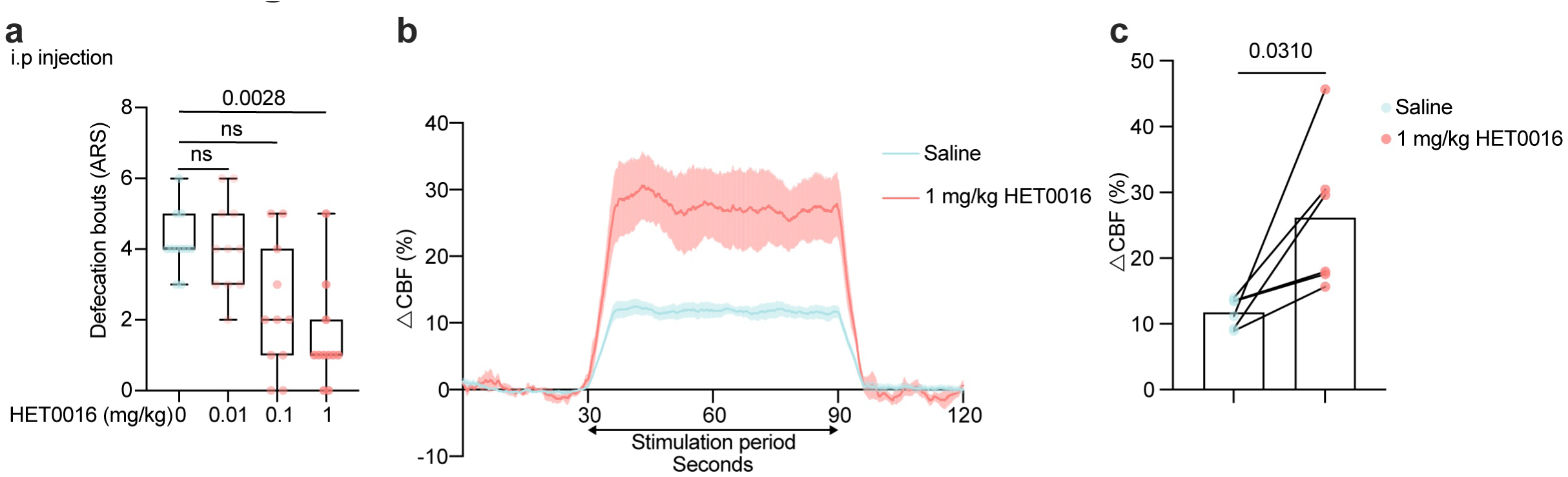
Dose-dependent and functional validation of HET0016. **a** Effects of different HET0016 concentrations on defecation bouts in ARS. Each dot represents one mouse. Kruskal-Wallis test followed by Dunn’s multiple comparisons test (*P* = 0.0005). **b, c** Functional hyperemia in the S1BF during whisker stimulation (**b**) and quantification of the corresponding plateau phase (**c**). Shaded areas represent SEM across mice. Each dot represents one mouse. Paired Student’s *t*-test. Data are presented as mean ± SEM.

**Fig. S11.**
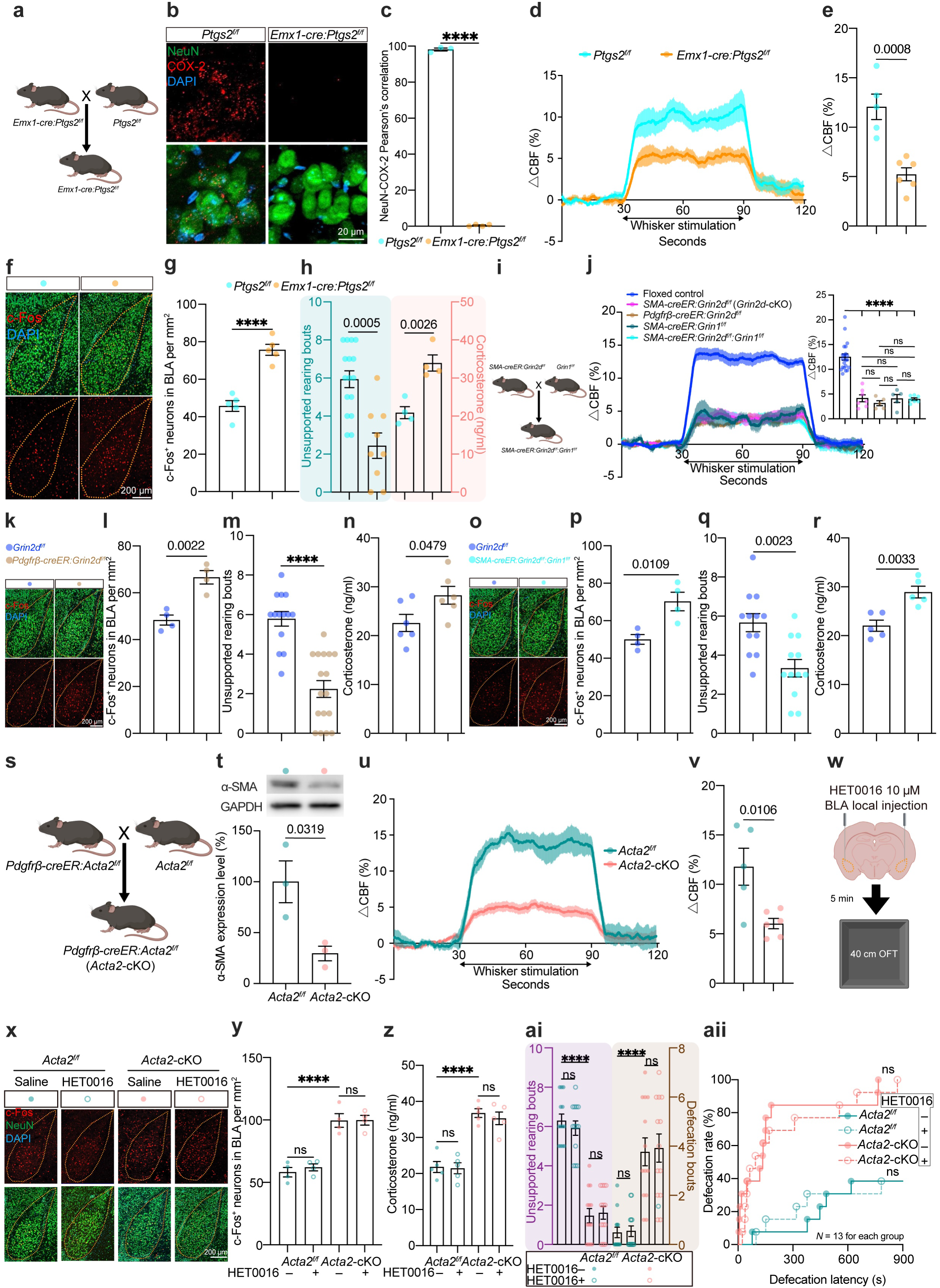
NVC dysfunction across multiple models drives negative emotionality. a–h. Validation of the *Emx1-cre:Ptgs2* model. Experimental schematic of mating strategy (**a**). Representative images (**b**) and quantification of neuronal COX-2 (encoded by *Ptgs2*) expression (**c**). Functional hyperemia in the S1BF (**d, e**), BLA neuronal activation (**f, g**), unsupported rearing bouts (**h, left**), and serum corticosterone (**h, right**) in the 40 cm OFT. **i–r** Validation of the mural cell-based NMDA conditional knock out models. Mating strategy (**i**), functional hyperemia (**j**), BLA neuronal activation (**k, l**, and **o, p**). Unsupported rearing (**m, q**) and serum corticosterone (**n, r**) in the 40 cm OFT. **s–v** Validation of the *Acta2*-cKO model. Mating strategy (**s**), α-SMA immunoblot quantification (**t**), and functional hyperemia in the S1BF (**u, v**). **w–aii** Effects of local BLA HET0016 injection in *Acta2*-cKO mice. Experimental design (**w**), BLA neuronal activation (**x, y**), serum corticosterone concentrations (**z**), unsupported rearing bouts (**ai, left**), and defecation bouts (**ai, right**), and defecation rate and latency in the 40 cm OFT (**aii**) with local BLA HET0016 injection. Each dot represents one mouse. Shaded areas represent SEM across mice in panel (**d**, **j**, and **u**). Floxed control mice group in (**j**) including comparable 8 *Grin2d^f/f^* mice, 7 *Grin1^f/f^* mice, and 3 *Grin2d^ff/^: Grin1^f/f^* mice. Control data (*Grin2d^f/f^*, *Grin1^f/f^*, *SMA-creER:Grin2d^f/f^*, and *SMA-creER:Grin1^f/f^*) are shared with Fig. 2g. Student’s *t-*test (**c**, **e**, **g**, **h right**, **l**, **n**, **p**, **r**, **t**, and **v**). Mann-Whitney test (**h left**, **m**, and **q**). Two-way ANOVA followed by Tukey’s multiple comparisons test for (**y,** *P* = 0.6564; **z**, *P* = 0.7144). Two-way ANOVA on rank-transformed data followed by Sidak’s multiple comparisons test (**ai left**, *P* = 0.4442; **ai right**, *P* = 0.7932). One-way ANOVA followed by Tukey’s multiple comparisons test (**j**, *P* < 0.0001). Gehan-Breslow-Wilcoxon test (**aii**). Sample sizes (*N*) are indicated in the panel (**aii**). **** indicates *P* < 0.0001. Data are presented as mean ± SEM.

**Fig. S12.**
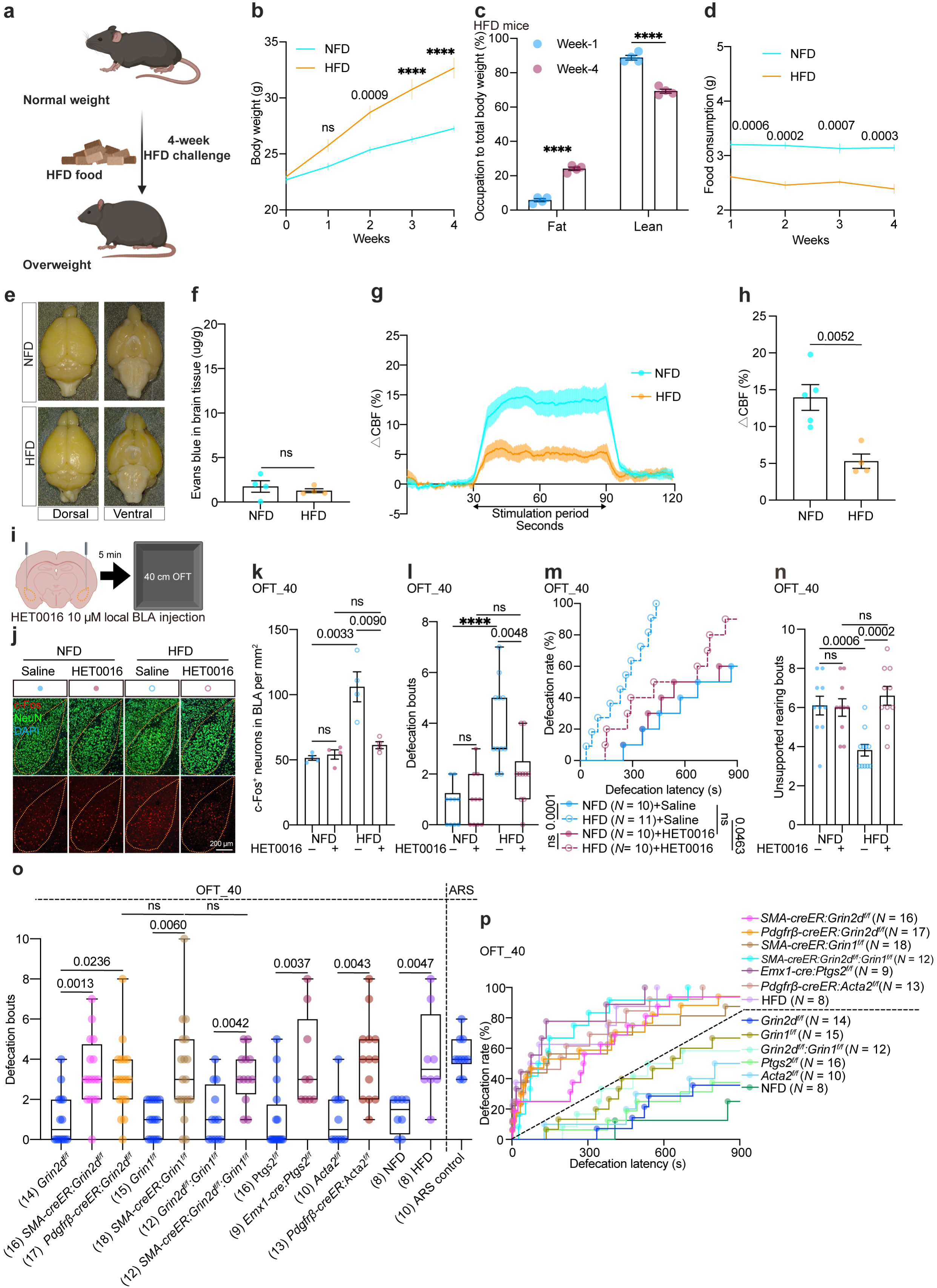
HFD-induced NVC dysfunction and its impact on emotional distress. a–d. High-fat diet (HFD) model validation. Schematic (**a**), body weight changes (**b**), body composition (fat and lean mass) changes (**c**), and food consumption (**d**) of mice with HFD or normal-fat diet (NFD). Student’s *t-*test. *N* = 16 mice per group for **b** and **d**, two-way ANOVA followed by Tukey’s multiple comparisons test (*P* < 0.0001). **e-f** Assessment of BBB integrity via Evans Blue injection (**e**) and quantification (**f**). student’s *t-*test. **g, h** Functional hyperemia in the S1BF (**g**) and quantification of the plateau phase (**h**). shaded areas represent SEM across mice in (**g**). Student’s *t-*test. **i–n** Effects of local BLA HET0016 injection in HFD mice. Experimental design (**i**), BLA neural activation (**j, k**), defecation bouts (**l**), defecation rate and latency (**m**), and unsupported rearing (**n**) of HFD mice in the 40 cm OFT with local BLA HET0016 injection. Two-way ANOVA followed by Tukey’s multiple comparisons test for (**k**), *P* = 0.0001. Two-way ANOVA on rank-transformed data followed by Sidak’s multiple comparisons test (**l** and **n**), *P* < 0.0001 for (**l**), and *P* = 0.0009 for (**n**). Gehan-Breslow-Wilcoxon test (**m**). Sample sizes (*N*) are indicated in the panel (**m**). **o, p** Defecation bouts (**o**), and defecation rate and latency (**p**) in the 40 cm OFT and ARS (panel **o** only). Kruskal-Wallis test followed by Dunn’s multiple comparisons test for (**o**) (P < 0.0001). Gehan-Breslow-Wilcoxon test for (**p**). Sample sizes (*N*) are indicated in the panels (**o** and **p**). Control data (*Grin2d^f/f^*and *SMA-creER:Grin2d^f/ff^*) are shared with Fig. 3a**-c**. Detailed statistical data for panel (**p**) are shown in the supplementary table S1, sub-table 6. Dotted line in (**p**) used to separate two cluster of distribution pattern between NVC-deficiency groups and their littermate controls. Each dot represents one mouse. **** indicates *P* < 0.0001. Data are presented as mean ± SEM.

**Fig. S13.**
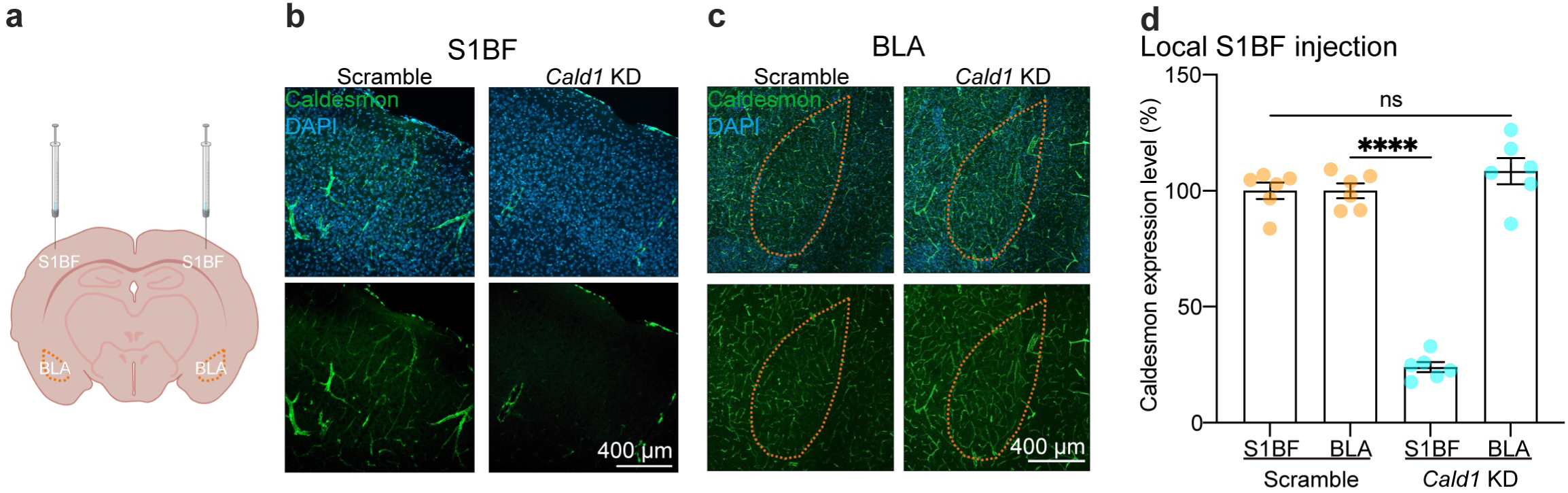
Specificity of S1BF *Cald1* knockdown. **a** Schematic showing the siRNA injection site in the S1BF and its spatial relationship with the BLA (orange dotted line). **b–d** Representative images of caldesmon IF stainningin the S1BF (**b**) and BLA (**c**) following local S1BF *Cald1* siRNA injection, with quantification in (**d**). Each dot represents one mouse. Two-way ANOVA followed by Tukey’s multiple comparisons test (*P* < 0.0001). **** indicates *P* < 0.0001. Data are presented as mean ± SEM.

**Fig. S14.**
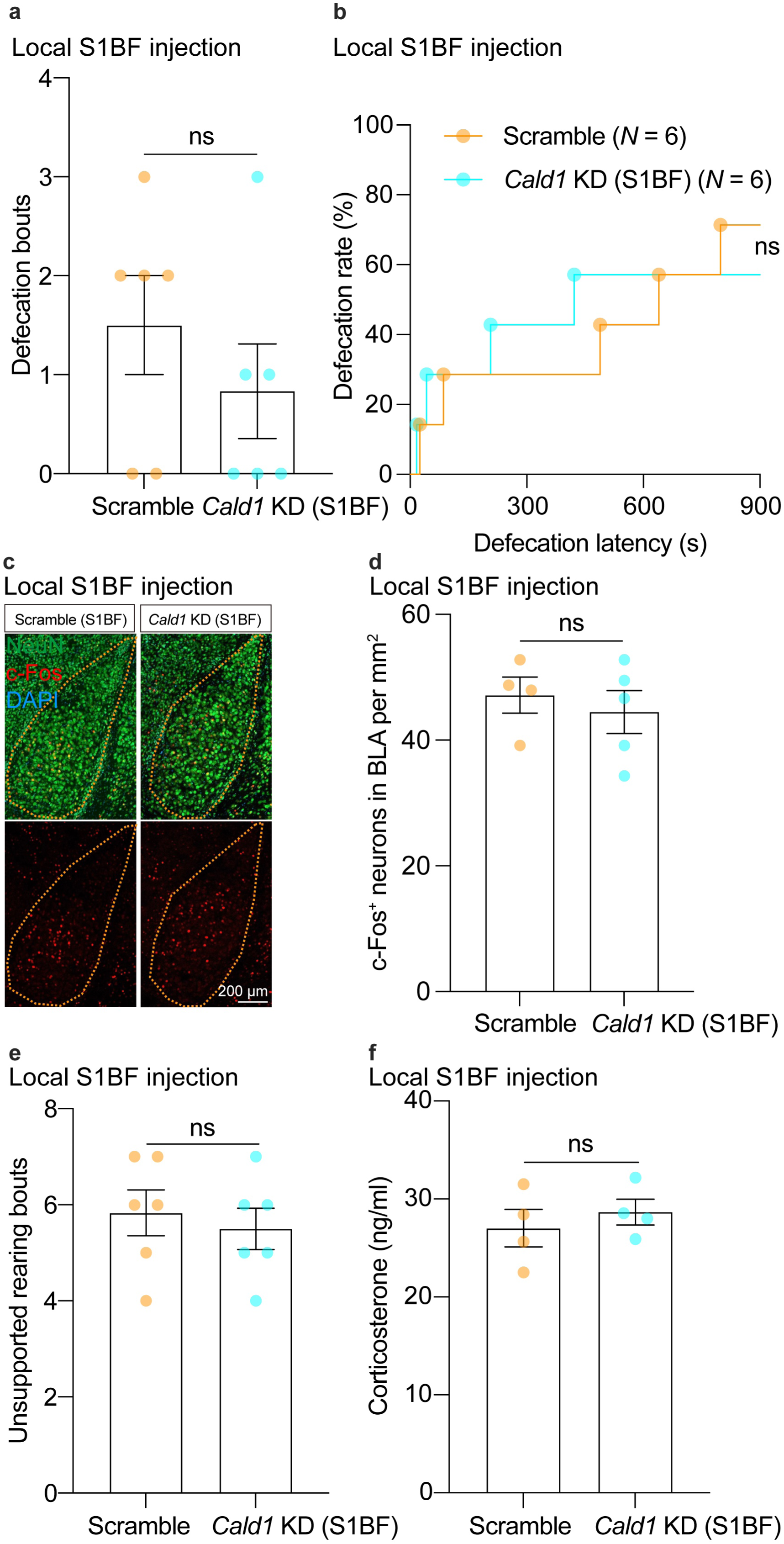
Impact of S1BF *Cald1* knockdown on emotional responses. a, **b** Defecation bouts (**a**), and defecation rate and latency (**b**) in the 40 cm OFT following S1BF *Cald1* KD in the 40 cm OFT. Each dot represents one mouse. Mann-Whitney test in (**a**). Sample sizes (*N*) are indicated in the panel (**b**). Gehan-Breslow-Wilcoxon test in (**b**). **c, d** Representative images (**c**) and quantification (**d**) of BLA neuronal activation in the 40 cm OFT. Each dot represents one mouse. Student’s *t-*test. **e, f** Unsupported rearing bouts (**e**) and serum corticosterone concentrations (**f**) in the 40 cm OFT. Each dot represents one mouse. Mann-Whitney test in (**e**), and student’s *t*-test in (**f**). Data are presented as mean ± SEM.

**Fig. S15.**
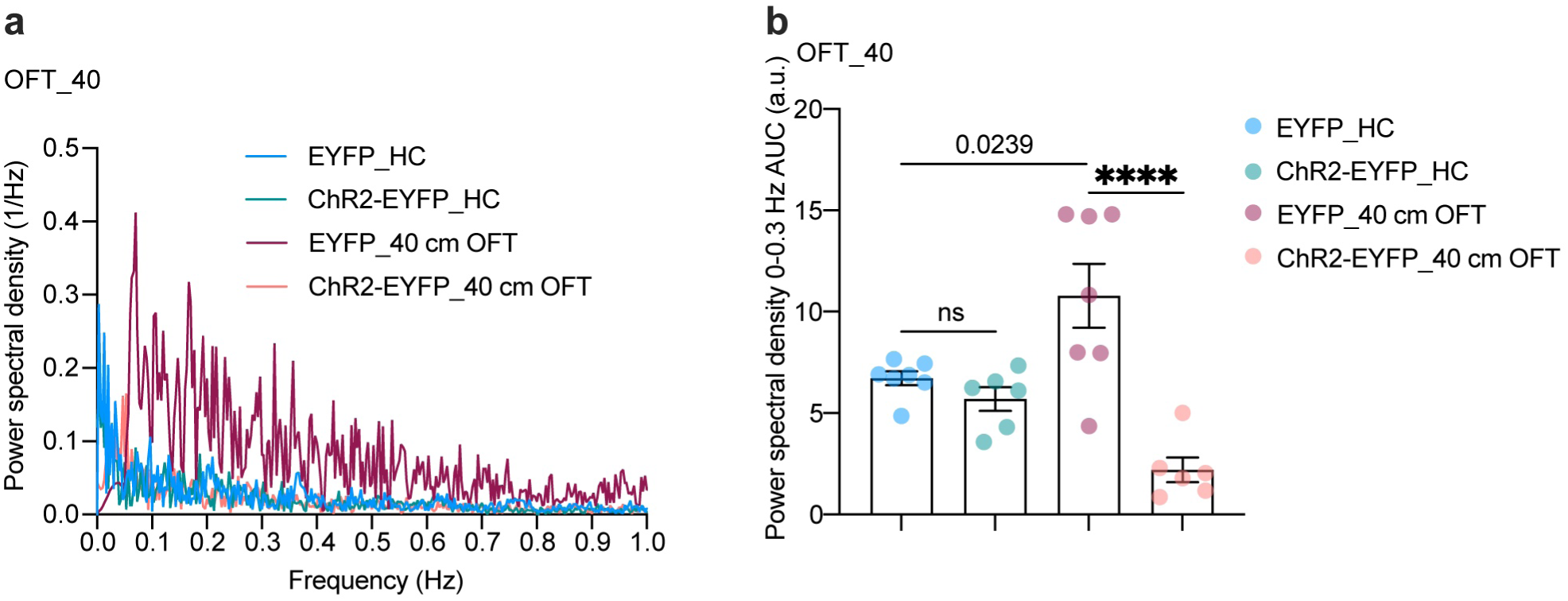
Spectral energy analysis of BLA blood flow signals. a, **b** Representative Fourier plot of BLA CBV signals from 0–1 Hz (**a**) and quantification of ultra-low-frequency oscillations (0–0.3 Hz) (**b**) in the 40 cm OFT. Each dot represents one mouse. One-way ANOVA followed by Tukey’s multiple comparisons test (*P* < 0.0001). **** indicates *P* < 0.0001. Data are presented as mean ± SEM.

**Fig. S16.**
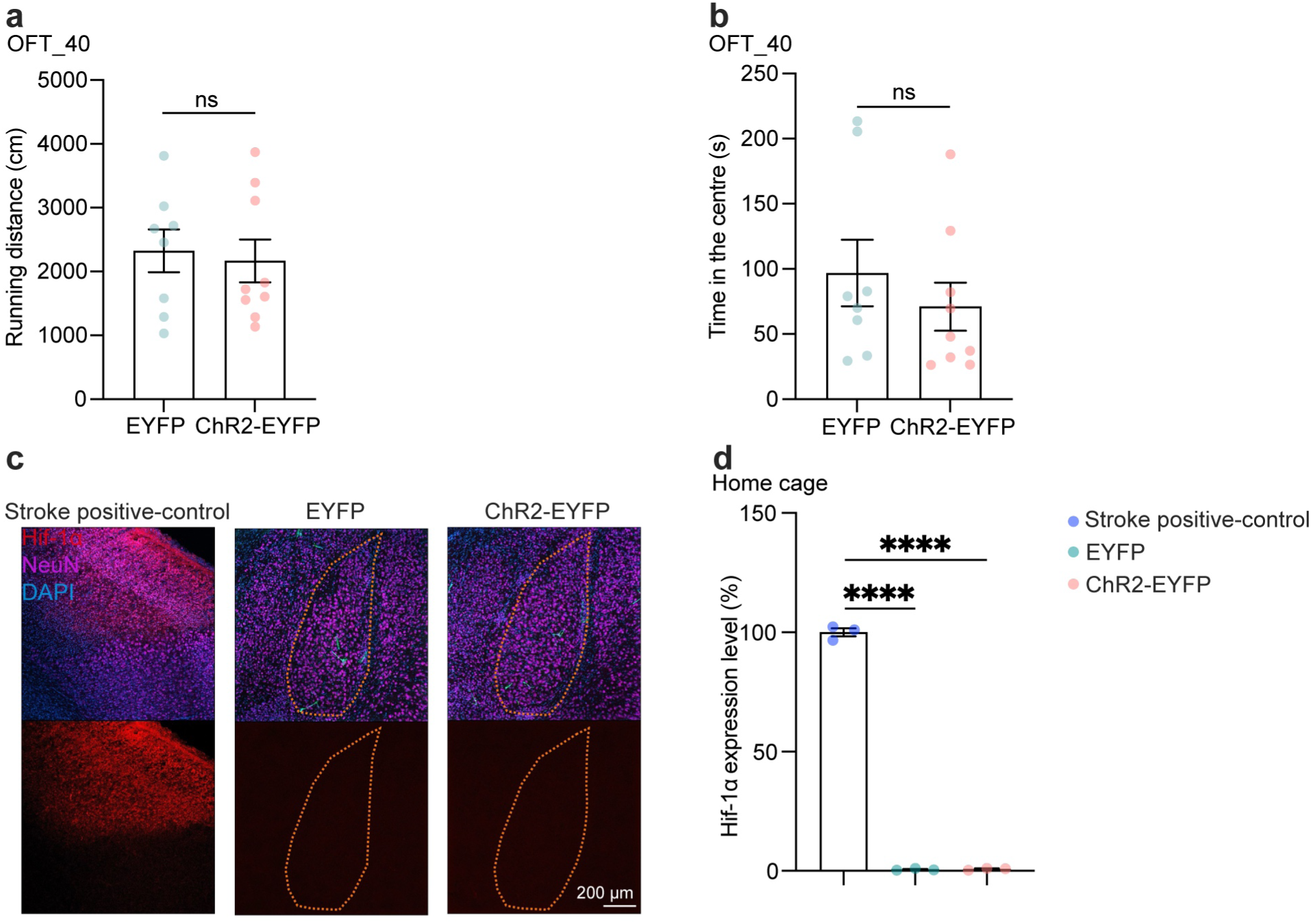
Supplementary data for BLA optogenetic stimulation. a, **b** Running distance (**a**) and time spent in the center zone (**b**) during optogenetic stimulation in the 40 cm OFT. Each dot represents one mouse. Student’s *t*-test. **c, d** Representative images (**c**) and quantification (**d**) of Hif-1α immunofluorescence in the BLA during the optogenetic stimulation in the home cage. Stroke model tissue serves as a positive control. Each dot represents one mouse. One-way ANOVA followed by Tukey’s multiple comparisons test (*P* < 0.0001). **** indicates *P* < 0.0001. Data are presented as mean ± SEM.

**Fig. S17.**
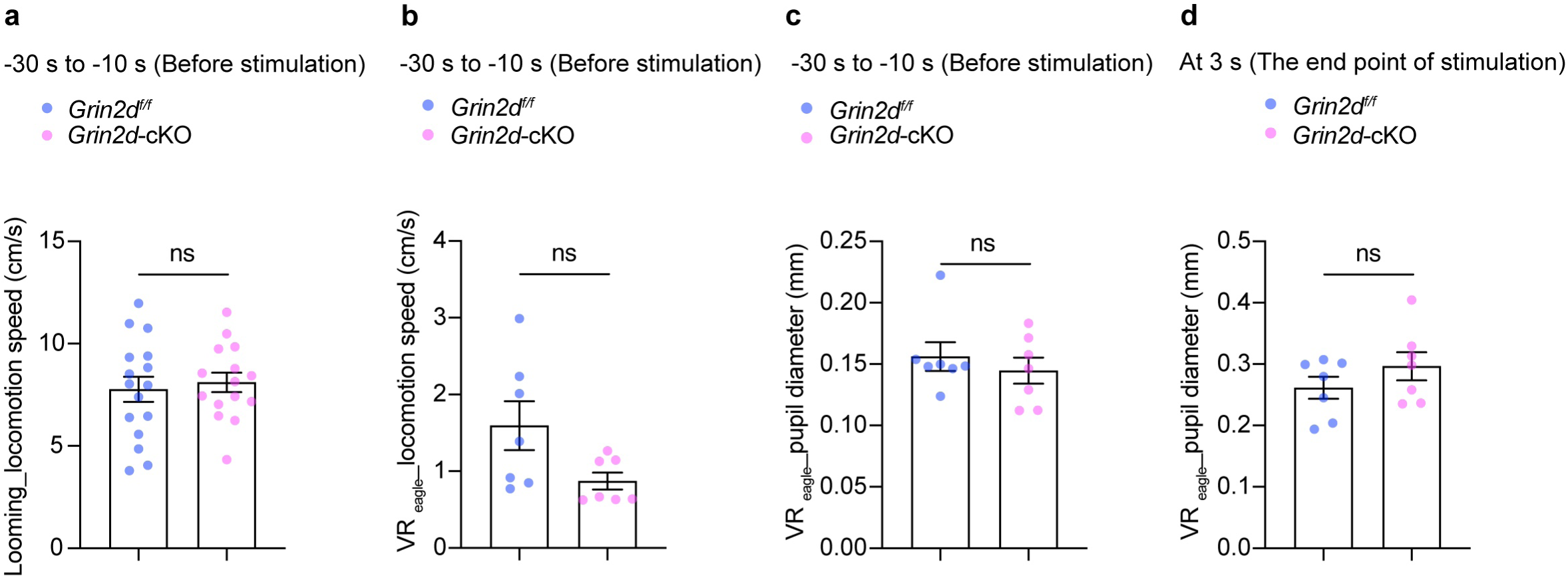
Supplementary locomotion speed and pupil diameter in looming and VR_eagle_ assays. a–d. Quantification of pre-stimulation locomotion speed in looming (**a**) and VR_eagle_ (**b**) assays, and pupil diameter before (**c**) and at the conclusion (**d**) of VR_eagle_ stimulation. Each dot represents one mouse. Student’s *t*-test. Data are presented as mean ± SEM.

**Fig. S18.**
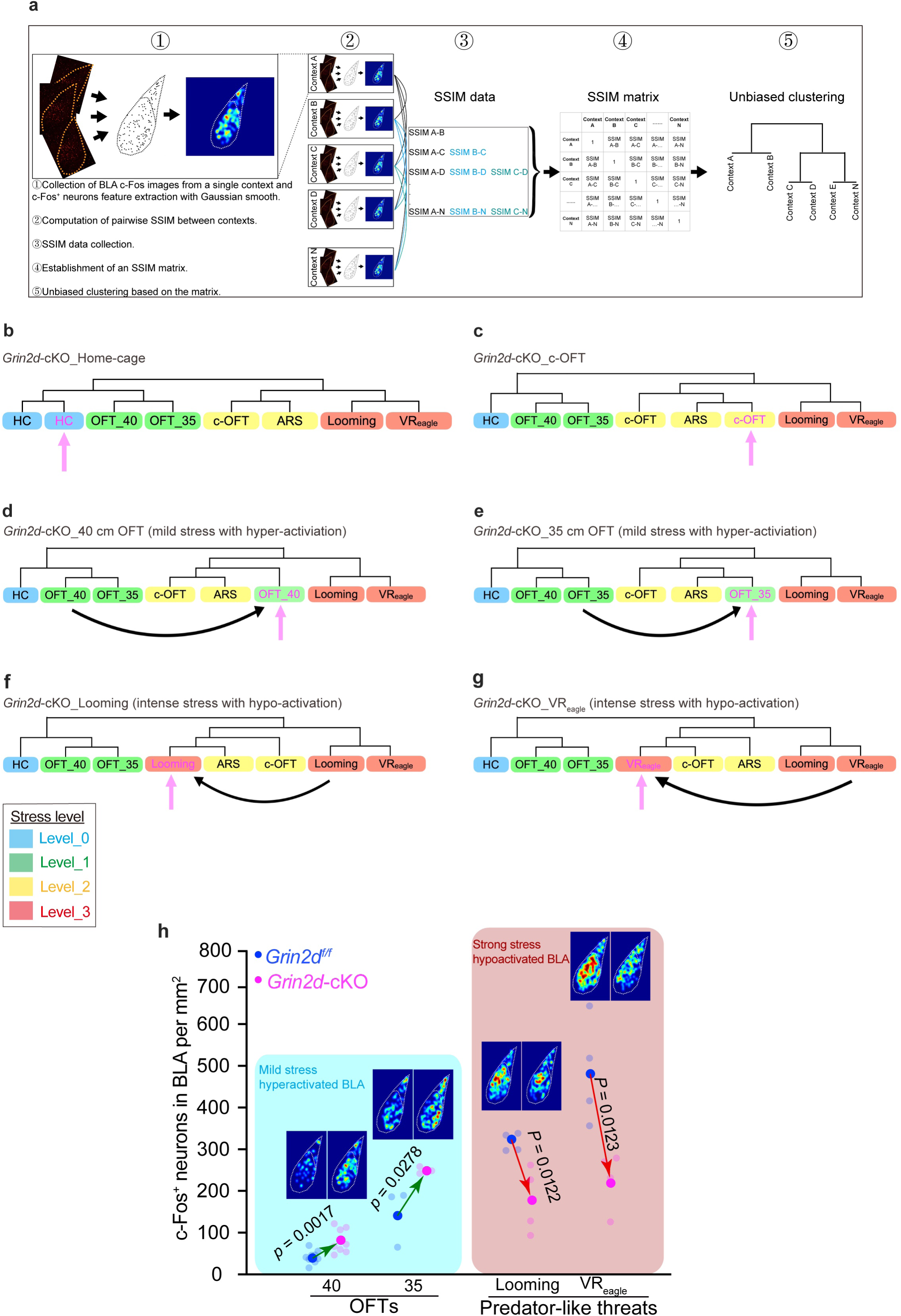
Methodological framework for topological clustering and clustering coordinates of *Grin2d*-cKO mice. **a** Schematic workflow for clustering data processing and generation. **b–g**, Clustering coordinates of *Grin2d*-cKO mice mapped against the unbiased classification library of littermate controls across individual contexts. **h** the summary of the load-dependent biphasic pattern. Quantification data shared with Fig. 6j, and heatmap shared with **Figure. 6l, m**. Each 50% transparent dot represents one mouse. The non-transparent dot represents the mean value. Student’s *t*-test. Data are presented as mean ± SEM.

**Fig. S19.**
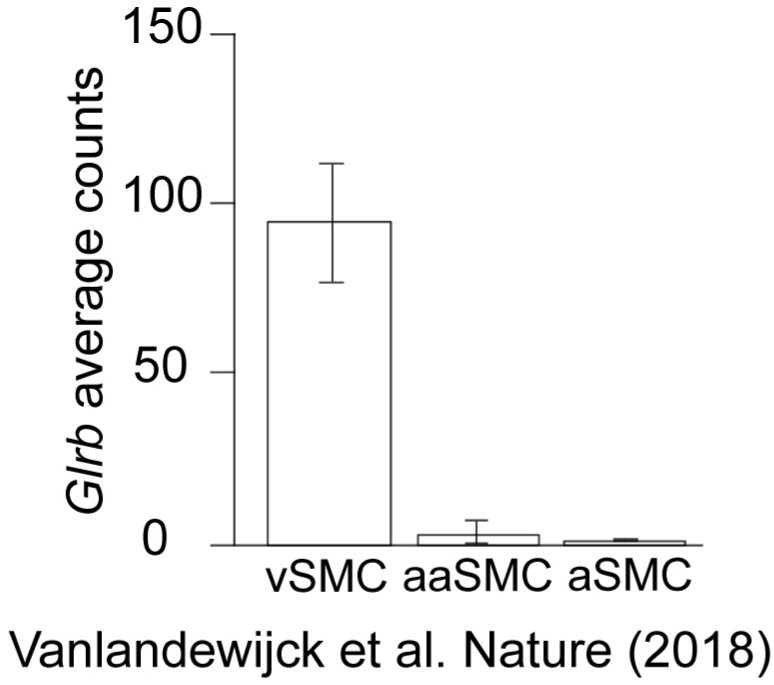
Glrb expression profile in the vascular transcriptome database. Transcriptional levels of *Glrb* in venous SMCs (vSMCs), arteriolar SMCs (aaSMCs), and arterial SMCs (aSMCs). Data are presented as mean ± SEM.

**Fig. S20.**
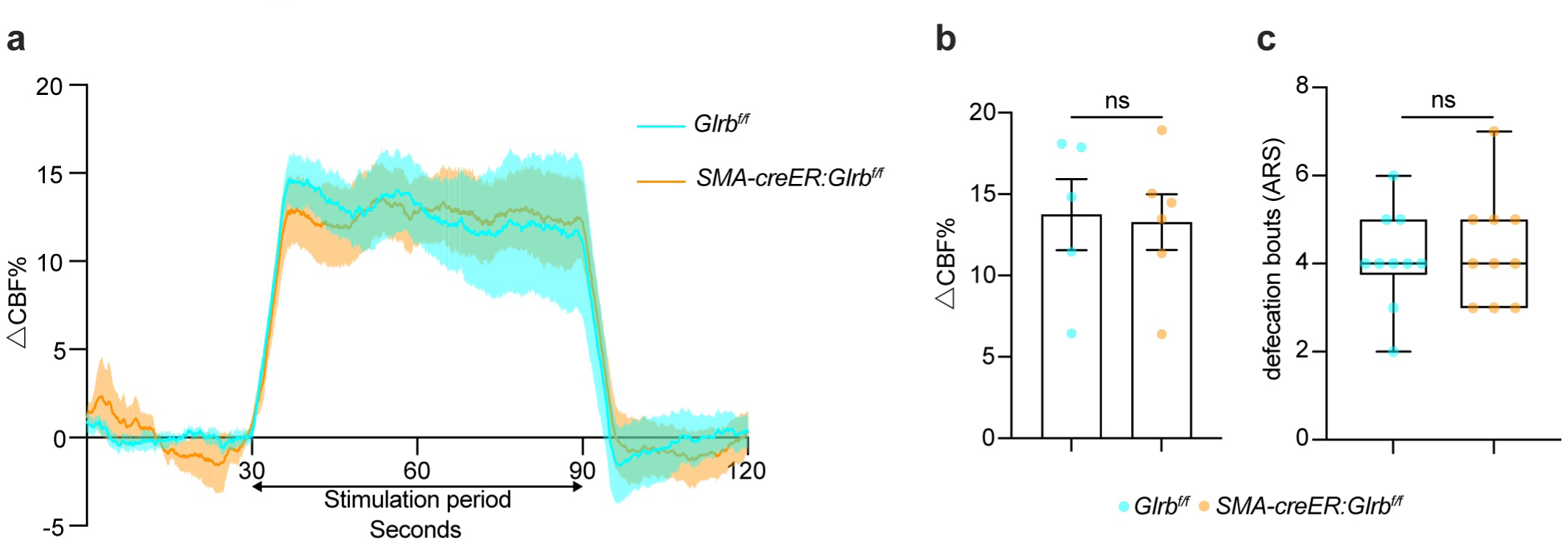
Functional and behavioral validation of SMC-specific *Glrb*-cKO mice. a, **b** Functional hyperemia in the S1BF (**a**) and quantification of the plateau phase (**b**). **c** Defecation bouts of mice during the ARS. Each dot represents one mouse. Shaded areas represent SEM across mice in (**a**). Student’s *t*-test (**b**) and Mann-Whitney test (**c**). Data are presented as mean ± SEM.

**Fig. S21.**
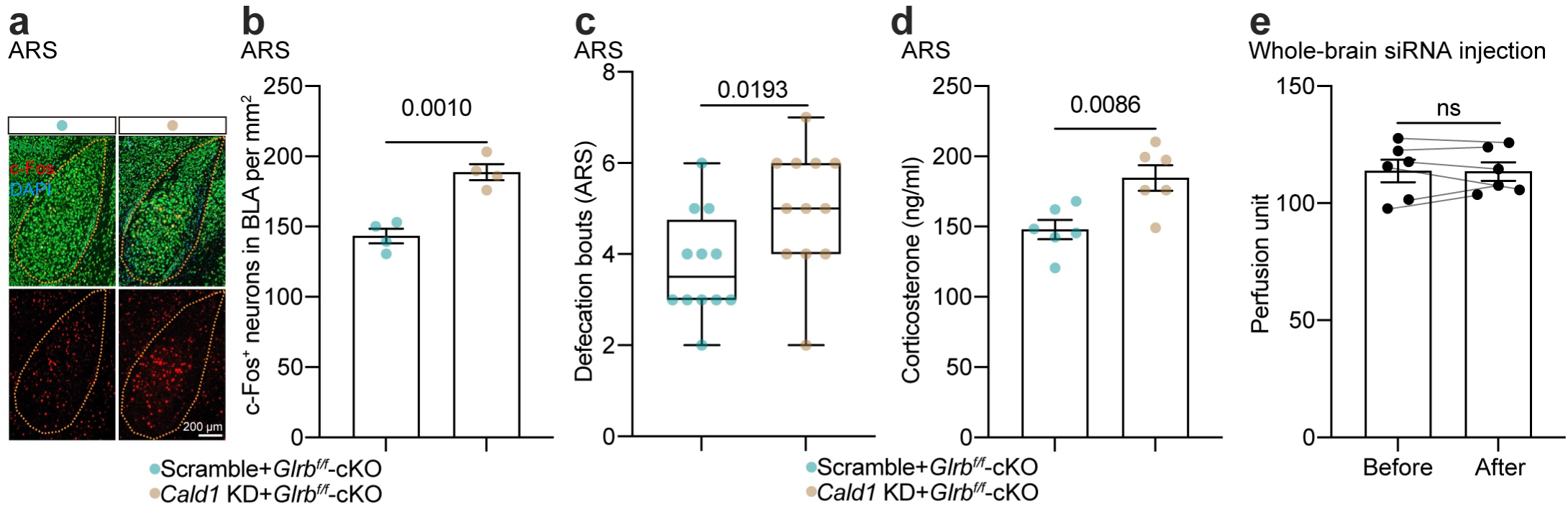
Local BLA *Cald1* knockdown occludes the NVC enhancement in *Glrb*-cKO mice. a, **b** Representative images (**a**) and quantification (**b**) of BLA neuronal activation of *Glrb*-cKO mice with local BLA *Cald1* KD in ARS. Student’s *t*-test. **c, d** Defecation bouts (**c**) and serum corticosterone (**d**) in *Glrb*-cKO mice with local BLA *Cald1* KD in ARS. Mann-Whitney test (**c**) and Student’s *t*-test (**d**). **e** Baseline cerebral perfusion analysis before and after *Cald1* siRNA cisterna magna injection in the *Glrb*-cKO mice. Paired Student’s *t*-test. Each dot represents one mouse. Data are presented as mean ± SEM.

**Fig. S22.**
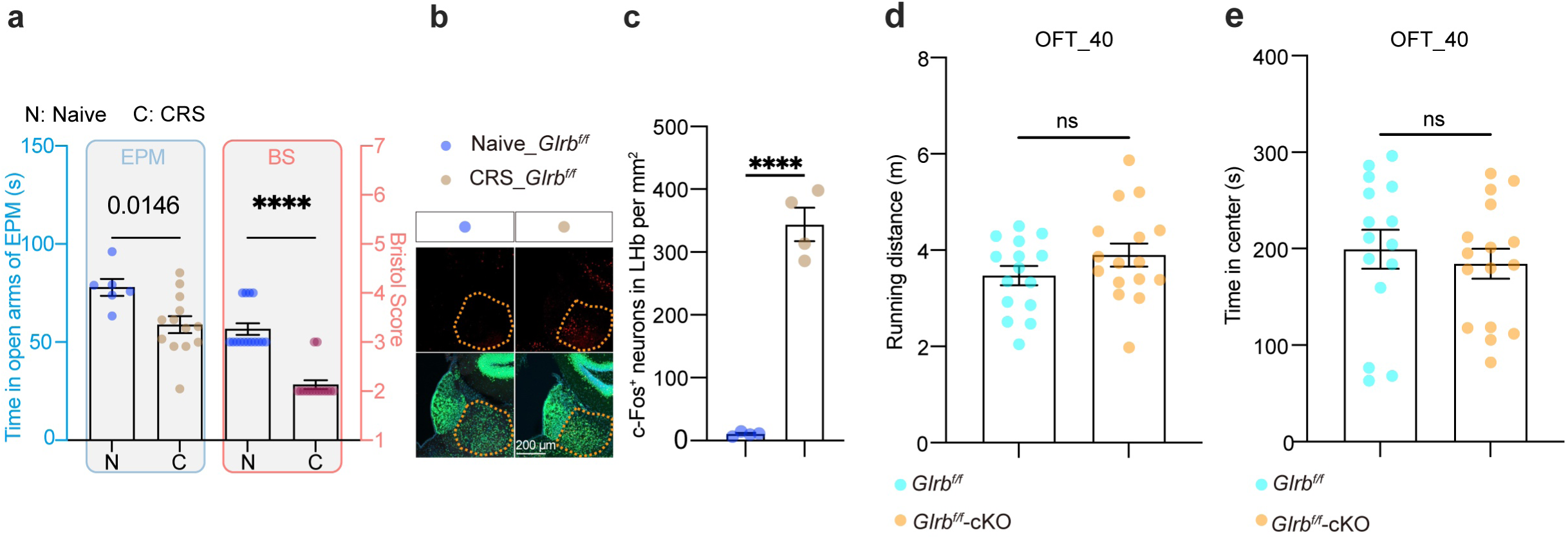
CRS modelling and *Glrb*-cKO mice naïve behaviours in the 40 cm OFT. **a** Time spent in the open arms of the elevated plus maze after CRS modelling (EPM, left) and fecal quality (Bristol score, right, data similar to that in Fig. 8m) at day-28 of CRS. Student’s *t*-test (left) and Mann-Whitney test (right). **b, c** Representative images (**b**) and quantification (**c**) of neuronal activation in the lateral habenula (LHb) after CRS modelling. Student’s *t*-test. **d, e** Total running distance (**d**) and time spent in the center zone (**e**) in the 40 cm OFT. Naïve indicates mice without CRS modelling as controls. Each dot represents one mouse. Student’s *t*-test. **** indicates *P* < 0.0001. Data are presented as mean ± SEM.

**Fig. S23.**
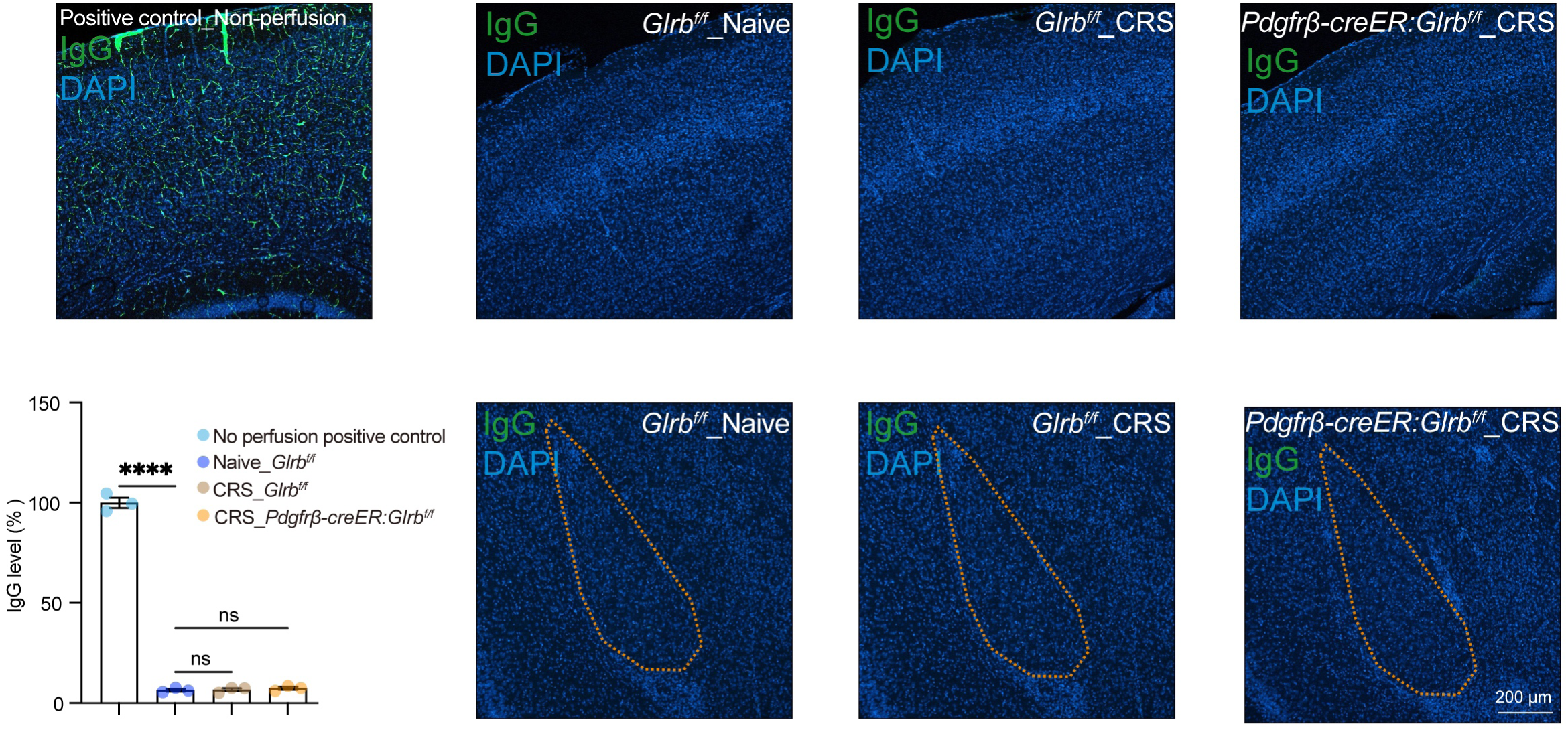
Assessment of BBB integrity following CRS. Representative images (including non-perfusion positive control) and quantification of IgG fluorescence in the S1BF and BLA after CRS modelling. Each dot represents the average IgG level per mouse, calculated from three ROIs in both the cortex and the BLA. One-way ANOVA followed by Tukey’s multiple comparisons test (*P* < 0.0001). **** indicates *P* < 0.0001. Data are presented as mean ± SEM.

**Fig. S24.**
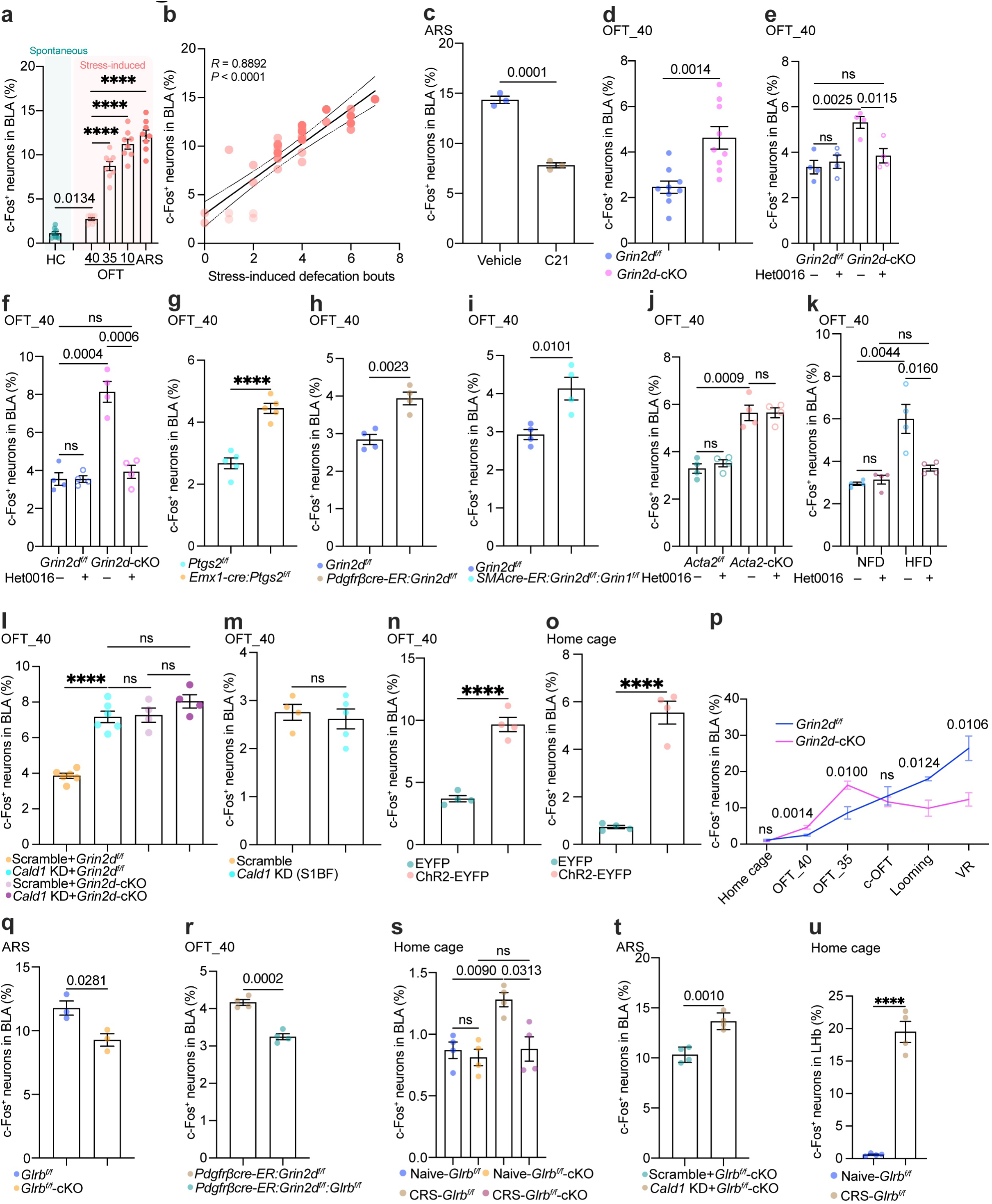
Statistical quantification of the percentage of c-Fos positive neurons in the BLA. (a–u) Re-quantification of c-Fos positive neuron ratios (normalized to total NeuN positive neurons in the BLA) corresponding to the original intensity-based data presented in Figure 1c (**a**), Figure 1e (**b**), Figure 1o (**c**), Figure 2m (**d**), Figure 3g (**e**), Figure 3m (**f**), Figure S11g (**g**), Figure S11l (**h**), Figure S11p (**i**), Figure S11y (**j**), Figure S12k (**k**), Figure 4m (**l**), Figure S14d (**m**), Figure 5i (**n**), Figure 5r left (**o**), Figure 6j (**p**), Figure 8b (**q**), Figure 8h (**r**), Figure 8p left (**s**), Figure S21b (**t**), and Figure S22c (**u**), respectively.

## Supplementary tables

**Supplementary Table S1.**
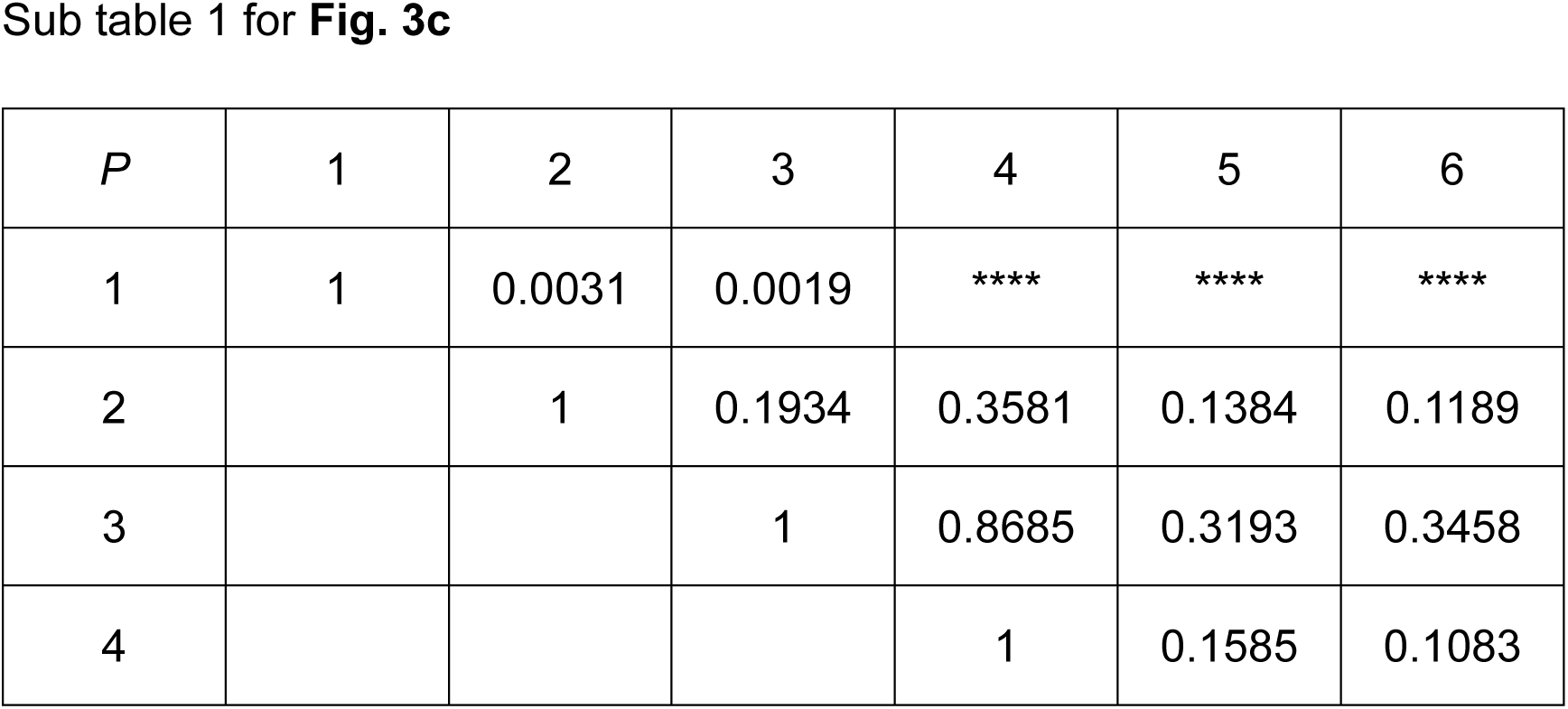
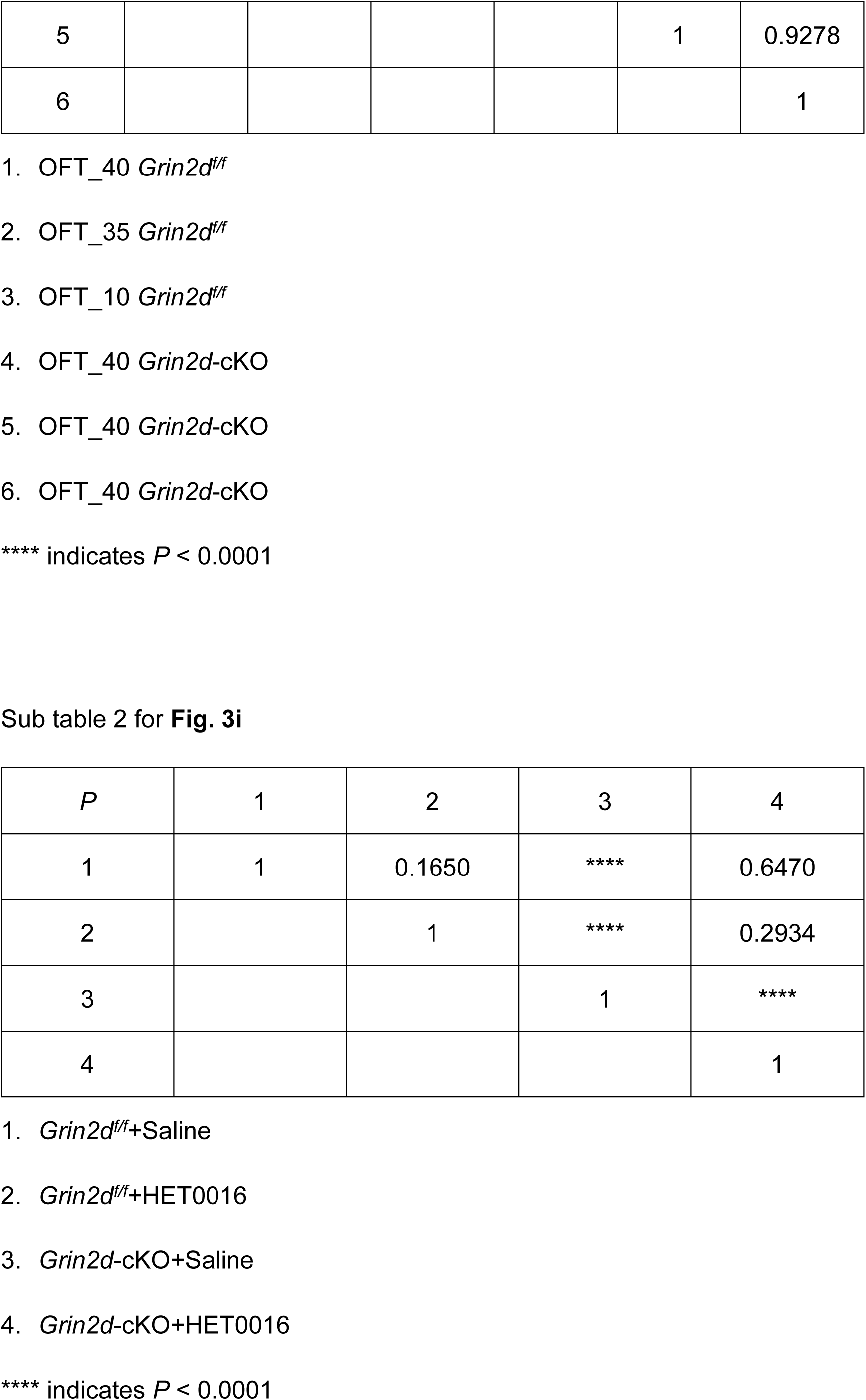
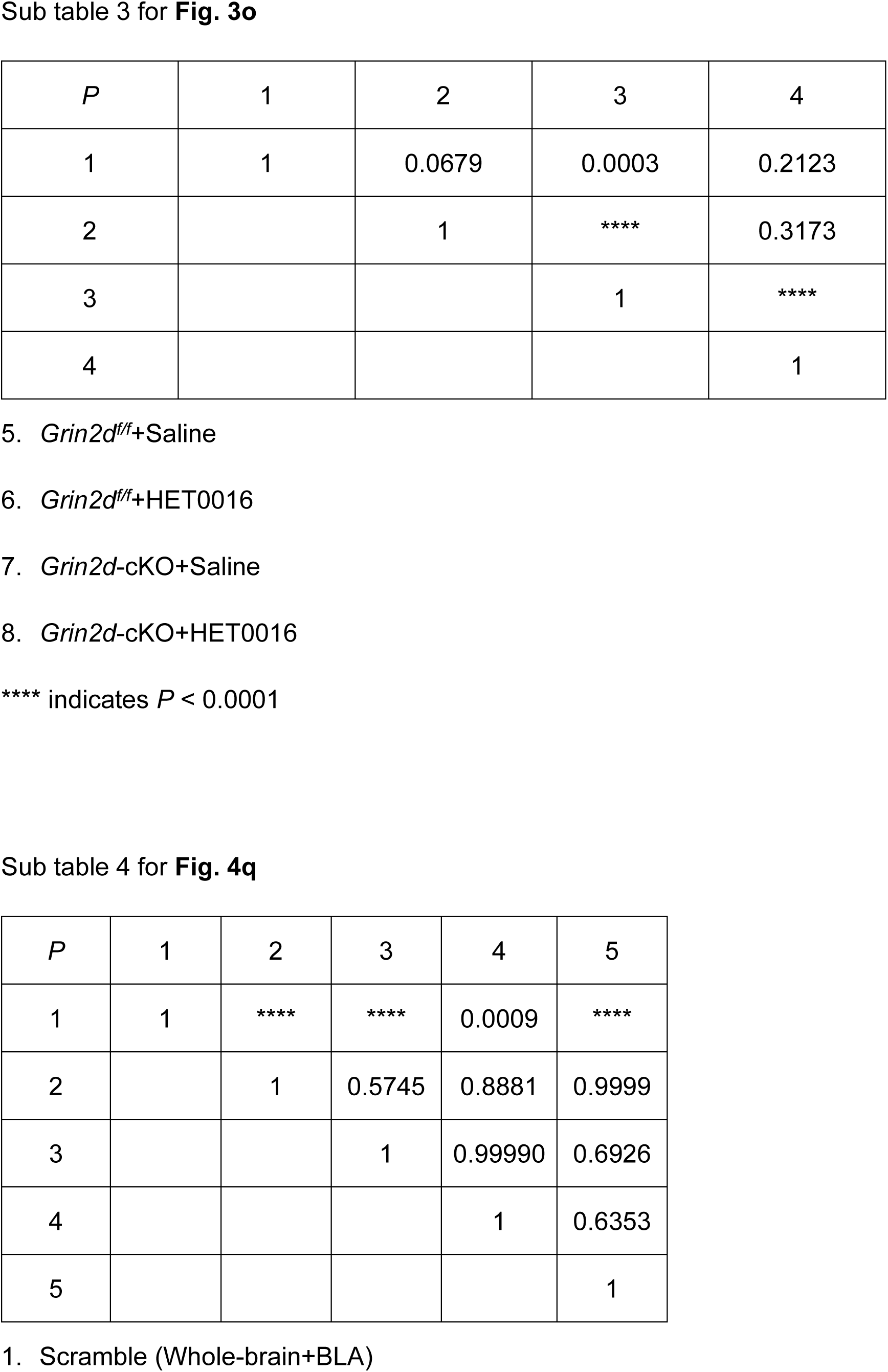
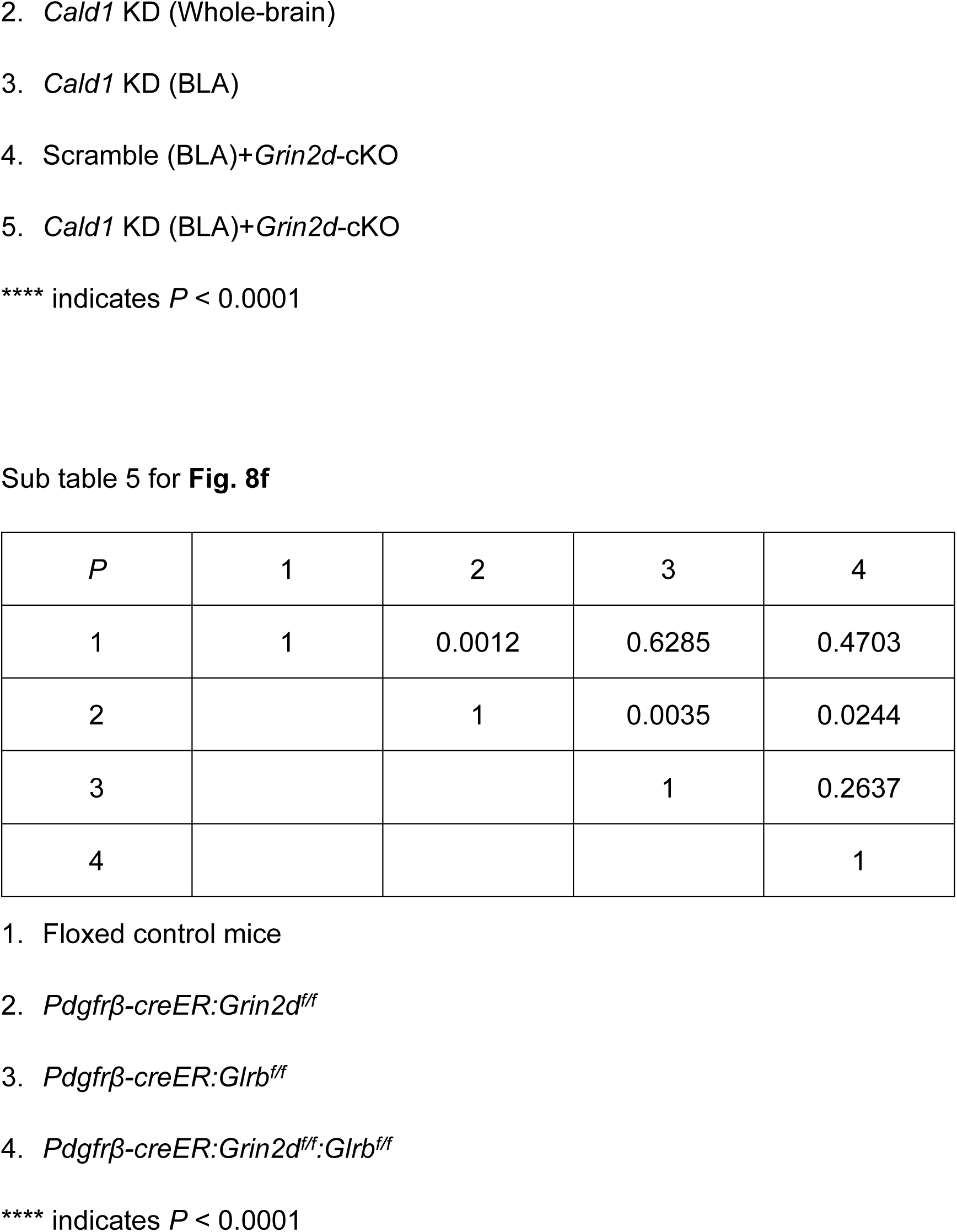
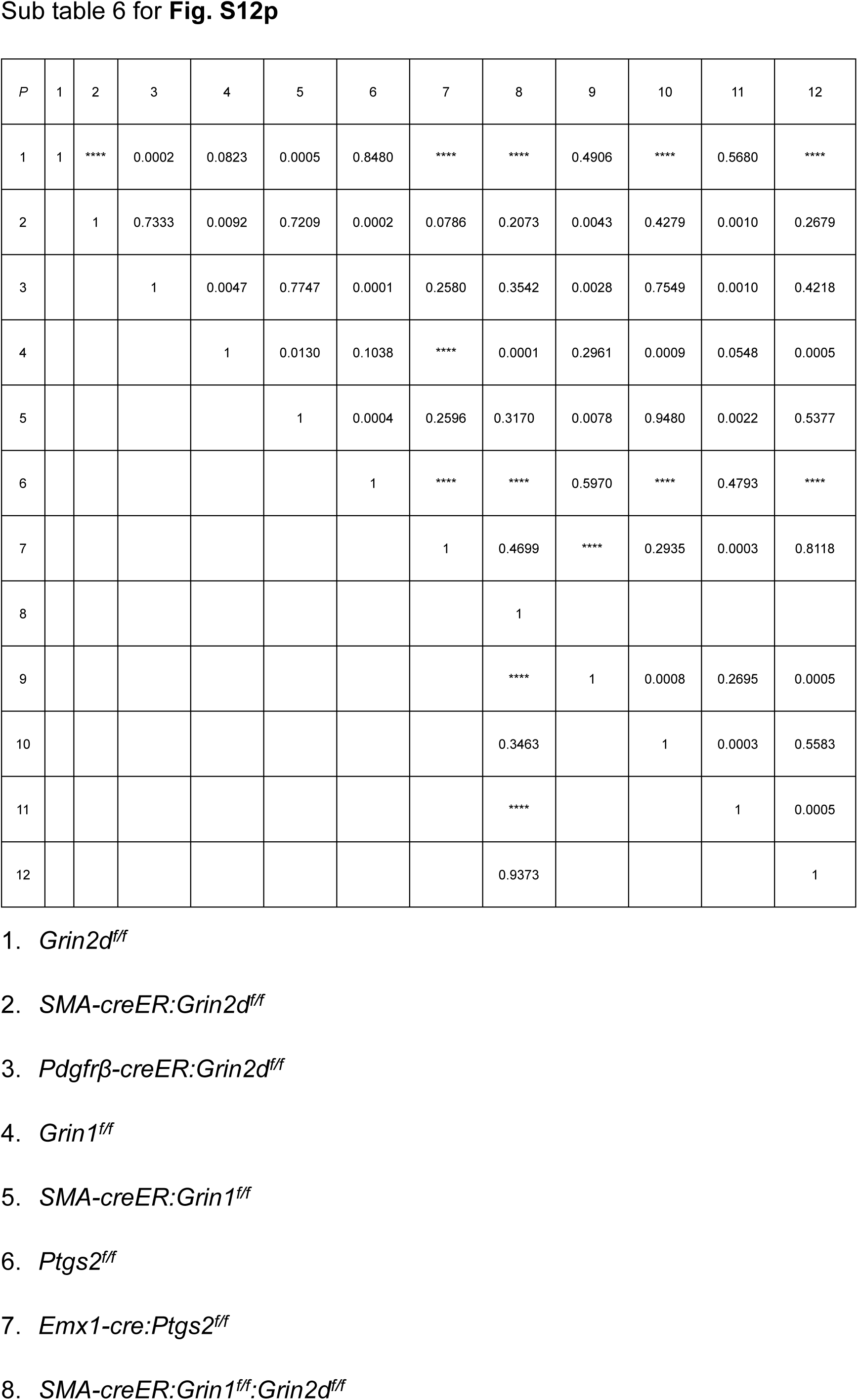
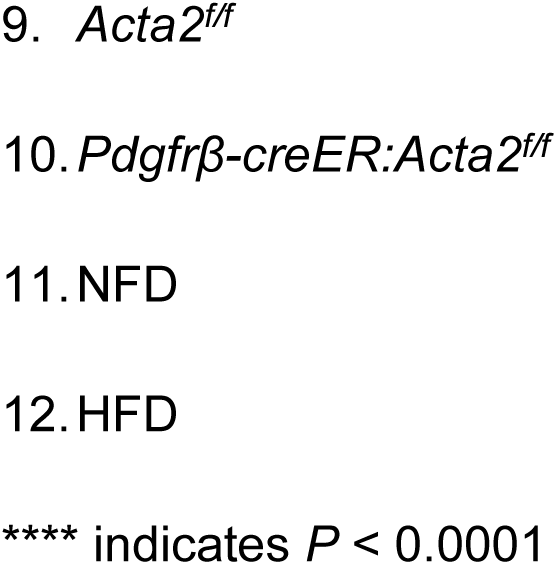
*P_value_* of Defecation rate-latency curve Sub table 1 for Fig. 3c.

**Supplementary Table S2.** Sample source checklist.

| No. | Location | Sample type | Source |
| --- | --- | --- | --- |
| 1 | Fig. 1b | Immunofluorescence staining | Brain slice |
| 2 | Fig. 1n | Immunofluorescence staining | Brain slice |
| 3 | Fig. 2b | Semi-quantified PCR | Sorted cultured cell |
| 4 | Fig. 2c | RNAscope | Brain slice |
| 5 | Fig. 2d | Immunofluorescence staining | Isolated MCA |
| 6 | Fig. 2l | Immunofluorescence staining | Brain slice |
| 7 | Fig. S1a | RNAscope | Brain slice |
| 8 | Fig. S1c | Western blot | Cerebellum lysates |
| 9 | Fig. S1d | Western blot | Sorted cultured cell |
| 10 | Fig. S1e | Western blot | Cortex lysates |
| 11 | Fig. S2a | Immunofluorescence staining | Tissue slice |
| 12 | Fig. S3i | Hematoxylin and eosin staining | Intestine slice |
| 13 | Fig. S3o | Absorbance measurement | Brain lysates |
| 14 | Fig. S4a | Western blot | Sorted culture cell |
| 15 | Fig. S4b | Immunofluorescence staining | Isolated MCA |
| 16 | Fig. 3f | Immunofluorescence staining | Brain slice |
| 17 | Fig. 3l | Immunofluorescence staining | Brain slice |
| 18 | Fig. S11b | Immunofluorescence staining | Brain slice |
| 19 | Fig. S11f | Immunofluorescence staining | Brain slice |
| 20 | Fig. S11k | Immunofluorescence staining | Brain slice |
| 21 | Fig. S11o | Immunofluorescence staining | Brain slice |
| 22 | Fig. S11t | Western blot | Cortex lysates |
| 23 | Fig. S11x | Immunofluorescence staining | Brain slice |
| 24 | Fig. S12e | Absorbance measurement | Brain lysates |
| 25 | Fig. S12j | Immunofluorescence staining | Brain slice |
| 26 | Fig. 4b | Western blot | Sorted culture cell |
| 27 | Fig. 4d | Immunofluorescence staining | Brain slice |
| 28 | Fig. 4f | Semi-quantified PCR | Cortex lysate |
| 29 | Fig. 4g | Immunofluorescence staining | Brain slice |
| 30 | Fig. 4h | Immunofluorescence staining | Brain slice |
| 31 | Fig. 4l | Immunofluorescence staining | Brain slice |
| 32 | Fig. S13b | Immunofluorescence staining | Brain slice |
| 33 | Fig. S13c | Immunofluorescence staining | Brain slice |
| 34 | Fig. S14c | Immunofluorescence staining | Brain slice |
| 35 | Fig. 5a | Immunofluorescence staining | Brain slice |
| 36 | Fig. 5h | Immunofluorescence staining | Brain slice |
| 37 | Fig. 5q | Immunofluorescence staining | Brain slice |
| 38 | Fig. S16c | Immunofluorescence staining | Brain slice |
| 39 | Fig. 7a | RNAscope | Brain slice |
| 40 | Fig. 7c | Immunofluorescence staining | Brain slice |
| 41 | Fig. 7f | Semi-quantified PCR | Acute sorted cell |
| 42 | Fig. 7g | Immunofluorescence staining | Cultured cell line |
| 43 | Fig. 7h | Immunofluorescence staining | Brain slice |
| 44 | Fig. 7i | Electron microscopy image | Brain slice |
| 45 | Fig. 8a | Immunofluorescence staining | Brain slice |
| 46 | Fig. 8g | Immunofluorescence staining | Brain slice |
| 47 | Fig. 8o | Immunofluorescence staining | Brain slice |
| 48 | Fig. S21a | Immunofluorescence staining | Brain slice |
| 49 | Fig. S22b | Immunofluorescence staining | Brain slice |
| 50 | Fig. S23 | Immunofluorescence staining | Brain slice |

**Supplementary Table S3.**
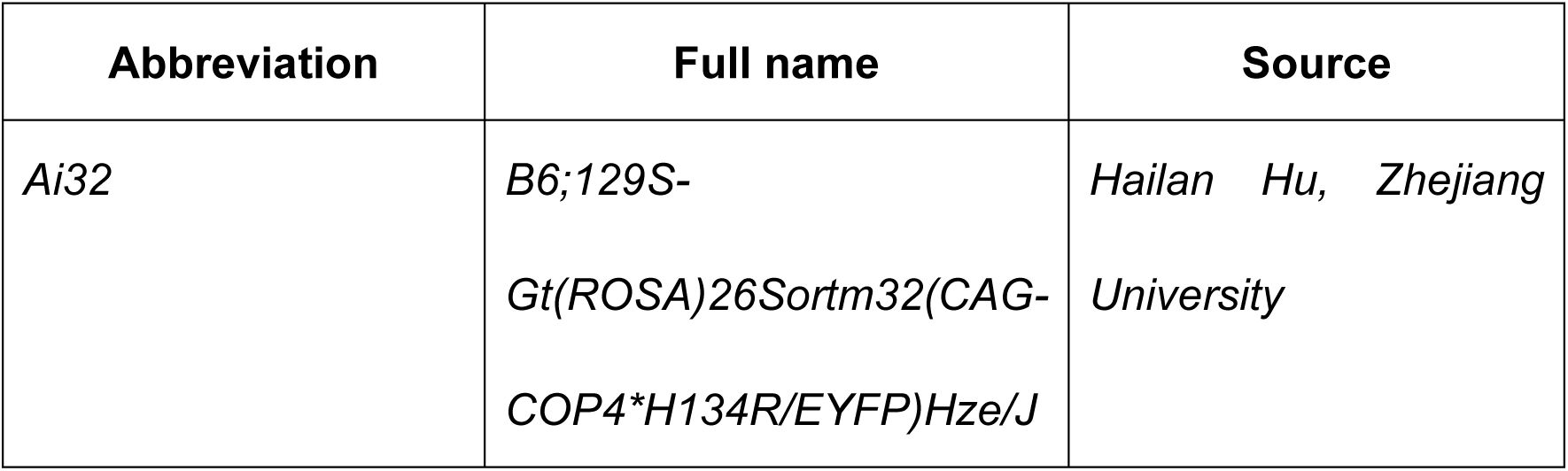
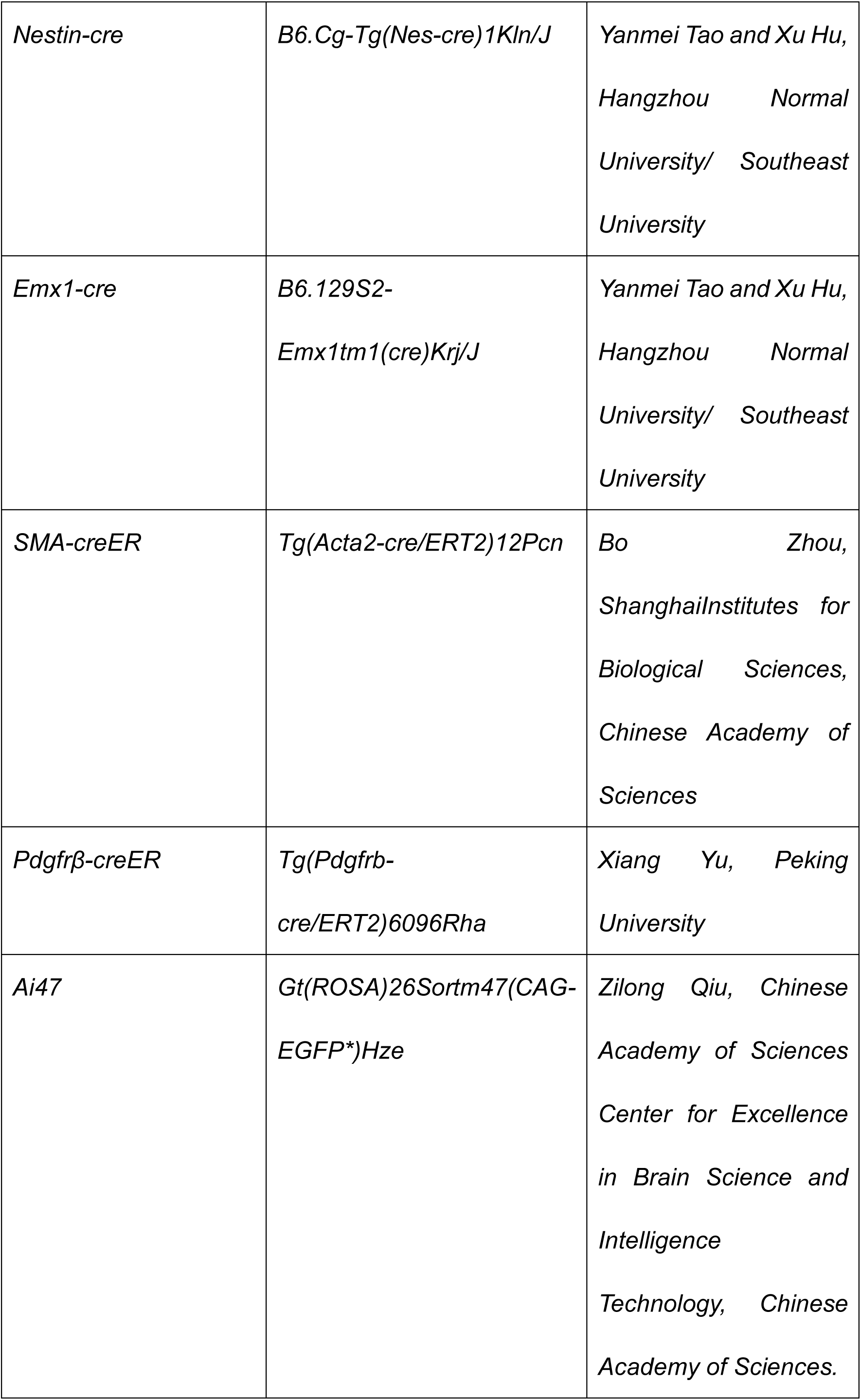
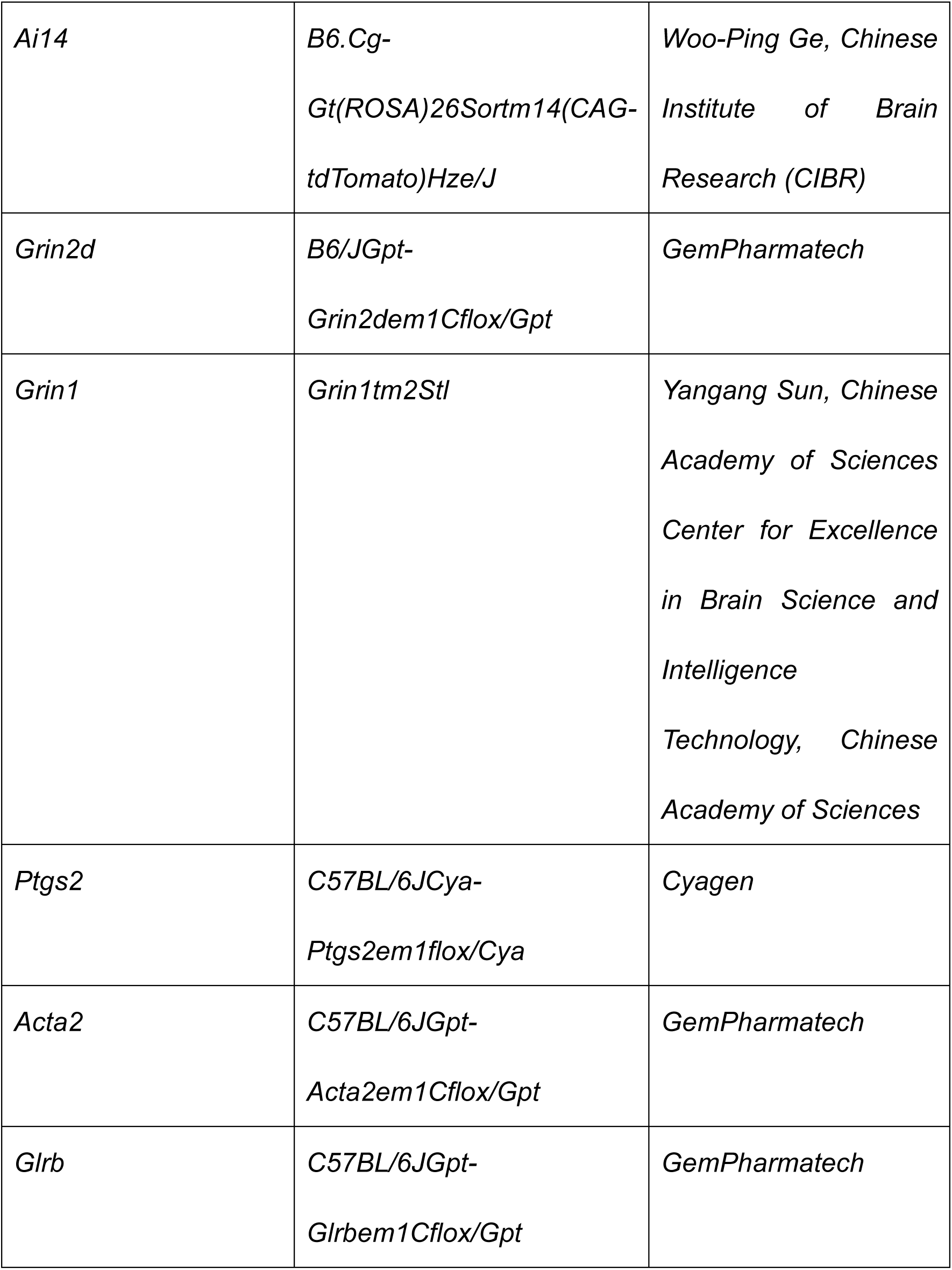
Mice information.

**Supplementary Table S4.** Antibodies information.

| **Name** | **Host** | **Cat#** | **Dilution** | **Company** |
| --- | --- | --- | --- | --- |
| Anti-NMDAR2D Antibody | Mouse | MAB5578 | 1:400 (IF)<br>1:2,500<br>(WB)<br>1:1,000 (IP) | Merck |
| Anti-Smoothelin antibody | Rabbit | ab219652 | 1:400 (IF) | abcam |
| α-Smooth Muscle Actin (ACTA2)<br>Rabbit mAb | Rabbit | A17910 | 1:2,500<br>(WB) | ABclonal |
| ACE2 Recombinant Rabbit<br>Monoclonal Antibody (SN0754) | Rabbit | MA5-32307 | 1:400 (IF) | Invitrogen |
| Gephyrin Polyclonal Antibody | Rabbit | PA5-29036 | 1:400 (IF) | Invitrogen |
| BD Pharmingen™ Purified Rat Anti-Mouse CD31 | Rat | 557355 | 1:200 (IF) | BD |
| Anti-NeuN purified Antibody | Guinea<br>Pig | ABN90P | 1:800 (IF) | Merck |
| COX2/PTGS2 Rabbit pAb | Rabbit | A1253 | 1:400 (IF) | ABclonal |
| c-Fos antibody | Rabbit | 226 003 | 1:1,000 (IF) | SYSY |
| Recombinant Alexa Fluor® 488 Anti-Caldesmon/CDM antibody | Rabbit | ab208116 | 1:400 (IF)<br>1:2,500 (WB) | abcam |
| iFluor™ 488 Conjugated Goat anti-mouse IgG polyclonal Antibody | Goat | HA1125 | 1:400 (IF) | HUABIO |
| Mouse Aminopeptidase N/CD13 Antibody | Goat | AF2335 | 1:200 (IF) | R&D |
| CD140b (PDGFRB) Monoclonal Antibody (APB5), APC, eBioscience™ | Rat | 17-1402-82 | 1:50 (FCM) | Invitrogen |
| Mice IgG | Mouse | A7028 | 1:1,000 (IP) | Beyotime |
| Donkey anti-Mouse IgG (H+L) ReadyProbes™ Secondary Antibody, Alexa Fluor™ 488 | Donkey | R37114 | 1:1,000 (IF) | Invitrogen |
| Donkey anti-Rabbit IgG (H+L) ReadyProbes™ Secondary Antibody, Alexa Fluor™ 488 | Donkey | R37118 | 1:1,000 (IF) | Invitrogen |
| Donkey anti-Rabbit IgG (H+L) Highly Cross-Adsorbed Secondary Antibody, Alexa Fluor™ 546 | Donkey | A10040 | 1:1,000 (IF) | Invitrogen |
| Donkey anti-Rat IgG (H+L) Highly<br>Cross-Adsorbed Secondary<br>Antibody, Alexa Fluor™ 488 | Donkey | A21208 | 1:1,000 (IF) | Invitrogen |
| Donkey anti-Rat IgG (H+L) Highly<br>Cross-Adsorbed Secondary<br>Antibody, Alexa Fluor™ 555 | Donkey | A78945 | 1:1,000 (IF) | Invitrogen |
| Donkey anti-Rat IgG (H+L) Highly<br>Cross-Adsorbed Secondary<br>Antibody, Alexa Fluor™ Plus 647 | Donkey | A48272 | 1:1,000 (IF) | Invitrogen |
| Alexa Fluor® 647 AffiniPure™<br>Donkey Anti-Guinea Pig IgG (H+L) | Donkey | 706-605-148 | 1:1,000 (IF) | Jackson<br>ImmunoR<br>esearch |
| β-Tubulin Mouse mAb | Mouse | AC021 | 1:5,000<br>(WB) | ABclonal |
| Goat Anti-Mouse IgG, HRP<br>Conjugated | Goat | CW0102 | 1:10,000<br>(WB) | CWBIO |
| Goat Anti-Rabbit IgG, HRP<br>Conjugated | Goat | CW0103 | 1:10,000<br>(WB) | CWBIO |
| DDDDK-Tag Rabbit mAb | Rabbit | AE063 | 1:400 (IF) | ABclonal |
| Hif-1α | Rabbit | ER1802-41 | 1:100 (IF) | HUABIO |

**Supplementary Table S5.**
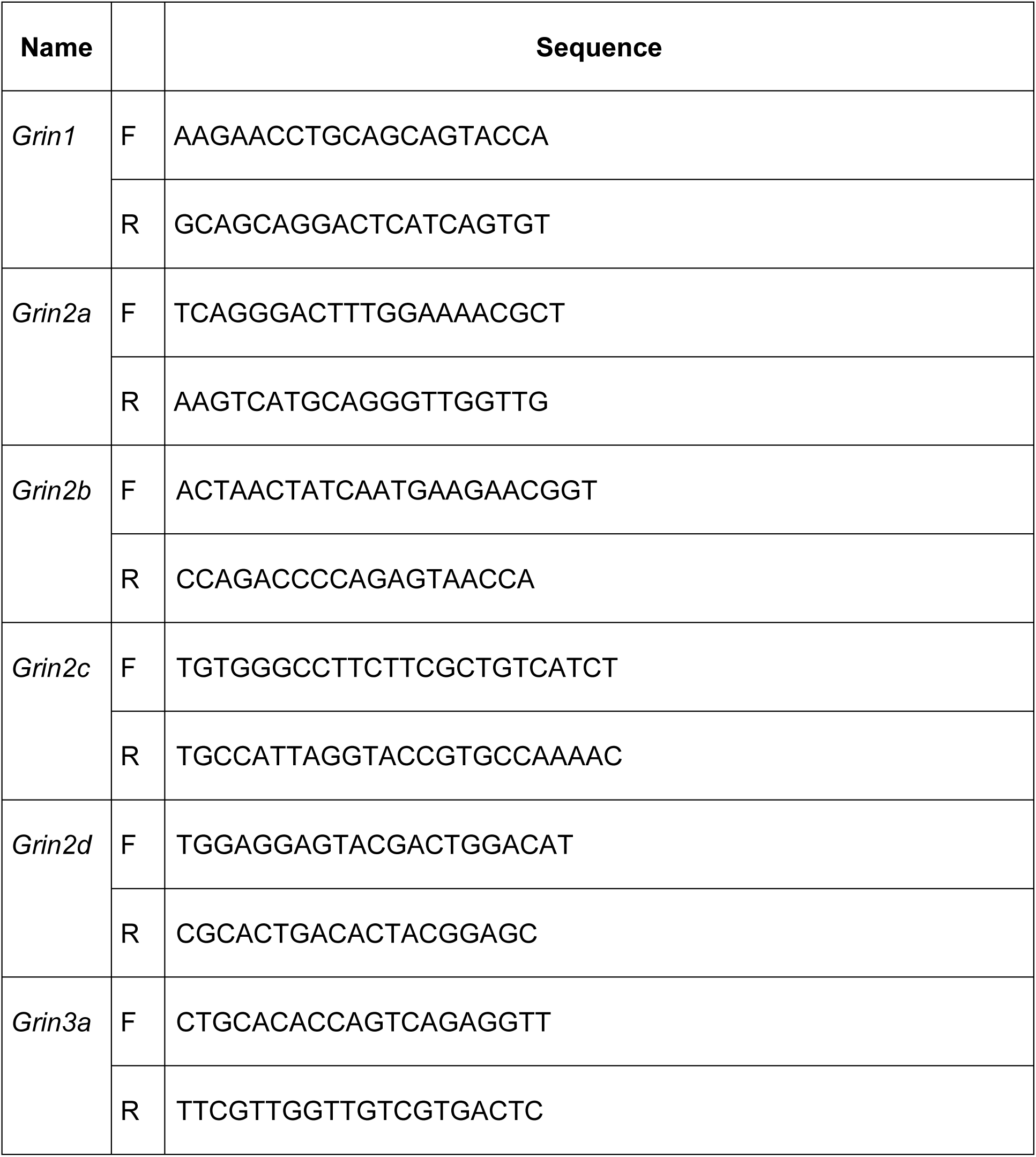
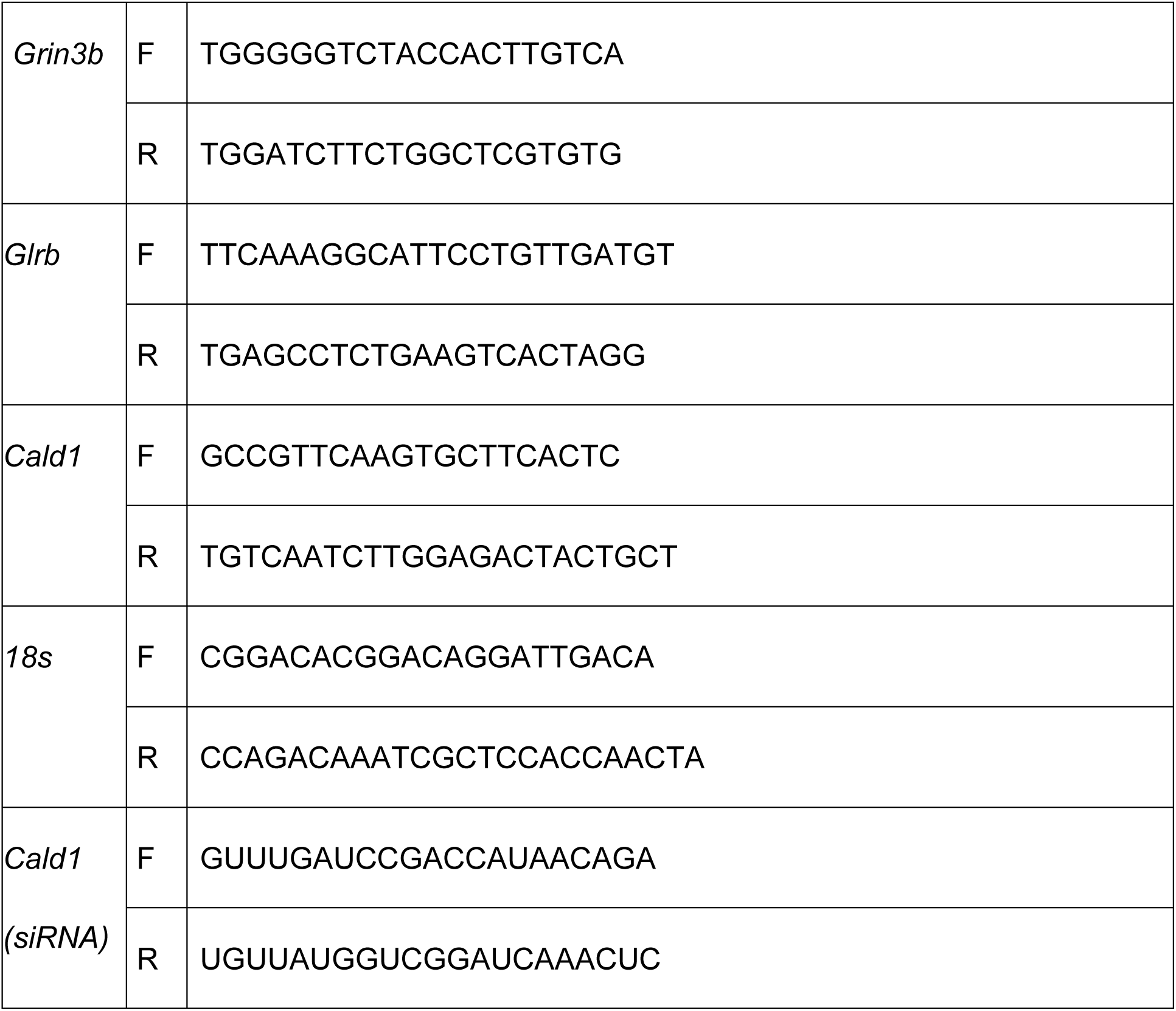
Primer pairs information.

## Supplementary Video

**Supplementary Video S1. Topological distribution of arterioles in the BLA.**

**Supplementary Video S2. Pupil diameter changes during the stimulation in the VR_eagle_ stimulation assay.**

